# Single-nucleus transcriptomic analysis reveals the regulatory circuitry of myofiber XBP1 during regenerative myogenesis

**DOI:** 10.1101/2024.06.03.597179

**Authors:** Aniket S. Joshi, Micah B. Castillo, Meiricris Tomaz da Silva, Anh Tuan Vuong, Preethi H. Gunaratne, Radbod Darabi, Yu Liu, Ashok Kumar

## Abstract

Endoplasmic reticulum (ER) stress-induced unfolded protein response (UPR) are activated in skeletal muscle in multiple conditions. However, the role of the UPR in the regulation of muscle regeneration remains less understood. We demonstrate that gene expression of various markers of the UPR is induced in both myogenic and non-myogenic cells in regenerating muscle. Genetic ablation of XBP1, a downstream target of the IRE1α arm of the UPR, in myofibers attenuates muscle regeneration in adult mice. Single nucleus RNA sequencing (snRNA-seq) analysis showed that deletion of XBP1 in myofibers perturbs proteolytic systems and mitochondrial function in myogenic cells. Trajectory analysis of snRNA-seq dataset showed that XBP1 regulates the abundance of satellite cells and the formation of new myofibers in regenerating muscle. In addition, ablation of XBP1 disrupts the composition of non-myogenic cells in injured muscle microenvironment. Collectively, our study suggests that myofiber XBP1 regulates muscle regeneration through both cell-autonomous and -non-autonomous mechanisms.

**HIGHLIGHTS:**

- The UPR is activated in different cell types during muscle regeneration.
- Targeted deletion of XBP1 impairs muscle regeneration in adult mice
- Myofiber XBP1 regulates satellite cell dynamics during regenerative myogenesis
- Myofiber XBP1 regulates abundance of non-myogenic cells in regenerating muscle

## INTRODUCTION

The skeletal muscle is composed of post-mitotic muscle cells called myofibers that are formed by the fusion of several mononucleated myoblasts during embryonic development. The regenerative capacity of skeletal muscle is attributed to the presence of muscle stem cells, called satellite cells, which reside between sarcolemma and basal lamina in a mitotically quiescent state (Yin et al., 2013). Following muscle damage, satellite cells undergo several rounds of proliferation followed by their differentiation into myoblasts. Finally, myoblasts fuse with each other or with the damaged myofibers to accomplish muscle repair (Relaix et al., 2021; Sousa-Victor et al., 2022; Yin et al., 2013). While satellite cells are critical for skeletal muscle regeneration, successful muscle regeneration involves the participation of several other cell types, such as neutrophils, macrophages, lymphocytes, fibro-adipogenic progenitors (FAPs), and endothelial cells (Sousa-Victor et al., 2022; Tidball, 2017). In addition, muscle regeneration involves the coordinated activation of an array of signaling pathways that are activated not only in satellite cells but also in damaged myofibers and other cell types that support muscle regeneration (Dumont et al., 2015a).

Skeletal muscle regeneration is an energy-dependent process, which involves the synthesis of many growth factors and a new set of cytoskeletal, membrane, and contractile proteins. Endoplasmic reticulum (ER) is the major site for protein synthesis and folding in mammalian cells, including skeletal muscle (Harding et al., 1999). In many conditions, which involve increased demand of protein synthesis, the protein-folding capacity of the ER lumen is diminished mainly due to the accumulation of misfolded or unfolded proteins - a state frequently referred to as ER stress. The stress in the ER leads to the activation of intracellular signal pathways called unfolded protein response (UPR) which is initiated by the phosphorylation and dimerization of protein kinase R (PKR)-like ER kinase (PERK) and inositol-requiring enzyme 1 (IRE1) and the proteolysis of activating transcription factor 6 (ATF6). The UPR attenuates stress in the ER by inhibiting translation, degrading mRNA and proteins, and increasing the folding capacity in the ER lumen (Hollien et al., 2009; Hollien and Weissman, 2006; Maurel et al., 2014; Wang and Kaufman, 2014). While the activation of UPR is a physiological response aimed at restoring homeostasis, chronic ER stress induces prolonged activation of the UPR, termed the “maladaptive UPR” or ER overload response (EOR), which can lead to deleterious consequences, such as insulin resistance, inflammation, and cell death (Hetz et al., 2019).

Accumulating evidence suggests that the components of the UPR pathways play important roles in the regulation of satellite cell function and skeletal muscle regeneration (Afroze and Kumar, 2019; Bohnert et al., 2018; Roy et al., 2024). For example, PERK-mediated signaling in satellite cells is essential for their self-renewal and for the regeneration of adult skeletal muscle (Xiong et al., 2017; Zismanov et al., 2016). IRE1α is the most conserved branch of the UPR that plays a major role in resolving ER stress. The activation of IRE1 leads to three major downstream outputs: the activation of c-Jun N-terminal kinase (JNK), the splicing of XBP1 mRNA, and the degradation of targeted mRNA and microRNAs, a process referred to as regulated IRE1-dependent decay (RIDD) (Hollien et al., 2009; Hollien and Weissman, 2006; Maurel et al., 2014; Wang and Kaufman, 2014). Recent studies have demonstrated that IRE1α/XBP1 signaling in myofibers promotes skeletal muscle regeneration in wild-type mice and in the mdx model of Duchenne muscular dystrophy (He et al., 2021; Roy et al., 2021). However, the cellular and molecular mechanisms through which myofiber IRE1α/XBP1 signaling regulates muscle regeneration remain largely unknown.

The emergence of single-cell RNA sequencing (scRNA-seq) has opened a new era in cell biology where cellular identity and heterogeneity can be defined by transcriptome dataset (Kolodziejczyk et al., 2015). In addition, single-nucleus RNA sequencing (snRNA-seq) has been developed as an alternative or complementary approach to characterize cellular diversity in tissues where there are difficulties in isolating intact cells (e.g., skeletal muscle, kidney, and bone) for transcriptome profiling due to their large size, tight interconnections, and fragility (Denisenko et al., 2020; Habib et al., 2017). Indeed, scRNA-seq was recently used to delineate cellular diversity at different stages of muscle regeneration in adult mice (De Micheli et al., 2020; Oprescu et al., 2020). Furthermore, the snRNA-seq approach has been used to understand the transcriptional heterogeneity in multinucleated skeletal muscle in normal and disease conditions (Chemello et al., 2020; Kim et al., 2020; Pass et al., 2023; Petrany et al., 2020).

In the present study, we first analyzed the scRNA-seq dataset to understand how the markers of ER stress/UPR are regulated in various cell types present in regenerating skeletal muscle of mice at different time points after injury. By performing snRNA-seq on skeletal muscle of myofiber-specific *Xbp1*-knockout mice, we investigated the role of XBP1 in the activation of downstream molecular pathways in the injured muscle microenvironment. Our results demonstrate the temporal activation of various markers of ER stress/UPR, ERAD, and ER overload response (EOR) in different cell types at various time points after injury. Moreover, snRNA-seq revealed that in addition to regulating satellite cell function, myofiber XBP1 regulates the activation of proteolytic systems, mitochondrial function, and abundance of various non-myogenic cells in regenerating skeletal muscle of adult mice.

## RESULTS

### Activation of UPR during muscle regeneration

We first sought to investigate how the gene expression of various components of ER stress and UPR are regulated in different cell types present in injured muscle microenvironment. Using the published scRNA-seq dataset (GSE143435) about muscle regeneration in mice (De Micheli et al., 2020), we first confirmed the presence of various cell types, such as muscle progenitor cells, mature skeletal muscle, fibro-adipogenic progenitors (FAPs), endothelial cells, tenocytes, proinflammatory and anti-inflammatory macrophages among other cell types in skeletal muscle of mice at day (D) 0, 2, 5, and 7 after muscle injury (**Supplemental Fig. S1A**). Consistent with the time course of muscle injury and regeneration (Dumont et al., 2015a; Sousa-Victor et al., 2022), the proportion of a few cell types, such as macrophages and other immune cells was drastically increased at day 2 and 5 and decreased at day 7 after injury. Similarly, proportion of endothelial cells and smooth muscle cells (SMCs) was reduced at day 2 and then gradually increased at day 5 and 7 post-injury. This analysis also confirmed that abundance of muscle progenitor cells was at peak at day 5 and then reduced at day 7 post-injury confirming that scRNA-seq dataset recapitulate the changes in the proportion of different cell types observed during skeletal muscle regeneration (**Fig. S1B**).

We next investigated how the gene expression of various markers of the UPR and ER-associated degradation (ERAD) are regulated in different cell types during muscle regeneration. We used the UPR gene set associated with GO term UPR or ERAD. Results showed that a few UPR molecules such as, *Atf4*, *Hspa5* and *Stub1* were constitutively expressed in most cell types present in uninjured and injured skeletal muscle **(Fig. 1)**. In contrast, there were certain molecules expressed only in specific cell types and their expression levels changed at different stages of muscle regeneration. For example, while *Atf3* is expressed in abundance in macrophages, Schwann cells, glial cells, and SMCs, it is highly expressed at D0 in muscle progenitor cells and lymphocytes and its levels are reduced after muscle injury. There were also some markers of the UPR (e.g., *Atf4*, *Ccnd1*, *Eif2a*, *Serp1*, *Xbp1*, *Derl1*, and *Creb3l1*) that showed increased expression in muscle progenitor cells following muscle injury. While mature myofibers showed relatively lower expression of various markers of the UPR, a few molecules (e.g., *Atf4*, *Herpud2*, *Stub1*, *Vapb*, *Xbp1*, and *Derl1*) were induced in response to injury.

**Figure 1.**
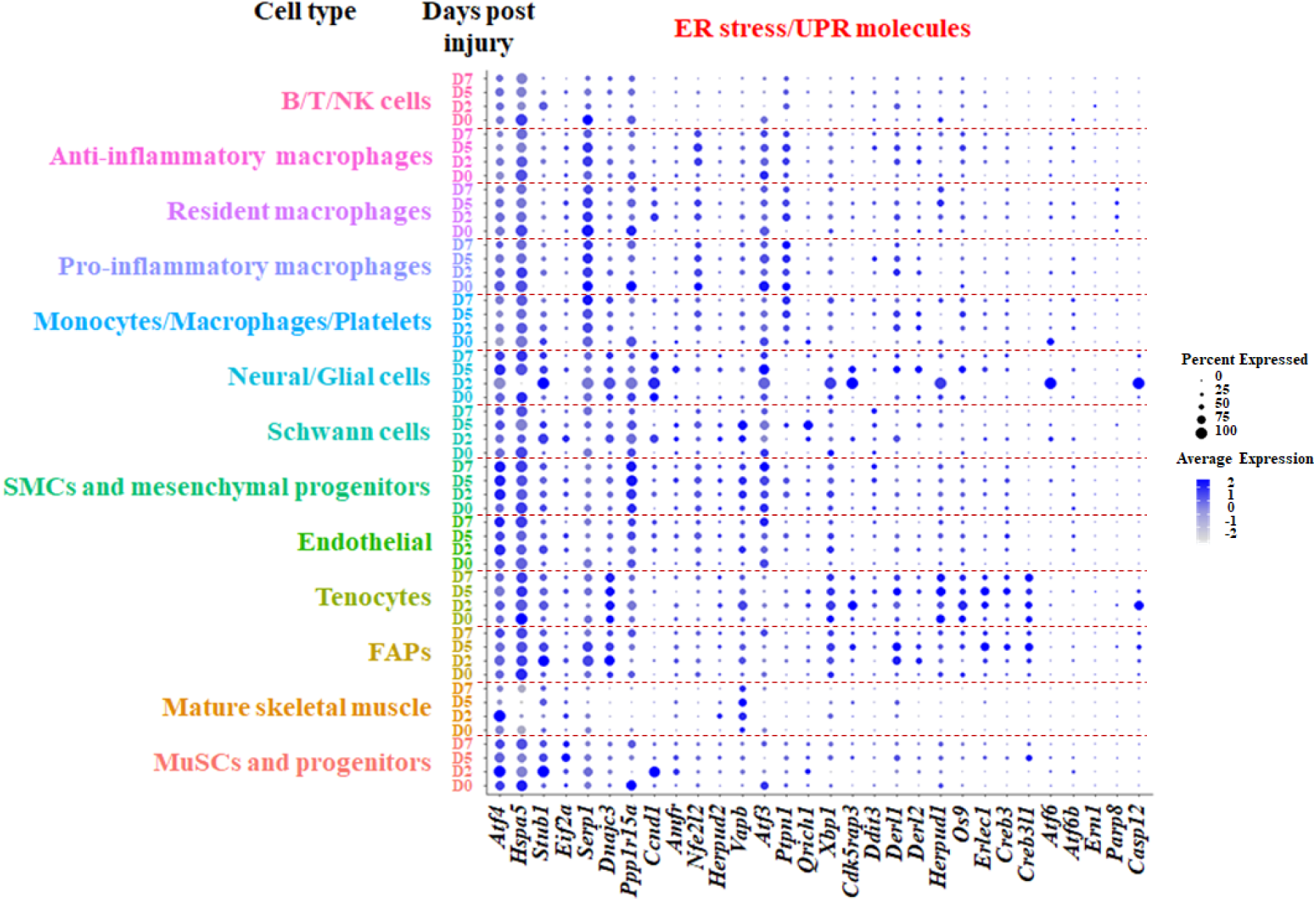
Gene expression of UPR molecules during muscle regeneration. The scRNA-seq dataset (GSE143437) was analyzed using R software (v4.2.2). Dot plot showing the changes in gene expression of ER stress/UPR molecules in different cell types and at different time points during muscle regeneration.

Interestingly, the gene expression of *Vapb*, which is required for ER protein quality control, was more pronounced in mature myofibers compared to muscle progenitor cells in regenerating skeletal muscle (**Fig. 1**).

Like UPR markers, we also found increased expression of a few ERAD-related molecules (e.g., *Calr*, *Hsp90b1*, and *Sec61b*) in all cell types with no to minimal changes at different time points following muscle injury (**Fig. S2**). In contrast, a few molecules were highly up regulated in response to muscle injury. For instance, the expression of *Canx*, *Psmc6*, *Sgta*, *Ube2j2*, *Ubxn4*, *Ube2j1*, *Ube2g2*, *Get4*, *Rcn3*, *Ubqln1*, *Aup1*, *Tor1*, and *Ubxn6* was increased in muscle progenitor cells at D2 and D5 following muscle injury. There were also specific ERAD molecules (i.e., *Dnajb9*, *Dnajb2*, *Faf1*, and *Fbxo6*) which were specifically induced in mature muscle cells following injury. In addition to myogenic cells, the markers of ERAD were also found to be upregulated in other cell types such as macrophages, neural cells, tenocytes, and FAPs. Remarkably, the basal level of expression of Rcn3 was high in FAPs, tenocytes, and glial cells that was further increased upon muscle injury suggesting the cell type specific regulation of ERAD during muscle regeneration **(Fig. S2)**.

Using the same dataset, we also examined the expression of EOR genes across different cell types at time points following muscle injury. Gene expression of several EOR molecules was found to be increased in different cell types, including muscle progenitor cells (e.g., *Ccd47*, *Ppp1r15b*, *Tmco1*, *Trp53*, *Bax*, and *Ube2k*) and mature muscle cells (*Gsk3b*, *Atp2a1*, *Spop*, *Atg10*, *Itpr1*, *Ube2k*, and *Aifm1*). Similar to muscle progenitor cells, we found a few molecules (e.g. *Atp2a1*, *Spop*, *Atg10*) were highly expressed in mature myofibers compared to muscle progenitor cells. Our analysis also showed that some other cell types, such as macrophages, glial cells, Scwann cells, SMCs, endothelial cells, FAPs, and tenocytes show variable gene expression of EOR molecules **(Fig. S3)**. Altogether, these results suggest that the markers of UPR, ERAD and EOR are induced in various cell types present in regenerating muscle of adult mice.

### Distinct clusters of nuclei in regenerating muscle identified by snRNA-seq

We have previously reported that myofiber-specific ablation of IRE1α (gene name: *Ern1*) attenuates skeletal muscle regeneration in response to injury in adult mice through its major downstream effector, the X-box protein 1 (XBP1) transcription factor (Roy et al., 2021). To understand how myofiber XBP1 regulates the transcriptomic profile in muscle and other cell types in injured muscles, we employed muscle specific *Xbp1* knockout (henceforth *Xbp1^mKO^*) and littermate control (i.e., *Xbp1^fl/fl^*) mice as described (Bohnert et al., 2019; Roy et al., 2021). The TA muscle of mice was injured by intramuscular injection of 1.2% BaCl_2_ solution, whereas contralateral uninjured muscle served as control. Muscle tissues were collected on day 5 or 21 post injury, followed by performing histological analysis and snRNA-seq (**Fig. 2A**). Consistent with our previously published report (Roy et al., 2021), average cross-sectional area (CSA) of newly formed myofibers and number of myofibers containing two or more centrally localized nuclei were significantly reduced in 5d-injured TA muscle of *Xbp1^mKO^* mice compared with corresponding TA muscle of *Xbp1^fl/fl^* mice (**Fig. 2B-D**). Moreover, there was also a significant reduction in the average myofiber CSA of regenerating myofibers in TA muscle of *Xbp1^mKO^* mice compared to *Xbp1^fl/fl^* mice on day 21 post-injury (**Fig. S4A, B**) confirming that genetic ablation of XBP1 in myofibers inhibits skeletal muscle regeneration in adult mice.

**Figure 2.**
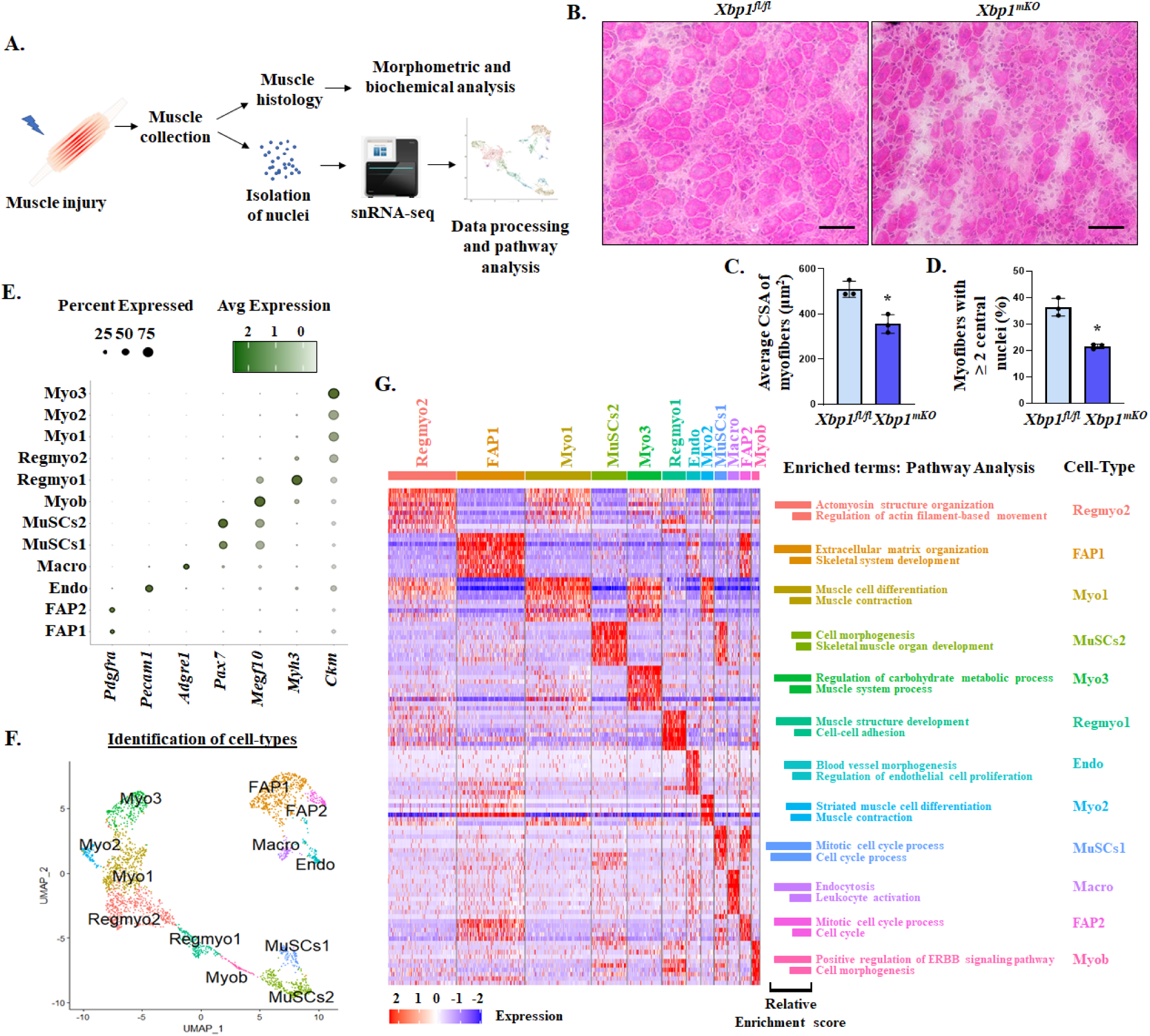
Single nucleus RNA-sequencing (snRNA-seq) analysis identifies different cell types in regenerating muscle. **(A)** TA muscle of *Xbp1^fl/fl^* and *Xbp1^mKO^*mice was injured using intramuscular injection of 1.2% BaCl_2_ solution. Schematics presented here show TA muscle histological analysis or isolation of nuclei followed by performed by snRNA-seq. **(B)** Representative photomicrographs of H&E-stained transverse sections of 5d-injured TA muscle of *Xbp1^fl/fl^* and *Xbp1^mKO^*mice. Scale bar, 50 µm. **(C)** Average myofiber cross sectional area (CSA) and **(D)** proportion of myofibers containing two or more centrally located nuclei. n=3 mice in each group. Data are presented as mean ± SEM. *p ≤ 0.05, values significantly different from corresponding muscle of *Xbp1^fl/fl^* mice analyzed by unpaired Student *t* test. Pre-processed 10X Genomics sequencing data of 5d-injured muscle of *Xbp1^fl/fl^* and *Xbp1^mKO^* was analyzed on R software for the presence of nuclei of different cell types. **(E)** Dot plot representing the proportion of cells and average expression of known genes associated with distinct cell types. **(F)** UMAP plot representing manually annotated clusters for cell type identity. **(G)** Validation of cellular identities by differentially expressed genes (DEG) followed by pathway enrichment analysis. Representative heatmap showing top 10 enriched genes per cluster (left panel) and enriched terms and cell type identities for corresponding clusters (right panel).

We next performed snRNA-seq on 5d-injured TA muscle of *Xbp1^fl/fl^* and *Xbp1^mKO^* mice followed by analysis of transcriptome data using bioinformatics tools. Seurat objects for *Xbp1^fl/fl^* and *Xbp1^mKO^* groups were individually processed for quality control, normalization, dimensionality reduction, clustering and elimination of doublet reads in an unbiased manner. Only the nuclei expressing 500 to 20,000 genes (nFeature_RNA), and less than 5% mitochondrial genes (percent.mt) were selected for further analysis **(Fig. S4C)**. The individual objects were then integrated to achieve homogeneous normalization of clusters between the two groups. We observed 12 spatially distributed nuclei clusters in the integrated object, which were visualized through Uniform Manifold Approximation and Projection (UMAP) plot **(Fig. S4D)**.

For the annotation of cluster identities, we analyzed the expression of specific gene markers, which have been consistently used for cellular identification, such as *Pax7* for satellite cells (MuSCs); *Megf10* for myoblasts (Myob); *Myh3* for regenerating (eMyHC^+^) muscle cells (Regmyo); *Ckm* for mature myofibers (Myo); *Pdgfra* for FAPs; *Adgre1* for macrophages (Macro); and *Pecam1* for endothelial (Endo) cells (Agarwal et al., 2020; Chemello et al., 2020; Holterman et al., 2007; Jaynes et al., 1988; Seale et al., 2000; Waddell et al., 2018). Dot plot representation showed exclusive and distinct expression of these marker genes to the spatially distributed clusters, which were manually annotated to the corresponding cell-type identities as shown in the UMAP plot **(Fig. 2E, F)**. This annotation showed that the nuclei of Myob, Endo and Macro cells were observed in single discrete clusters, whereas nuclei of MuSCs, Regmyo, Myo, and FAP cells were observed in more than one cluster. To validate the cell-type identification and to understand the observed multi-clustering of nuclei corresponding to the same identity, we analyzed the Gene Ontology (GO) biological processes and pathways associated with the distinct gene expression profiles of each cluster. The differentially expressed genes (DEGs), with the threshold of Log2FC ≥ |1| and p-value < 0.05, were identified using the ‘FindAllMarkers’ function across all clusters and the enriched genes in each cluster were used for pathway analysis using Metascape Gene Annotation and Analysis resource. A heatmap showing the top 10 DEGs per cluster is presented in **Fig. 2G**.

We next proceeded to validate clusters’ identity and characterize nuclei functionality. Enriched genes in the nuclei of MuSCs1 cluster showed an association with cell cycle regulation, whereas those for MuSCs2 nuclei were associated with skeletal muscle organ development and cell morphogenesis, suggesting that MuSCs1 nuclei resemble proliferating satellite cells while MuSCs2 nuclei resemble satellite cells committed to differentiation **(Fig. 2G)**. Indeed, we found multiple mitosis-associated genes, including kinesin superfamily members (*Kif4*, *Kif11*, *Kif15*, *Kif23*, *Kif20b*, and *Kif24*), Centromere protein E and F (*Cenpe*, *Cenpf*), and *Diaph3* among others, enriched in nuclei of MuSCs1, i.e., in proliferating satellite cells **(Fig. S5A)**. In contrast, MuSCs2 nuclei included enriched genes, such as *Meg3*, *Megf10*, *Cdon*, *Met*, *Dag1*, *Dmd*, *Tgfbr3*, *Heyl*, and *Notch3* **(Fig. S5B)**. Satellite cell differentiation and migration is positively regulated by many genes, including maternally expressed gene 3 (*Meg3*) (Cheng et al., 2020; Liu et al., 2023), *Megf10* (Holterman et al., 2007), Cell adhesion associated oncogene related (*Cdon*) (Bae et al., 2020), and HGF-receptor (c-Met or *Met*) (Webster and Fan, 2013). Moreover, the dystrophin-associated glycoprotein complex-encoding genes, such as *Dag1* and Dystrophin (*Dmd*), which were enriched in MuSCs2 cluster, have been implicated to play a crucial role in the regulation of satellite cell polarity and asymmetric division (Dumont et al., 2015b), thereby, governing self-renewal of satellite cells. Similarly, the Notch receptor, *Notch3*, and the Notch target gene, *Heyl*, have roles in the maintenance of satellite cell quiescence (Bjornson et al., 2012; Gioftsidi et al., 2022). Furthermore, a recent study asserted TGFβ-receptor 3 (*Tgfbr3*) as a unique marker of self-renewing MuSCs (Okafor et al., 2023). Therefore, the MuSCs2 cluster harbors self-renewing satellite cells in addition to those committed to myogenic lineage.

Myoblasts (Myob cluster) are proliferating mononucleated cells that differentiate and fuse with injured myofibers, leading to muscle repair. ERBB receptors (ERBB1-4) play a crucial role in an array of cellular functions including cell growth, proliferation, apoptosis, migration, and adhesion and ERBB2 positively regulates myoblast cell survival (Andrechek et al., 2002). Enriched genes in the Myob nuclei cluster were associated with the positive regulation of ERBB signaling pathway and the process of cell morphogenesis **(Fig. 2G)**.

Regenerating myofibers (Regmyo clusters) are newly formed muscle cells that express the embryonic isoform of myosin heavy chain (eMyHC; gene name: *Myh3*) (Agarwal et al., 2020; Yin et al., 2013). Our analysis showed the presence of two clusters of *Myh3*^+^ (Regmyo) nuclei in regenerating muscle **(Fig. 2G)**. Investigation of the enriched genes in Regmyo1 cluster showed an association with muscle structure development and cell-cell adhesion whereas Regmyo2 nuclei showed association with actomyosin structure organization and regulation of actin filament-based movement. Fusion-competent myoblasts highly express genes regulating membrane proteins required for both cell adhesion and cell-cell fusion (Hindi and Millay, 2022; Hindi et al., 2013). Many fusion-related molecules have now been identified, including Myomaker (*Tmem8c*), N-cadherin (*Cdh2*), Myoferlin (*Myof*), Caveolin 3 (*Cav3*), and Nephronectin (*Npnt*) (Hindi et al., 2013; Millay et al., 2013; Millay et al., 2014). We observed that the genes encoding for all these profusion molecules were highly enriched in the nuclei of Regmyo1 cluster **(Fig. S6A)**. In contrast, there was higher expression of neonatal (or perinatal) isoform of MyHC (neo-MyHC; gene name: *Myh8*), which is also expressed in regenerating muscle, in the nuclei of Regmyo2 cluster. In addition, we observed enrichment of genes related to muscle growth and maturation (*Myh4*, *Ctnna3*, *Igfn1*, *Myoz1*) (Cracknell et al., 2020; Schiaffino et al., 2015; Wu et al., 2022) and Ca^2+^ handling and metabolism-related genes (*Gpt2*, *Rora*, *Stim1*, and *Pgm2*) (Cicatiello et al., 2022; Conte et al., 2021; Lau et al., 2004; Marceca et al., 2020) in the nuclei of Regmyo2 cluster **(Fig. S6B)**, suggesting structural growth and metabolic adaptation of regenerating myofibers.

Regenerated myofibers express high levels of muscle creatine kinase (*Ckm*), a marker of mature myofibers (Myo clusters). Interestingly, our snRNA-seq analysis showed three distinct clusters of *Ckm*^+^ nuclei (Myo1, 2 and 3). Enriched genes in Myo1 and 2 clusters were associated with biological processes of muscle cell differentiation and muscle contraction, whereas enriched genes in Myo3 clusters were associated with pathways related to carbohydrate metabolism and muscle system process **(Fig. 2G)**. We first investigated the potential reasoning of a three-cluster division of *Ckm*^+^ nuclei. All these nuclei expressed *Ttn* (Titin), a pan-muscle marker. In addition, *Myh1* (Type IIX) and *Myh4* (Type IIB) genes were readily observed as compared to *Myh2* (Type IIA) while the expression pattern was not distinct in nuclei clusters **(Fig. S7A)**, suggesting that nuclear heterogeneity in the *Ckm*^+^ clusters was not due to muscle fiber type. We then investigated the differentially expressed genes to understand the distribution of *Ckm*^+^ nuclei. We found multiple common enriched genes in Myo1 and Myo2 clusters, however, *Myh8* and *Col24a1* genes were significantly enriched in the Myo1 cluster compared to Myo2 **(Fig. S7B)**, suggesting that the Myo1 nuclei exhibit genes potentially involved in terminal differentiation and structural maturity in continuation to the Regmyo2 cluster. Further investigation showed that the Myo2 cluster was highly enriched in lncRNA genes located within the *Dlk1*-*Dio3* locus, including maternally imprinted *Meg3* and *Mirg*, and paternally expressed *Rtl1* **(Fig. S7C)**. Finally, many genes involved in muscle hypertrophy and metabolic activity (*Hs3st5*, *Rcan2*, *Cd36*, *Mylk4*, *Kcnn2*, *Osbpl6*, *Fgf1*, *Pfkfb1*, *Pdk4*) were enriched in the Myo3 cluster suggesting functional adaptation and hypertrophic growth of regenerated myofibers **(Fig. S7D)**.

Muscle niche also involves many other cell types, including macrophages (Macro), endothelial (Endo) cells, and FAPs that play important roles in muscle regeneration following acute damage (De Micheli et al., 2020; Sousa-Victor et al., 2022; Tidball, 2017). Our snRNA-seq analysis identified nuclei pertaining to these cell-types in clusters that are spatially distributed away from the muscle nuclei. Identification of DEGs followed by biological process and pathway enrichment analysis showed that enriched genes in the nuclei of the Macro cluster were associated with endocytosis and leukocyte activation, while those for Endo nuclei showed association with blood vessel morphogenesis and the regulation of endothelial cell proliferation **(Fig. 2G)**. In contrast, analysis of gene expression in the nuclei of FAP clusters, FAP1 and 2, showed association with the biological processes of extracellular matrix (ECM) organization and cell cycle, respectively. This suggests that FAP2 nuclei resemble proliferating cells whereas FAP1 nuclei contribute to ECM organization, which is vital to skeletal muscle regeneration **(Fig. 2G)**. Altogether, the cellular identities annotated to each cluster are validated for investigating the mechanisms of action of myofiber XBP1 in regenerative myogenesis.

### XBP1 regulates proteolytic systems and mitochondrial function during regenerative myogenesis

To understand the mechanisms by which myofiber XBP1 promotes muscle regeneration, we examined various features, including the spatial distribution of clusters, the abundance of nuclei for each cellular identity, and differentially expressed genes in the myonuclear populations of *Xbp1^fl/fl^* and *Xbp1^mKO^*mice.

The spatial distribution of clusters was similar between *Xbp1^fl/fl^* and *Xbp1^mKO^* objects, visualized using split-UMAP plots **(Fig. 3A)**. Due to a difference in the total number of sequenced nuclei for the two objects (2816 nuclei for *Xbp1^fl/fl^*and 2140 for *Xbp1^mKO^*), we analyzed the proportion of nuclei per cluster instead of the absolute number of nuclei. This analysis showed a marked reduction in the proportion of nuclei in the clusters of MuSCs1 and 2 (∼3-fold), Myob (∼5-fold), and Regmyo1 (∼1.5-fold); a modest increase (∼1.1-fold) in the proportion of nuclei in Regmyo2 and Myo3; and a significant increase in the clusters Myo1, 2 and 3 (∼2-, 2- and 3-fold, respectively) in the regenerating TA muscle of *Xbp1^mKO^* mice compared to littermate *Xbp1^fl/fl^* mice **(Fig. 3B)**.

**Figure 3.**
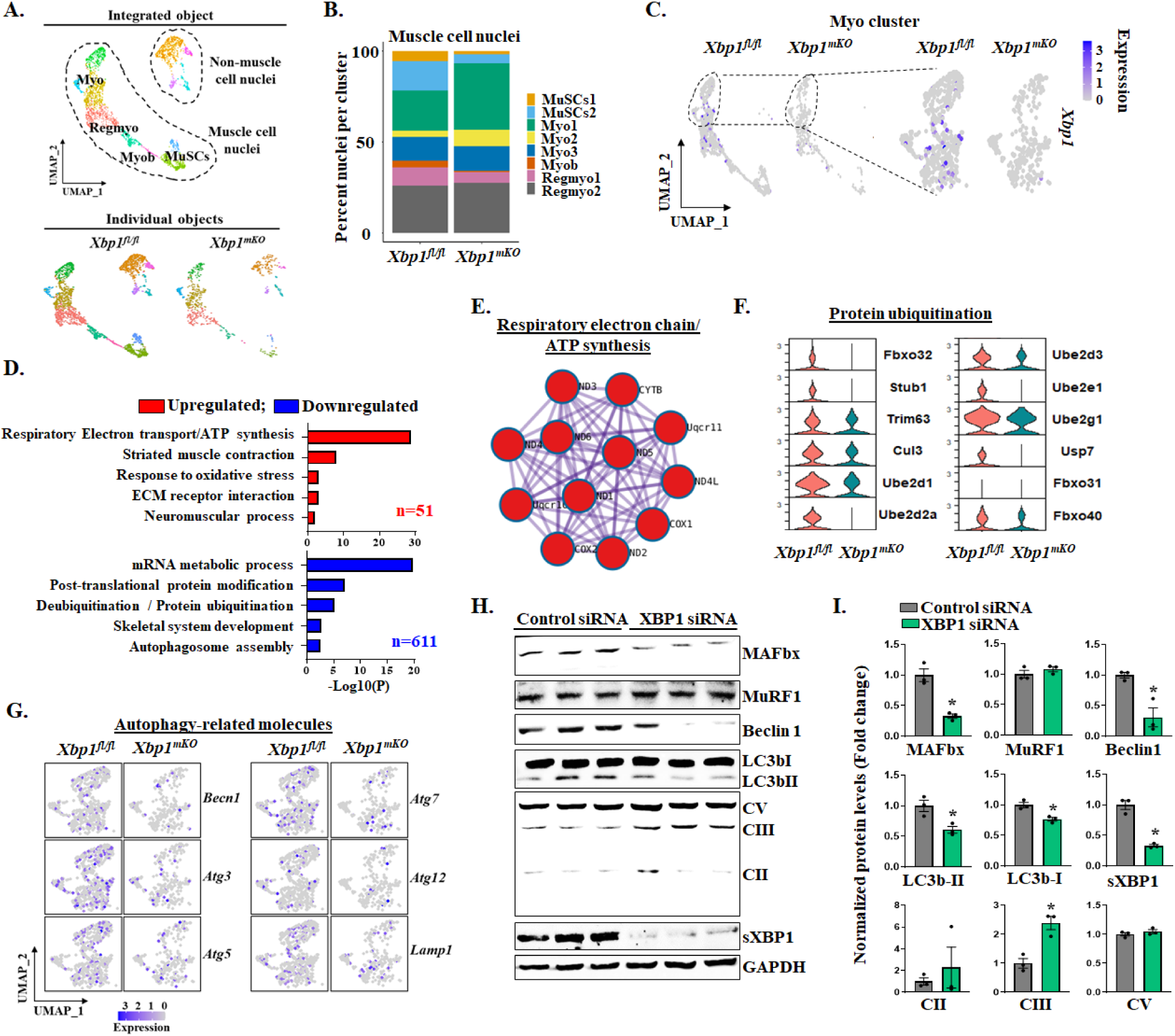
XBP1 regulates proteolytic pathways and mitochondrial OXPHOS levels in regenerating muscle. **(A)** The integrated Seurat object of the injured muscles of *Xbp1^fl/fl^*and *Xbp1^mKO^* mice were classified based on myogenic and non-myogenic nuclei. **(B)** Proportion of nuclei in different clusters of myogenic cells in 5d-injured TA muscle of *Xbp1^fl/fl^* and *Xbp1^mKO^*mice. **(C)** Feature plot showing gene expression of *Xbp1* in all muscle nuclei or mature myofiber (Myo) cluster (right panel) of *Xbp1^fl/fl^*and *Xbp1^mKO^* mice. **(D)** Differentially expressed genes (DEG) in Myo clusters were identified and used for pathway enrichment analysis using Metascape. Bar graphs show enriched biological processes and pathways associated with upregulated (red) and downregulated (blue) genes in *Xbp1^mKO^*compared to *Xbp1^fl/fl^* mice. **(E)** Protein-protein interaction plots of the upregulated genes associated with respiratory electron chain/ATP synthesis pathway. **(F)** Violin plot showing downregulation of gene expression of protein ubiquitination-related molecules and **(G)** Feature plots showing downregulation of gene expression of autophagy-related molecules in *Xbp1^mKO^* mice compared to *Xbp1^fl/fl^* mice. **(H)** Primary myoblast cultures were incubated in differentiation medium for 48 h followed by transfection with control or XBP1 siRNA. Myotubes were collected after 24 h of transfection and cell lysates were used for immunoblotting. Immunoblots presented show protein levels of MAFbx (*Fbxo32*), MuRF1 (Trim63), Beclin1, LC3B, OXPHOS complexes, sXBP1, and unrelated protein GAPDH in myotubes transfected with control or XBP1 siRNA. **(I)** Quantification of protein levels of MAFbx, MuRF1, Beclin1, LC3bI and II, sXBP1 and OXPHOS complexes CII, III and V. n=3 biological replicates. Data are presented as mean ± SEM. *p ≤ 0.05; values significantly different from cultures transfected with control siRNA analyzed by unpaired Student *t* test.

Since our knockout model ablates XBP1under the promoter of *Ckm* gene, we first checked the expression of XBP1 in *Xbp1^fl/fl^* and *Xbp1^mKO^* mice and subsequently analyzed the XBP1-mediated gross transcriptomic alterations across all *Ckm*^+^ nuclei. A single subset group containing Myo nuclei (Myo1, 2 and 3) was created from the initial processed data for downstream analysis. Deletion of *Xbp1* was confirmed using split-UMAP feature plots **(Fig. 3C)**. Next, we analyzed the DEGs (Log2FC ≥ |0.25| and p-value < 0.05) in *Xbp1^mKO^ Ckm*^+^ myonuclei compared to corresponding nuclei of controls. This analysis showed that 662 genes were significantly dysregulated in *Xbp1^mKO^* mice compared to *Xbp1^fl/fl^* mice. Strikingly, 611 of these genes were downregulated while only 51 genes were upregulated. Pathway enrichment analysis revealed that the upregulated genes were associated with respiratory electron transport, striated muscle contraction, response to oxidative stress, ECM receptor interaction, and neuromuscular process **(Fig. 3D)**. Protein-protein interaction models for upregulated genes showed gene clusters related to respiratory electron chain/ATP synthesis process suggesting increased gene expression of molecules related to oxidative phosphorylation **(Fig. 3E)**. GO term and pathway analysis showed that downregulated gene sets were associated with mRNA metabolic process, post-translational protein modification, protein ubiquitination and deubiquitination, skeletal system development and autophagosome assembly **(Fig. 3D)**.

In response to ER stress, activated sXBP1 protein translocate to the nucleus and regulates the gene expression of multiple molecules involved in enhancing the protein folding capacity of the ER and/or promoting the degradation of unfolded or misfolded proteins by a process called ERAD (Hetz et al., 2019). We analyzed the expression levels of various genes associated with the protein ubiquitination and autophagy process. A significant downregulation in gene expression of multiple molecules of ubiquitination-proteasome system, including E3 ubiquitin ligases (*Fbxo32* (MAFbx), *Stub1*), ubiquitin ligase assembly-scaffolding protein (*Cul3*), E2 ubiquitin enzymes (*Ube2d1*, *Ube2d2a*, *Ube2d3*, *Ube2e1*, *Ube2g1*), and deubiquitinating enzyme (*Usp7*) was observed in *Ckm*^+^ myonuclei of *Xbp1^mKO^* mice compared to *Xbp1^fl/fl^*mice. By contrast, the gene expression of the muscle specific E3 ubiquitin ligase *Trim63* (i.e. MuRF1) was comparable between the two genotypes **(Fig. 3F)**. We also observed a reduction in the number of nuclei expressing autophagy-related markers (*Becn1*, *Atg3*, *Atg5*, *Atg7*, *Atg12*, and *Lamp1*) in *Xbp1^mKO^* compared to *Xbp1^fl/fl^* group **(Fig. 3G)**.

To confirm the role of XBP1 in the regulation of components of mitochondrial respiratory chain, ubiquitin-proteasome system, and autophagy, we next studied the effect of knockdown of XBP1 on the levels of a few proteins related to these pathways in cultured mouse primary myotubes. Primary myoblasts isolated from hindlimb muscle of wild type mice were incubated in differentiation medium for 48 h followed by transfection with control or XBP1 siRNA. Western blot analysis showed that knockdown of XBP1 in cultured myotubes represses the levels of MAFbx (but not MuRF1), Beclin1, and LC3BII/LC3BI ratio whereas some of the components of OXPHOS complexes (CII, CIII, and CV) were increased **(Fig. 3H, I)**. While repression of the markers of ubiquitin-proteasome system and autophagy upon knockdown of XBP1 is consistent with our previous report (Bohnert et al., 2019), the increase in levels of OXPHOS proteins was quite intriguing. It is known that similar cellular mechanisms are involved in developmental, post-natal, and adult regenerative myogenesis (Kang and Krauss, 2010). To further investigate the impact of XBP1 deletion in muscle, we also investigated whether genetic deletion of XBP1 in myofibers also affects the levels of OXPHOS proteins during postnatal myogenesis. There was a significant increase in the levels of total OXPHOS protein, complex III, and complex V in TA and gastrocnemius (GA) muscle of 2-week-old *Xbp1^mKO^* mice compared to *Xbp1^fl/fl^* mice **(Fig. S8A-F)**. Surprisingly, there was no significant difference in the levels of OXPHOS proteins in GA muscle of 10-week-old *Xbp1^fl/fl^*and *Xbp1^mKO^* mice **(Fig. S8G-I)** suggesting that the levels of mitochondrial OXPHOS protein are transiently increased in regenerating/developing muscle of *Xbp1^mKO^*mice, which may be a compensatory mechanism to support the myogenesis in *Xbp1*-null myofibers. Altogether, these results suggest that XBP1 regulates the gene expression of various components of ubiquitin-proteasome system, autophagy, and mitochondrial function during regenerative myogenesis.

### XBP1 regulates the formation of new myofibers during regenerative myogenesis

We next investigated the transcriptomic alterations in the nuclei of *Myh3*-positive cells. Analysis of DEGs revealed significant changes in the gene expression of 968 molecules, with repression of 802 and upregulation of 166. Pathway enrichment analysis of the DEGs showed that the upregulated genes were associated with respiratory electron transport, mitochondrial biogenesis, muscle contraction, regulation of cell migration, and ECM organization, whereas the downregulated genes were associated with RHO GTPase cycle, muscle structure development, regulation of muscle cell and myotube differentiation, and endocytosis **(Fig. 4A)**. Further investigation of the deregulated genes involved in the identified biological processes and pathways showed multiple enriched genes, including *mt-Co1*, *mt-Co2*, *mt-Co3*, *mt-Nd4*, *Acta1*, *Neat1*, *Tnnc2*, *Camk1d*, and *Taco1* that regulate the respiratory electron transport and oxidative phosphorylation process **(Fig. 4B)**. We found downregulation of multiple genes, including *Cdh2*, *Tmem8c*, *Mef2c*, *Nfatc2*, *Nfatc3*, *Gsk3b*, *Cdon*, *Myoz1*, *Foxp1*, and *Rb1*, indicating an impairment in muscle structure development, potentially through impairment of fusion process **(Fig. 4C)**. Indeed, we have recently reported that XBP1 transcription factor induces the gene expression of multiple profusion molecules, including *Tmem8c* (also known as Myomaker) to promote myoblast fusion during myogenic differentiation (Joshi et al., 2024). While we have used muscle creatine kinase (MCK)-Cre line which is predominately expressed in differentiated muscle cells, it can also be expressed at low levels in satellite cells. Indeed, our RT-PCR and qPCR analysis showed that in addition to muscle tissues, there was also a small but significant reduction in the mRNA levels of XBP1 in freshly isolated satellite cells of *Xbp1^mKO^* mice compared to littermate *Xbp1^fl/fl^*mice (**Fig. S9**). This reduction in the XBP1 levels in satellite cells may be sufficient to reduce their fusion with injured myofibers of *Xbp1^mKO^*mice leading to the reduction in muscle regeneration.

**Figure 4.**
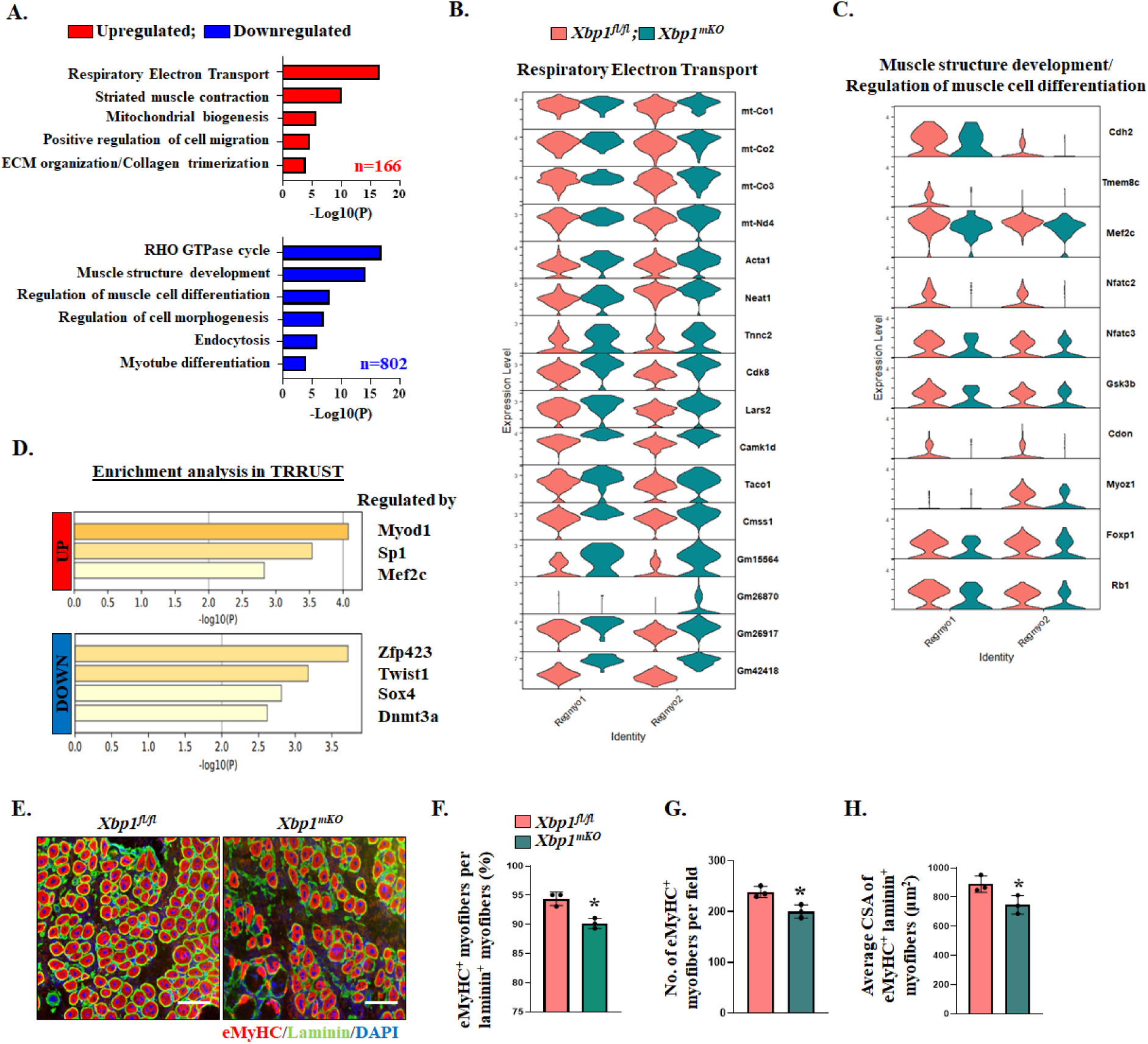
Myofiber XBP1 regulates formation of new myofibers during muscle regeneration. **(A)** Pathway enrichment analysis of differentially expressed genes in clusters of eMyHC^+^ regenerating myonuclei (Regmyo1 and Regmyo2) of *Xbp1^fl/fl^* and *Xbp1^mKO^*mice. Violin plots show gene expression of the **(B)** upregulated molecules associated with respiratory electron transport system and **(C)** downregulated molecules involved in muscle differentiation and structure development. **(D)** Enrichment analysis in TRRUST database showing transcriptional regulators of the upregulated and downregulated genes in *Xbp1^mKO^* mice compared to *Xbp1^fl/fl^* mice. **(E)** Transverse sections of 5d-injured TA muscle of *Xbp1^fl/fl^* and *Xbp1^mKO^* mice were immunostained for eMyHC and laminin protein. Nuclei were counterstained by DAPI. Representative photomicrographs demonstrating eMyHC^+^ regenerating myofibers in 5d-injured TA muscle. Scale bar, 50 µm. Quantitative analysis of **(F)** eMyHC^+^ myofibers per laminin^+^ myofibers, **(G)** number of eMyHC^+^ myofibers per field, and **(H)** average cross-sectional area of eMyHC^+^ laminin^+^ myofibers. n=3 mice in each group. Data are presented as mean ± SEM and analyzed by unpaired Student t test. *p ≤ 0.05; values significantly different from injured TA muscle of *Xbp1^fl/fl^* mice.

We next analyzed the key transcriptional regulators of the deregulated genes using the TRRUST (Transcriptional Regulatory Relationships Unraveled by Sentence-based Text mining) tool. Results showed that *Myod1*, *Sp1*, and *Mef2c* transcription factors are involved in the regulation of the upregulated genes, whereas the downregulated genes are potentially controlled by the transcriptional regulators, such as *Zfp423*, *Twist1*, *Sox4* and *Dnmt3a* **(Fig. 4D)**. To validate snRNA-seq analysis, we also performed immunostaining for eMyHC (gene name: *Myh3*) on 5d-injured TA muscle section of *Xbp1^fl/fl^*and *Xbp1^mKO^* mice, followed by quantitative analysis. Results showed that the number and cross-sectional area (CSA) of eMyHC^+^ myofibers were significantly reduced in 5d-injured TA muscle of *Xbp1^mKO^* mice compared to littermate *Xbp1^fl/fl^* mice (**Fig. 4E-H)**. Collectively, these results suggest that targeted ablation of XBP1 delays the formation of new myofibers during skeletal muscle regeneration in adult mice.

### XBP1 regulates chronological alterations along the myogenic lineage

Muscle regeneration is a highly coordinated process that involves stage specific activation of various myogenic regulatory factors and signaling pathways (Relaix et al., 2021). We next investigated whether genetic ablation of XBP1 alters the gene expression patterns along a pseudotime axis resembling the transition of muscle cells along the myogenic lineage. For this analysis, we selectively considered only the muscle cell nuclei (excluding the non-muscle cell nuclei from the entire nuclei population). Using Monocle2 package, gene expression patterns were analyzed, and the trajectory path was mapped along the pseudotime axis. As expected, we observed trajectory line originating from clusters of satellite cells, followed by myoblasts and regenerating myofibers, and eventually leading to the clusters of mature myofibers in *Xbp1^fl/fl^* mice. Interestingly, the trajectory analysis of *Xbp1^mKO^* myonuclei distinctly differed from the *Xbp1^fl/fl^*mice at the origin (satellite cells) and showed altered nodes in the clusters of regenerating myofibers **(Fig. 5A)**.

**Figure 5.**
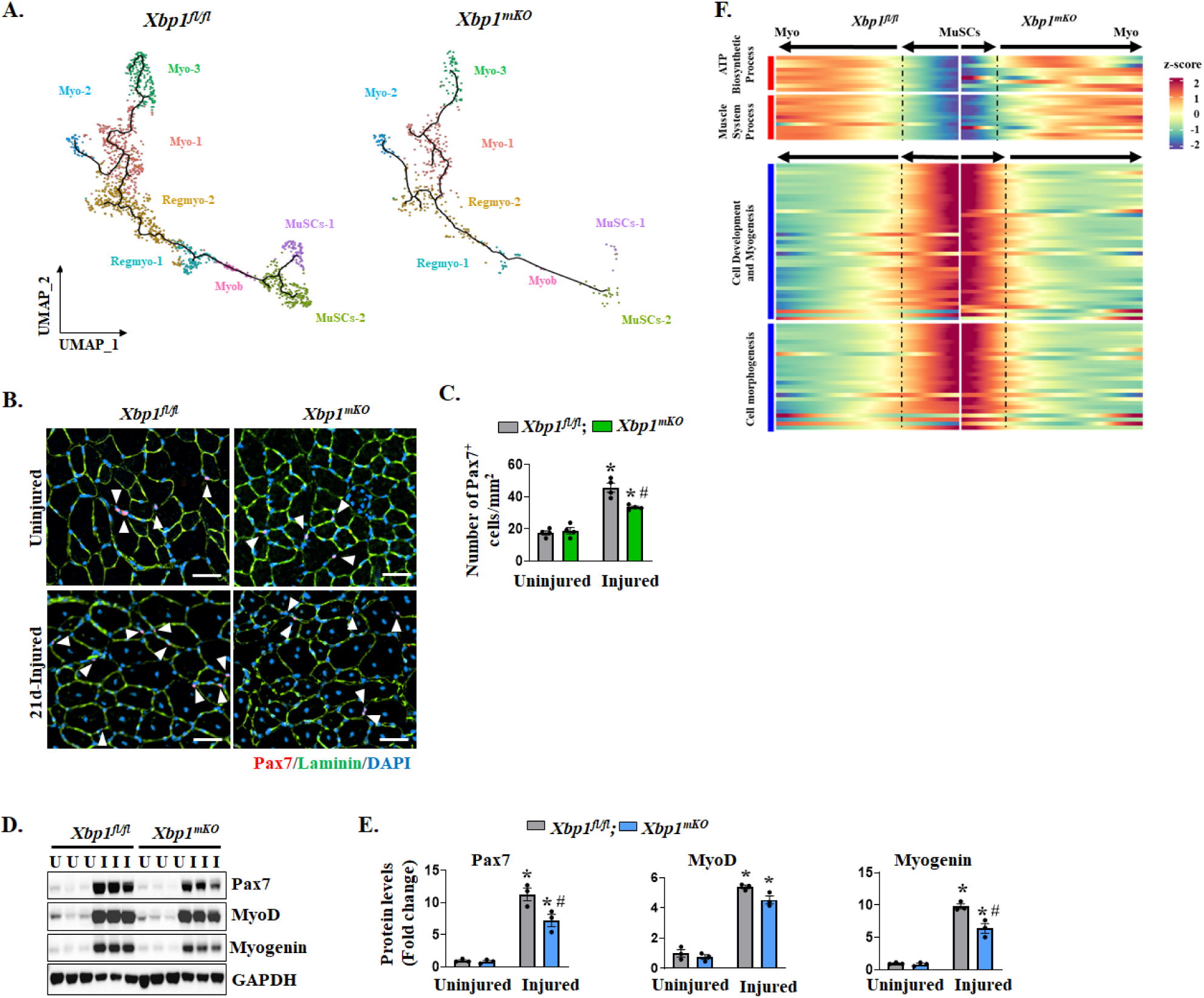
XBP1 regulates the myogenesis trajectory and alters the transcriptomic profiles of muscle progenitor cells during muscle regeneration. **(A)** Trajectory path of myonuclei along the pseudotime axis resembling the myogenic lineage was plotted using the Monocle2 package. UMAP plots show trajectories of myonuclei in injured TA muscles of *Xbp1^fl/fl^* and *Xbp1^mKO^* mice. **(B)** Representative images of uninjured and 21d-injured TA muscle sections of *Xbp1^fl/fl^* and *Xbp1^mKO^* mice after immunostaining for Pax7 and laminin protein. DAPI was used to identify nuclei. Scale bar, 50 µm. **(C)** Quantification of number of Pax7^+^ cells per unit area in uninjured and 21d-injured muscle of *Xbp1^fl/fl^* and *Xbp1^mKO^*mice. **(D)** Immunoblots, and **(E)** quantification of levels of Pax7, MyoD and Myogenin protein in uninjured and 5d-injured TA muscle of *Xbp1^fl/fl^*and *Xbp1^mKO^* mice. n=3-4 mice in each group. Data are presented as mean ± SEM. *p ≤ 0.05, values significantly different from corresponding uninjured muscle of *Xbp1^fl/fl^*and *Xbp1^mKO^* mice, and #p ≤ 0.05, values significantly different from 5d- or 21d-injured muscle of *Xbp1^fl/fl^* mice analyzed by two-way ANOVA, followed by Tukey’s multiple comparison test. **(F)** Heatmaps showing upregulated and downregulated gene sets along the pseudotime axis.

Consistent with our prior analysis, the MuSCs2 cluster of *Xbp1^fl/fl^* mice showed a division in the trajectory, one following the myogenic lineage (satellite cells committed to differentiation) and the other retracting away (self-renewing satellite cells). Interestingly, the trajectory path for *Xbp1^mKO^* mice originated from MuSCs2 cluster, rather than MuSCs1 (proliferating MuSCs). The limited proportion of proliferating MuSCs in *Xbp1^mKO^* mice might be a potential cause of the failure to recognize the MuSCs1 cluster in the trajectory model. However, unlike the *Xbp1^fl/fl^* mice, the trajectory line for *Xbp1^mKO^* mice did not show any division for self-renewing satellite cells suggesting that myofiber-specific deletion of XBP1 leads to an impairment in the satellite cell self-renewal ability during adult muscle regeneration **(Fig. 5A)**. Comparative analysis of the distribution of marker gene expression in conjunction with the nuclei population size between *Xbp1^fl/fl^* and *Xbp1^mKO^* mice clearly showed the reduction of *Pax7*, *Meg10*, and *Myh3* expressing nuclei along with an early and aberrant expression of *Ckm* gene in *Xbp1^mKO^* compared to *Xbp1^fl/fl^*group further suggesting early or premature differentiation at the expense of satellite cell population **(Fig. S10A)**.

We have previously reported that number of satellite cells are reduced in 5d-injured TA muscle of *Xbp1^mKO^*mice compared to *Xbp1^fl/fl^* mice (Roy et al., 2021). To validate the snRNA-Seq results about impact of myofiber-specific deletion of XBP1 on satellite cells during muscle regeneration, we performed immunohistochemistry for Pax7 on uninjured and 21d-injured TA muscle section of *Xbp1^fl/fl^* and *Xbp1^mKO^* mice. There was no significant difference in the number of Pax7^+^ cells in the uninjured TA muscle between the two genotypes. However, the number of satellite cells was significantly reduced in 21d-injured TA muscle of *Xbp1^mKO^* mice compared to corresponding muscle of *Xbp1^fl/fl^* mice (**Fig. 5B, C**). In addition, our western blot analysis showed that levels of Pax7 and Myogenin, but not MyoD, were significantly reduced in the 5d-injured TA muscle of *Xbp1^mKO^* mice compared to *Xbp1^fl/fl^*mice (**Fig. 5D, E**) further suggesting that myofiber-specific deletion of XBP1 inhibits the abundance of satellite cells during regenerative myogenesis.

We further analyzed the effect of myofiber-specific ablation of XBP1 on satellite cell dynamics by establishing single myofiber cultures from extensor digitorum longus (EDL) muscle of *Xbp1^fl/fl^* and *Xbp1^mKO^* mice. There was no significant difference in the number of Pax7^+^ or MyoD^+^ cells on freshly isolated EDL myofibers of *Xbp1^fl/fl^* and *Xbp1^mKO^* mice further suggesting that myofiber-specific deletion of XBP1 does not affect the abundance of satellite cells in uninjured muscle (**Fig. S11A-C**). We next performed Ki-67 staining at 48 h of culturing of EDL myofibers. Interestingly, the number of Ki67^+^ cells were significant reduced on cultured myofibers of *Xbp1^mKO^* mice compared with *Xbp1^fl/fl^* mice, suggesting that myofiber-specific deletion of XBP1 also inhibits proliferation of myofiber-associated satellite cells in an *ex vivo* model of muscle injury (**Fig. S11D, E**).

By performing immunostaining for Pax7 and MyoD protein as described (Hindi and Kumar, 2016; Roy et al., 2021), we also investigated the effect of myofiber-specific ablation of XBP1 on the self-renewal, proliferation, and differentiation of myofiber-associated satellite cells at 72 h of establishing the cultures. While there was no significant difference in the number of clusters per myofiber, there was a significant reduction in the number of cells per cluster in *Xbp1^mKO^*mice compared with *Xbp1^fl/fl^* mice (**Fig. S12A-C**). Our analysis also showed that there was a significant decrease in the number of self-renewing (Pax7^+^/MyoD^-^) and proliferating (Pax7^+^/MyoD^+^) cells per myofiber and a significant increase in the number of differentiating (Pax7^-^/MyoD^+^) cells per myofiber in *Xbp1^mKO^* cultures compared to *Xbp1^fl/fl^* cultures (**Fig. S12A, D, E, F**). These results suggest that in addition to reducing proliferation, myofiber-specific deletion of XBP1 inhibits self-renewal and induces precocious differentiation of satellite cells.

Further analysis of DEG of *Xbp1^mKO^* muscle nuclei compared to those of *Xbp1^fl/fl^* along the pseudotime axis revealed 697 deregulated genes, out of which 523 were upregulated and 174 were downregulated. GO term enrichment analysis showed that upregulated genes were involved in the processes of ATP biosynthesis, regulation of muscle contraction, striated muscle contraction, and muscle system process whereas downregulated genes were associated with cell morphogenesis, cellular component morphogenesis, cell development, and regulation of multicellular organismal development **(Fig. S10B)**. Differences in gene expression along the pseudotime were visually represented through heatmaps. Upregulated gene sets show distinct enrichment, whereas the downregulated genes show modest repression in their expression pattern in *Xbp1^mKO^* compared to *Xbp1^fl/fl^*mice **(Fig. 5F)**. Altogether, these results suggest that targeted deletion of XBP1 alters the temporal regulation of gene expression that leads to altered dynamics of satellite cells during muscle regeneration.

### Myofiber XBP1 regulates distinct molecular and signaling pathways in satellite cells

We next studied the transcriptomic alterations in the clusters of satellite cell nuclei (MuSCs1 and MuSCs2) by identifying the DEGs with a threshold of Log2FC ≥ |0.25| and p-value < 0.05 and by performing biological process enrichment analysis. Our analysis of MuSCs1 nuclei of *Xbp1^mKO^* mice showed 366 downregulated and 369 upregulated genes compared to *Xbp1^fl/fl^* group **(Fig. 6A)**. Moreover, biological process enrichment analysis for downregulated genes of MuSCs1 cluster revealed that multiple genes, including *Acvr1*, *Atf2*, *Ezh2*, *Cdk6*, *Kif11*, *Bub1*, *Bub3*, *Cdc42*, *Rrm1*, and *Smarcc1* showed association with the regulation of cell cycle process, whereas the genes *Runx1*, *Col4a2*, *Tgfbr2*, *Usp9x*, *Yes1*, *Fut8*, *Appl1*, *Pard3*, *Sp1*, and *Zfyve9* were associated with the process of cellular response to TGF-β stimulus **(Fig. 6A, B)**. In contrast, upregulated genes, including *Bax*, *Cox7a1*, *Cox8a*, *Gabarap*, *Sharpin*, *Epm2a*, *Phka1*, *Ndufaf1*, *Tigar* and *Chchd2* were associated with mitochondrion organization, while multiple genes including *Bcl3*, *Col4a1*, *Col7a1*, *Mmp11*, *Sfrp2*, *Tnr*, *Ntn4*, *Mmp19*, *Pxdn*, and *Adamtsl4* were associated with ECM organization **(Fig. 6A, C)**. Similarly, DEG analysis followed by identification of the associated biological processes for MuSCs2 cluster showed that downregulated gene sets in *Xbp1^mKO^* mice, including *Cdk6*, *Fgf13*, *Sox5*, *Rock1*, and *Kif13a* were related to biological processes of cell division; *Hmga2*, *Smarcb1*, *Tead3*, *Arid4a*, *Cnot1*, and *Cnot2* associated with stem cell population maintenance; and *Cdh2*, *Cflar*, *Nck1*, *Numb1*, *Zeb2*, and *Tanc2* associated with cell projection organization **(Fig. 6D, E)**. On the contrary, upregulated genes were associated with electron transport chain (*mt-Co1*, *mt-Co2*, *mt-Co3*, *mt-Cytb*, and *mt-Nd1-5*), muscle contraction and actomyosin structure organization (*Acta1*, *Actc1*, *Myh3*, *Tnnc2*, *Tnnt3*, *Tpm2*, *Pgam2*, and *Tmem8c*) in MuSCs2 nuclei of *Xbp1^mKO^* mice compared to corresponding nuclei of *Xbp1^fl/fl^* mice **(Fig. 6D, F)**.

**Figure 6.**
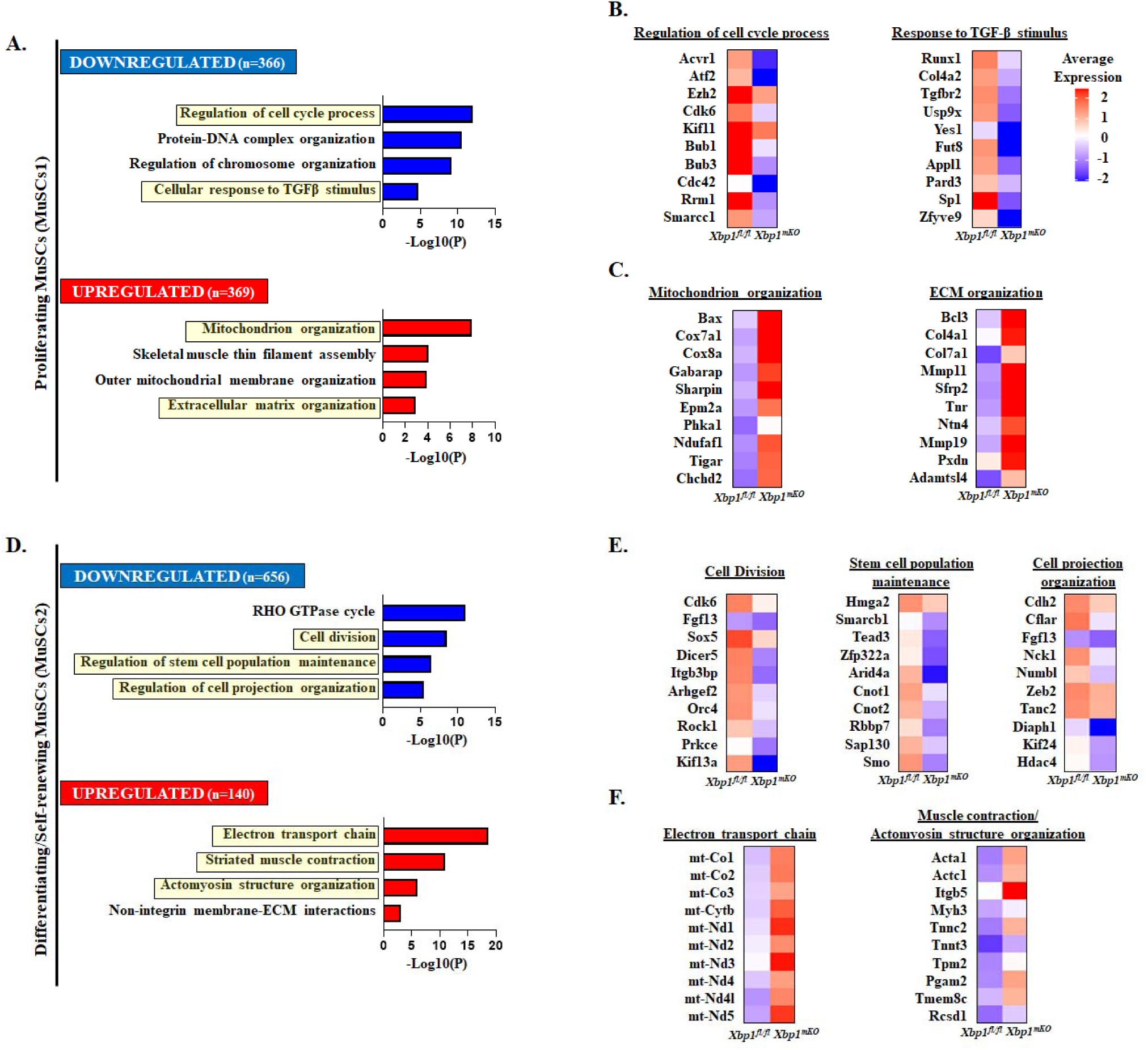
Myofiber XBP1 regulates transcriptomic profiles of satellite cells in regenerating muscle. Nuclei of satellite cell clusters (MuSCs1 and 2) in the snRNA-Seq analysis were used to identify differentially expressed genes followed by pathway enrichment analysis in *Xbp1^mKO^*mice compared to *Xbp1^fl/fl^* mice. **(A, D)** Bar graph showing enriched pathways associated with upregulated (red) and downregulated (blue) genes in MuSCs1 and MuSCs2 clusters respectively. Yellow boxes indicate pathways used for assessment of gene expression. Average gene expression of some of the **(B, E)** downregulated and **(C, F)** upregulated molecules involved in the enriched pathways.

To assess the potential signaling mechanisms affecting the abundance of satellite cells, we then combined the clusters of satellite cell nuclei (MuSCs1 and 2) and analyzed the DEGs by increasing the threshold of Log2FC ≥ |0.5| along with p-value < 0.05 in *Xbp1^mKO^*mice compared to control mice followed by pathway enrichment analysis. Results showed that overall upregulated genes in the clusters of *Xbp1^mKO^*satellite cells associated with the processes of muscle contraction, electron transport chain, signaling by TGF-β receptor complex, and regulation of muscle cell differentiation whereas downregulated genes were associated with RHO GTPase cycle, cell division, regulation of cytoskeleton organization, regulation of TOR signaling, and autophagy process **(Fig. 7A)**. Indeed, we observed multiple mitochondrial genes upregulated in the satellite cells of *Xbp1^mKO^*mice compared to *Xbp1^fl/fl^* mice as depicted by ridge plot analysis **(Fig. 7B)**. To further elucidate the mechanisms, we performed transcriptional regulator enrichment analysis in the combined satellite cell clusters in addition to the individual clusters MuSCs1 and 2, using the TRRUST database. Regardless of the combination of clusters, we observed that the upregulated genes were associated with the transcriptional regulators *Mef2c* and *Myod1* **(Fig. 7C)**. Moreover, a combination approach showed *Sp1* as another transcriptional regulator of the upregulated genes. While *Myod1* and *Mef2c* factors positively drive satellite cell differentiation and myogenesis process, *Sp1* transcription factor inhibits muscle cell differentiation (Vinals et al., 1997). Further analysis of the Myod1/Mef2c-regulated genes, including *Acta1*, *Atp2a1*, *Camk1d*, *Ckm*, *Col1a1*, *Myh1*, *Myh4*, *Myl1*, *Mylpf*, *Tnnc2* and *Tpm2* showed enriched expression in both the clusters of MuSCs **(Fig. 7D)**. Unlike the upregulated genes, the downregulated genes showed an association with different transcriptional regulators in MuSC1 (*Tal1*, *Notch1* and *Smad1*) and MuSC2 (*Smad6* and *Cux1*) clusters **(Fig. 7C)**. Interestingly, *Tal1*, *Notch1* and *Smad1* transcription factors play an important role in satellite cell activation, proliferation and differentiation and their inhibition leads to enhanced activation coupled with reduced proliferation and/or premature differentiation of satellite cells (Sousa-Victor et al., 2022).

**Figure 7.**
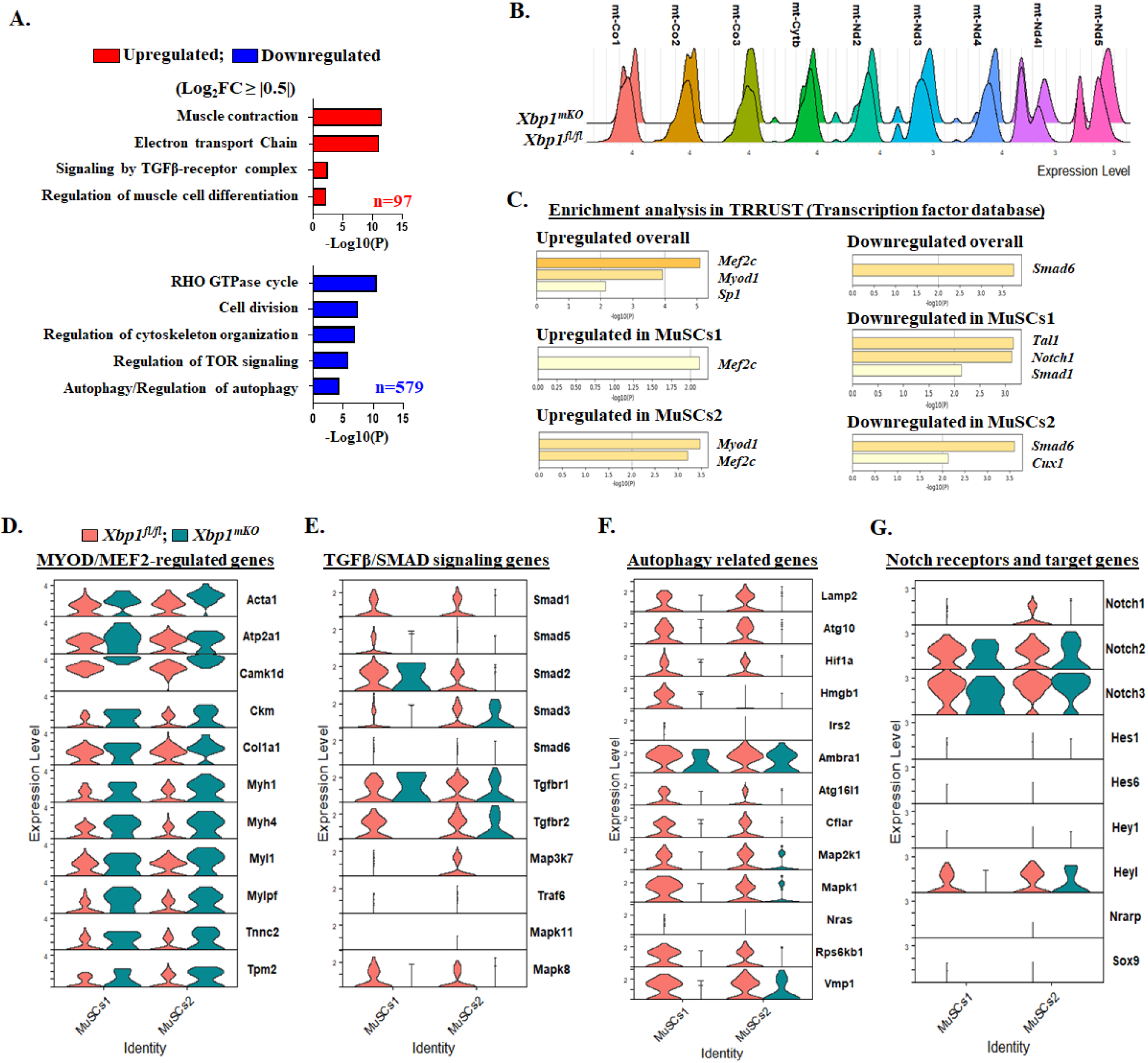
Myofiber XBP1 regulates distinct pathways in satellite cells during muscle regeneration. **(A)** Pathway enrichment analysis of differentially expressed genes across the combined cluster of satellite cell nuclei (MuSCs1 and MuSCs2). **(B)** Ridge plot showing gene expression of mitochondrial respiration associated molecules in satellite cells of *Xbp1^fl/fl^* and *Xbp1^mKO^* mice. **(C)** Enrichment of transcriptional regulators of upregulated and downregulated molecules in combined and individual clusters of satellite cell nuclei performed using TRRUST database. Violin plots for some of the deregulated genes **(D)** regulated by Myod1/Mef2c; and associated with **(E)** TGFβ/SMAD signaling pathway, **(F)** autophagy-lysosome pathway, and **(G)** Notch signaling pathway in nuclei of MuSCs1 and 2 clusters.

TGFβ signaling is another important mechanism that regulates satellite cell function during regenerative myogenesis. Canonical TGFβ signaling mediated by R-Smads and I-Smads leads to inhibition of satellite cell proliferation, impairment in differentiation and fusion, and eventually abrogation of muscle regeneration (Sousa-Victor et al., 2022). We have previously demonstrated that inducible deletion of TGFβ-activated kinase 1 (TAK1), an important signaling molecule of non-canonical TGFβ signaling, diminishes the self-renewal and proliferation of satellite cells during muscle regeneration following acute injury (Ogura et al., 2015). Our analysis showed the association of the downregulated genes in *Xbp1^mKO^* satellite cell nuclei with the regulatory factors Smad1 and Smad6. Further investigation of gene expression, as visualized through violin plots, showed reduced average expression of genes involved in the canonical TGF-β signaling (e.g. *Smad1*, *Smad5*, *Smad6*, *Tgfbr2*), non-canonical TGFβ signaling (e.g., *Map3k7*, *Traf6*, *Mapk11*, *Mapk8*), autophagy-related molecules (e.g., *Lamp2*, *Atg10*, *Hif1a*, *Hmgb1*, *Irs2*, *Ambra1*, *Atg16l1*, *Cflar*, *Map2k1*, *Mapk1*, *Nras*, *Rps6kb1*, and *Vmp1*), and Notch receptor (e.g. *Notch1*, but not *Notch2* or *Notch3*) and Notch target genes (e.g., *Hes1*, *Hes6*, *Hey1*, *Heyl*, *Nrarp*, and *Sox9*) in the satellite cell nuclei of *Xbp1^mKO^* mice compared to *Xbp1^fl/fl^* mice **(Fig. 7E-G)**.

These results suggest that deletion of XBP1 in myofibers reduces satellite cell abundance and myogenic function during muscle regeneration potentially through regulating TGFβ and Notch signaling and blunting autophagy in a cell non-autonomous manner.

### XBP1 regulates the abundance of non-myogenic cells

We next sought to determine whether ablation of XBP1 in myofibers also affects the transcriptional profiles of non-muscle cells. Initial analysis of the proportion of non-muscle nuclei showed a decrease in FAP2 (7.19% vs. 11.30%), and Macro (7.19% vs. 11.90%) nuclei, whereas proportion of nuclei in FAP1 (69.46% vs. 64.46%) and Endo (16.17% vs. 12.35%) cluster showed an increase in *Xbp1^mKO^* mice compared to *Xbp1^fl/fl^* mice **(Fig. 8A)**. Next, we performed DEG analysis followed by pathway enrichment to identify heterogeneity in biological processes regulated by XBP1. FAPs are a major source of various growth factors and components of ECM, both of which aid in muscle regeneration (Sousa-Victor et al., 2022). Our analysis of combined FAP clusters (FAP1 and FAP2) showed that 57 genes associated with electron transport chain, regulation of muscle system process, and carbohydrate catabolic process were upregulated, whereas 1165 genes associated with chromatin organization, cell migration, IL-6 signaling pathway, skeletal system development, and mechanisms associated with pluripotency were downregulated **(Fig. 8B)**. Endothelial cells (ECs) also play a critical role in muscle regeneration, especially affecting satellite cell function.

**Figure 8.**
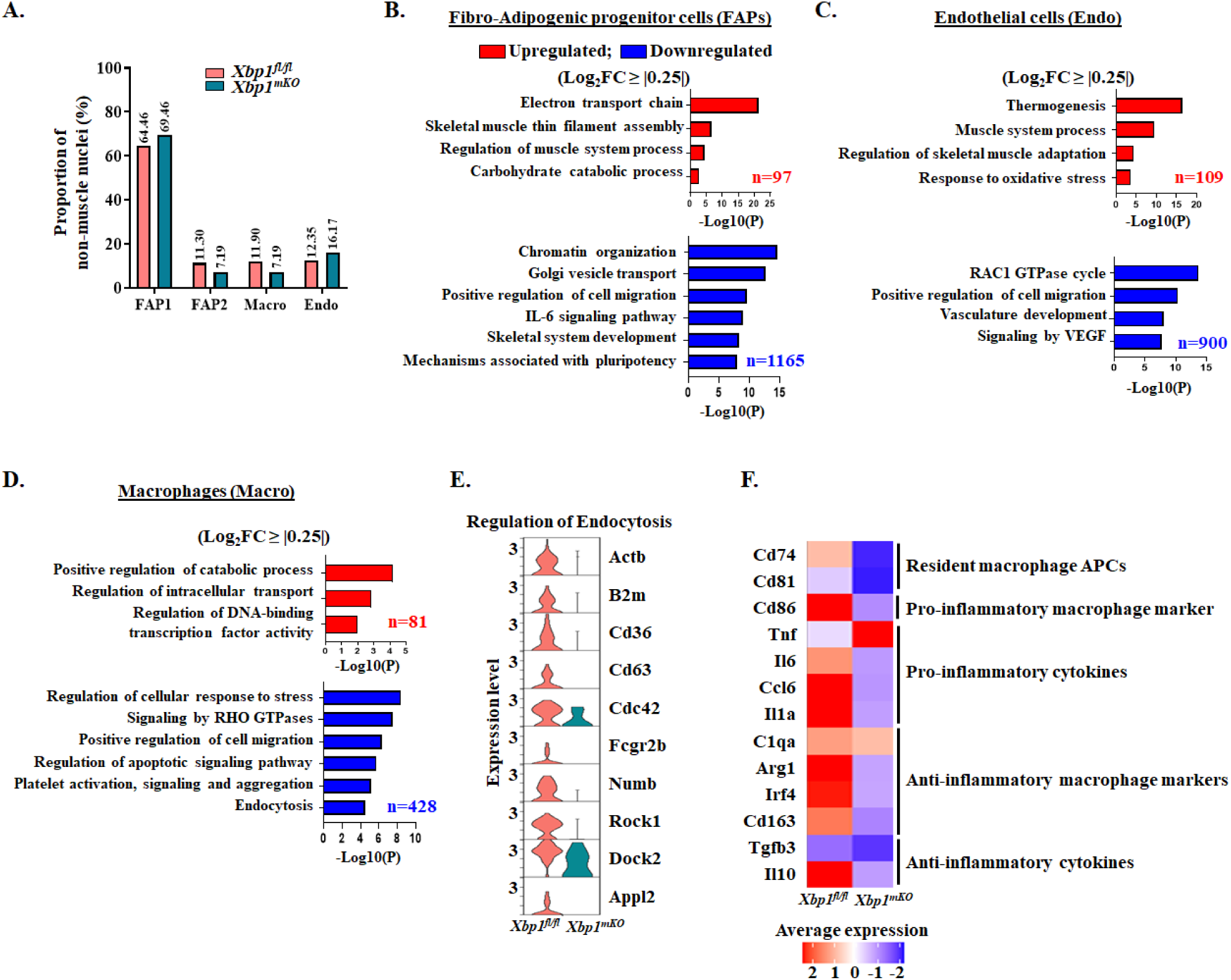
XBP1 in myofibers regulates the abundance of non-myogenic cells during regenerative myogenesis. **(A)** Proportion of non-muscle cell nuclei in injured TA muscle of *Xbp1^fl/fl^* and *Xbp1^mKO^*mice. Differential gene expression analysis was performed in nuclei of fibro-adipogenic progenitor cells (FAPs), endothelial cells (Endo), and macrophages (Macro) clusters. Pathway enrichment analysis of deregulated genes in **(B)** FAPs, **(C)** Endo, and **(D)** Macro clusters of *Xbp1^mKO^* mice compared to *Xbp1^fl/fl^* mice. **(E)** Violin plot showing gene expression levels of molecules related to regulation of endocytosis. **(F)** Heatmap showing average gene expression of cell surface markers and cytokines secreted by macrophages.

Multiple mechanisms, including ECs-mediated angiogenesis and secretion of growth factors (e.g., IGF1, HGF, bFGF, VEGF etc.), Notch ligand Dll4, and chemo-attractants positively regulate satellite cell proliferation and self-renewal and skeletal muscle repair (Sousa-Victor et al., 2022). Analysis of nuclei in Endo cluster of 5d-injured TA muscle of *Xbp1^mKO^* mice compared to *Xbp1^fl/fl^*mice showed that 109 genes were upregulated, whereas 900 genes were downregulated. The upregulated genes were associated with thermogenesis, muscle system process, regulation of skeletal muscle adaptation, and response to oxidative stress. By contrast, downregulated genes were associated with RAC1 GTPase cycle, positive regulation of cell migration, vasculature development, and signaling by VEGF **(Fig. 8C)**.

Macrophages are another important cell type that regulates muscle regeneration. Macrophages interact with myogenic cells and promote muscle regeneration by exerting immune and non-immune functions (Tidball, 2017). We investigated whether targeted deletion of XBP1 affects macrophage function in regenerating skeletal muscle. Our analysis showed 509 deregulated genes, out of which 81 genes were upregulated and 428 genes were downregulated in nuclei of macrophage cluster of *Xbp1^mKO^* mice compared to *Xbp1^fl/fl^*mice. Subsequent pathway enrichment analysis of the identified DEGs showed that upregulated genes were associated with positive regulation of catabolic process, regulation of intracellular transport, and negative regulation of DNA-binding transcription factor activity. In contrast, downregulated genes were associated with regulation of cellular response to stress, signaling by RHO GTPases, regulation of cell migration, regulation of apoptotic signaling pathway, platelet activation, signaling and aggregation, and endocytosis **(Fig. 8D)**. Endocytosis by macrophages (i.e., phagocytosis) is an important process that helps in clearing the damaged cells/cell debris following muscle injury (Tidball, 2017; Tidball et al., 2014). Gene expression of multiple molecules regulating endocytosis, such as *Actb*, *B2m*, *Cd36*, *Cd63*, *Cdc42*, *Fcgr2b*, *Numb*, *Rock1*, *Dock2*, and *Appl2* was downregulated in macrophages of 5d-injured TA muscle of *Xbp1^mKO^* mice compared to control mice **(Fig. 8E)**. Macrophages are also a major source for many cytokines and macrophage polarization from proinflammatory to anti-inflammatory phenotype is essential for timely and efficient muscle regeneration (Tidball, 2017; Tidball et al., 2014). We investigated the expression of various markers of macrophages and cytokines secreted by pro- and anti-inflammatory macrophages. Intriguingly, the markers of resident macrophage antigen presenting cells (*Cd74* and *Cd81*), pro-inflammatory macrophage (*Cd86*), and cytokines (*Il6*, *Ccl6* and *Il1a*) were drastically reduced in *Xbp1^mKO^* compared to *Xbp1^fl/fl^* mice. By contrast, we observed that gene expression of pro-inflammatory cytokine TNFα was induced in macrophages of *Xbp1^mko^* mice **(Fig. 8F)**. Surprisingly, the expression of anti-inflammatory macrophage markers (*C1qa*, *Arg1*, *Irf4* and *Cd163*) and cytokines (*Tgfb3*, *Il10*) were also reduced **(Fig. 8F)** suggesting that ablation of XBP1 disrupts overall macrophage activation and function contributing to the impairment of muscle regeneration.

## DISCUSSION

Skeletal muscle injury is a common manifestation of direct trauma such as muscle lacerations and contusions, indirect insults such as strains, and muscle degenerative diseases such as muscular dystrophies and inflammatory myopathies. However, the molecular and signaling mechanisms involved in muscle regeneration have not yet been completely elucidated. Accumulating evidence suggests that the UPR plays important roles in the regulation of muscle formation and regeneration (Afroze and Kumar, 2019; Bohnert et al., 2018). Earlier studies showed that the ATF6 arm of the UPR is activated during muscle development where it mediates apoptosis of a subpopulation of myoblasts that may be incompetent of handling cellular stresses (Nakanishi et al., 2005). The UPR may have a role in fine-tuning myogenic differentiation. Activation of the PERK arms of the UPR and levels of CHOP transiently increase during myogenic differentiation (Alter and Bengal, 2011). Recent studies also suggest that the PERK/eIF2α signaling is essential for self-renewal of satellite cells in skeletal muscle of adult mice (Xiong et al., 2017; Zismanov et al., 2016). In the present study, we first employed published scRNA-seq dataset to understand how the activation of the UPR and associated processes are affected at different time points after muscle injury. Our results demonstrate that the UPR, ERAD, and EOR are activated not only in myogenic cells, but also in other cell types, including macrophages, lymphocytes, endothelial cells, and FAPs present in the muscle microenvironment. Furthermore, gene expression of XBP1 and its known targets Dnajb9, Derl1, Atf4 and Calr was also induced in mature muscle cells at day 2 and 5 post injury (**Fig. 1**) suggesting that the XBP1-mediated UPR and ERAD play important roles for the regeneration of injured myofibers.

We recently reported that targeted deletion of IRE1α or XBP1 in differentiated myofibers inhibits skeletal muscle regeneration in adult mice (Roy et al., 2021). However, the mechanisms by which the IRE1α/XBP1 signaling in myofiber regulates muscle regeneration remained largely unknown. Because snRNA-seq is the preferred approach to understand the transcriptional heterogeneity in multinucleated cells, including myofibers, we performed snRNA-seq on 5d-injured TA muscle of control and muscle-specific *Xbp1*-knockout mice. Our analysis of snRNA-seq data confirmed the presence of nuclear clusters of myofibers, myoblasts, satellite cells, and non-muscle cells at day 5-post injury (**Fig. 2**). Importantly, our snRNA-seq experiments (**Fig. 3B**) and immunohistochemical analysis of muscle tissues (**Fig. 5**) confirmed our previously published findings (Roy et al., 2021) that myofiber-specific ablation of XBP1 significantly reduces the proportion of satellite cell myonuclei in regenerating muscle. Our snRNA-seq analysis and follow-up experiments using cultured EDL myofibers (**Fig. S11, 12**) further suggests that myofiber-specific ablation of XBP1 inhibits the self-renewal and proliferation and augments precocious differentiation of satellite cells which may be important mechanisms for the reduction of muscle regeneration in *Xbp1^mKO^*mice.

During ER stress, spliced XBP1 transcription factor augments the ERAD pathway, which involves recognition of misfolded proteins, their translocation into the cytosol followed by proteolysis through ubiquitination-proteasomal system or autophagy-lysosomal degradation (Wu and Rapoport, 2018). It is now increasingly evidenced that autophagy plays a key role in tissue regeneration and the maintenance of cellular homeostasis. While most studies have investigated the role of autophagy in satellite cells during muscle regeneration, it is noteworthy that autophagy in mature myofibers is critical for the degradation of necrotic myofibers following muscle damage. This can potentially affect the secretome-mediated satellite cell dynamics and the recruitment of non-muscle supporting cells during regenerative myogenesis (Call and Nichenko, 2020; Chen et al., 2022). Interestingly, XBP1 mediates autophagy directly by transcriptional regulation of Beclin1 (Margariti et al., 2013), a protein indispensable for the formation of autophagosomes, or indirectly through its interaction with FoxO1 protein (Kishino et al., 2017) or transcriptional regulation of Transcription factor EB (Zhang et al., 2021). Our snRNA-seq analysis and knockdown studies in cultured myotubes demonstrate that XBP1 controls the gene expression of components of the ubiquitin-proteasome system and autophagy (**Fig. 3, 4**). Therefore, it is likely that the inhibition of these proteolytic systems in *Xbp1*-deficient myofibers reduces the ERAD response resulting in persistent ER stress and perturbation in the muscle regeneration. Although the physiological significance remains unknown, we have also observed genetic ablation of XBP1 increases the gene expression of components of mitochondrial electron transport chain and oxidative phosphorylation (**Fig. 3, 4**). Interestingly, our experiments suggested that the levels of OXPHOS proteins are increased in regenerating or developing muscle, but not in uninjured muscle of adult *Xbp1^mKO^* mice (**Fig. S8**). It is possible that the increased expression of mitochondrial genes may be a compensatory response to augment muscle regeneration and improve functional mitochondria due to the inability of clearance of dysfunctional mitochondria through autophagy.

Consistent with our previously published report (Roy et al., 2021), trajectory analysis of myogenic lineage and analysis of transcriptome in nuclei of muscle cells showed that the formation or repair of new myofibers was significantly reduced in skeletal muscle of muscle-specific *Xbp1*-knockout mice compared with control mice **(Fig. 4** and **5)**. It is apparent that myofiber XBP1 regulates muscle formation in both cell autonomous and -non-autonomous manner. Deletion of XBP1 reduces the abundance of satellite cells and their progression into myogenic lineage resulting in reduced formation of eMyHC^+^ myofibers. Furthermore, our analysis of myonuclei suggests precocious differentiation of muscle progenitor cells, which impairs normal progression of muscle repair **(Fig. 5)**. Indeed, analysis of clusters of satellite cell nuclei suggested that the gene expression of molecules involved in stem cell population maintenance and cell division processes are diminished in myofiber *Xbp1*-knockout group.

Interestingly, we also observed that in addition to myofibers and myoblasts, the gene expression of molecules involved in mitochondrial organization and electron transport chain is perturbed in satellite cell nuclei of *Xbp1*-knockout mice **(Fig. 6)**. It is notable that healthy mitochondria are essential for the regenerative capacity of satellite cells. The loss of mitochondrial dynamics in various conditions such as aging or genetic muscle disorders deregulates the mitochondrial electron transport chain (ETC), leading to inefficient oxidative phosphorylation metabolism and mitophagy and increased oxidative stress (Sousa-Victor et al., 2022). While the exact mechanisms remain unknown, it is possible that loss of XBP1 results in deregulation of production of various growth factors and inflammatory cytokines, which affect mitochondrial dynamics in a paracrine manner. In addition, transient upregulation of autophagy is also critical for the proliferation, self-renewal, and differentiation of satellite cells (Sousa-Victor et al., 2022). Like myofibers, we also observed reduction in multiple autophagy-related molecules, including *Atg10*, *Hif1a*, *Hmgb1* and *Atg16l1*, which could also potentially regulate mitochondrial dynamics in satellite cells (**Fig. 7F**).

Another potential mechanism for impairment of muscle repair in *Xbp1^mKO^* mice is the disruption of various signaling pathways in satellite cells. Our analysis of satellite cell nuclei revealed the disruption of both canonical and non-canonical TGFβ signaling in *Xbp1*-knockout group. Previous studies from our group have shown that MAP3K7 (also known as TAK1), a component of non-canonical TGFβ signaling is essential for the self-renewal and proliferation of satellite cells and inducible deletion or pharmacological inhibition of TAK1 causes precocious differentiation of satellite cells (Ogura et al., 2015). Interestingly, gene expression of *Map3k7* was strongly diminished in the satellite cell nuclei of *Xbp1^mKO^* mice (**Fig. 7E**). Notch signaling also plays a critical role in self-renewal and maintenance of satellite cell pool in skeletal muscle (Bjornson et al., 2012; Gioftsidi et al., 2022). Our studies have shown that genetic ablation of XBP1 inhibits the gene expression of Notch receptors and Notch target genes in satellite cell nuclei (**Fig. 7G**). Indeed, inhibition of Notch signaling in satellite cells could be an important mechanism for reduced self-renewal capacity of myofiber-associated satellite cells in *Xbp1^mKO^* cultures (**Fig. S12**). These results are consistent with our previous findings which also demonstrated that myofiber-specific deletion of IRE1 (an upstream activator of XBP1) inhibits Notch signaling in regenerating skeletal muscle of adult mice (Roy et al., 2021).

In addition to satellite cells, muscle regeneration also involves participation of several other cell types, such as FAPs, endothelial cells, and pro- and anti-inflammatory macrophages (Tidball, 2017; Tidball et al., 2014). Interestingly, our analysis showed that overall proportion of non-muscle cell nuclei was considerably reduced in *Xbp1^mKO^* compared to *Xbp1^fl/fl^*group. Muscle repair involves sequential activation of pro-inflammatory and anti-inflammatory macrophages. M1 macrophages are pro-inflammatory and dominate the necrotic phase. In contrast, alternatively activated M2 macrophages are anti-inflammatory which prevail during the regenerative stage to facilitate myofiber repair (Tidball, 2017; Tidball et al., 2014). Indeed, transition from M1 to M2c macrophage phenotype is critical for skeletal muscle regeneration (Wang et al., 2014). The abundance of M1 and M2c macrophages is regulated by various factors, including proinflammatory and anti-inflammatory cytokines (Tidball, 2017). Intriguingly, our experiments showed that there was about 50% reduction in the proportion of macrophage nuclei and diminished gene expression of molecules involved in phagocytosis in *Xbp1^mKO^* group (**Fig. 8A, E**). Further analysis showed that markers of both proinflammatory and anti-inflammatory macrophages as well as cytokines (except TNF, which showed increased expression) were reduced in the macrophage nuclei of *Xbp1^mKO^* mice **(Fig. 8F)**. These finding suggest that XBP1 is essential for maintenance and timely transition of proinflammatory macrophages into anti-inflammatory macrophages and impairment in macrophage recruitment could be another mechanism for reduction in the muscle regeneration in *Xbp1^mKO^* mice. In summary, our snRNA-seq analysis has identified the role and molecular network through which myofiber XBP1 regulates skeletal muscle regeneration in adult mice.

### Limitation of the study

Our study also has a few drawbacks. For example, we have performed our experiments at only one time point after muscle injury. It would be important to understand whether XBP1 exerts similar effects at different stages of muscle regeneration. The number of sequenced nuclei in the *Xbp1*-knockout group is limited, which may have some effect on the coverage of all cell types or subtypes during muscle regeneration and the trajectory analysis. As is the case for most of the approaches, snRNA-seq measures the mRNA levels only in nuclei of the cell, but not in cytoplasm. To capture the changes in whole transcriptome, it may be more useful to perform scRNA-seq and snRNA-seq experiments in parallel. Furthermore, many results such as changes in mitochondrial content should be validated by additional biochemical and histochemical approaches. The composition of non-myogenic cells in regenerating muscle of control and *Xbp1*-knockout mice should also be validated using cell biology approaches, such as flow cytometry. Finally, it would be interesting to determine how the components of UPR are affected in skeletal muscle of animal models of various muscle disorders.

## STAR Methods

### Key resources table

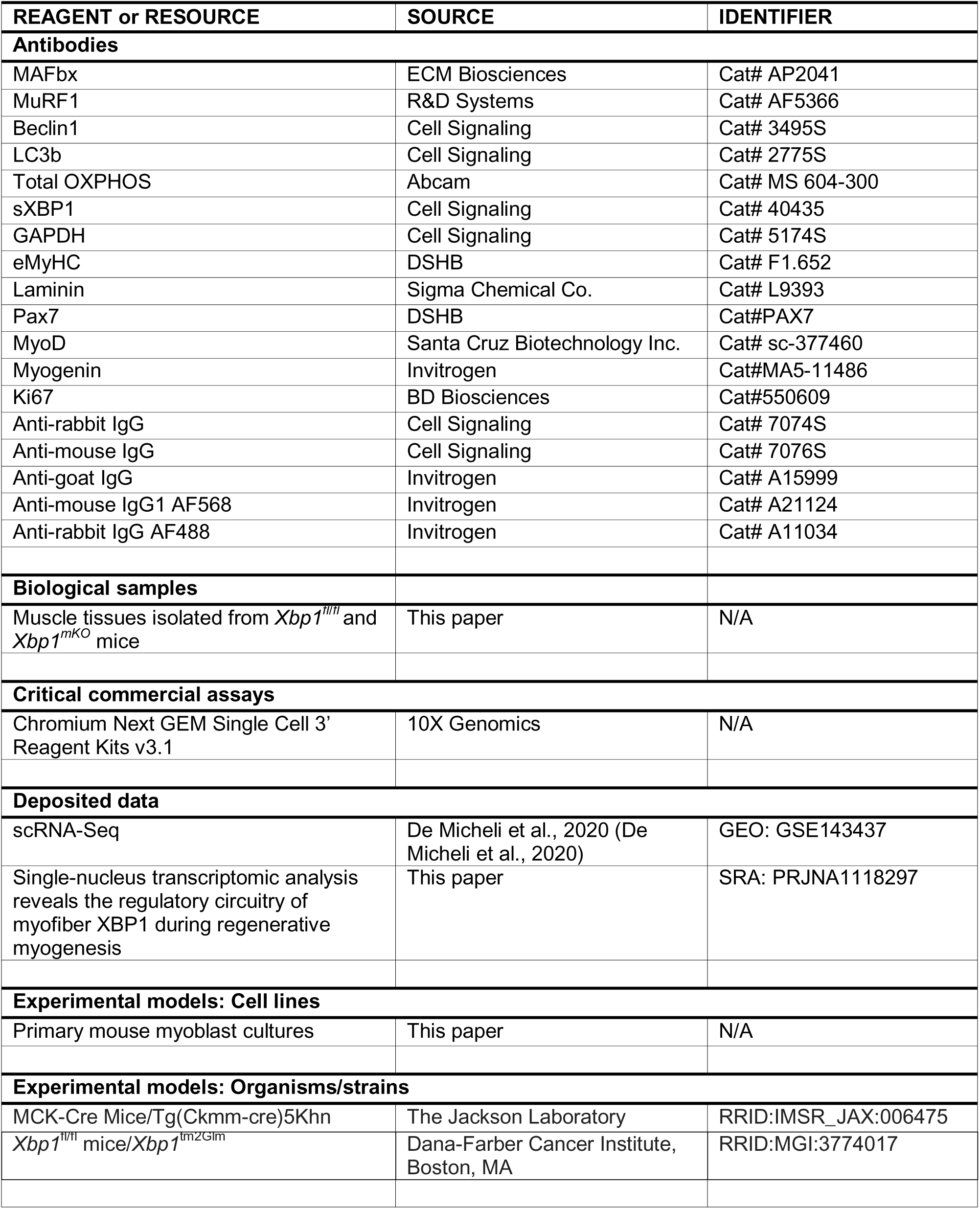
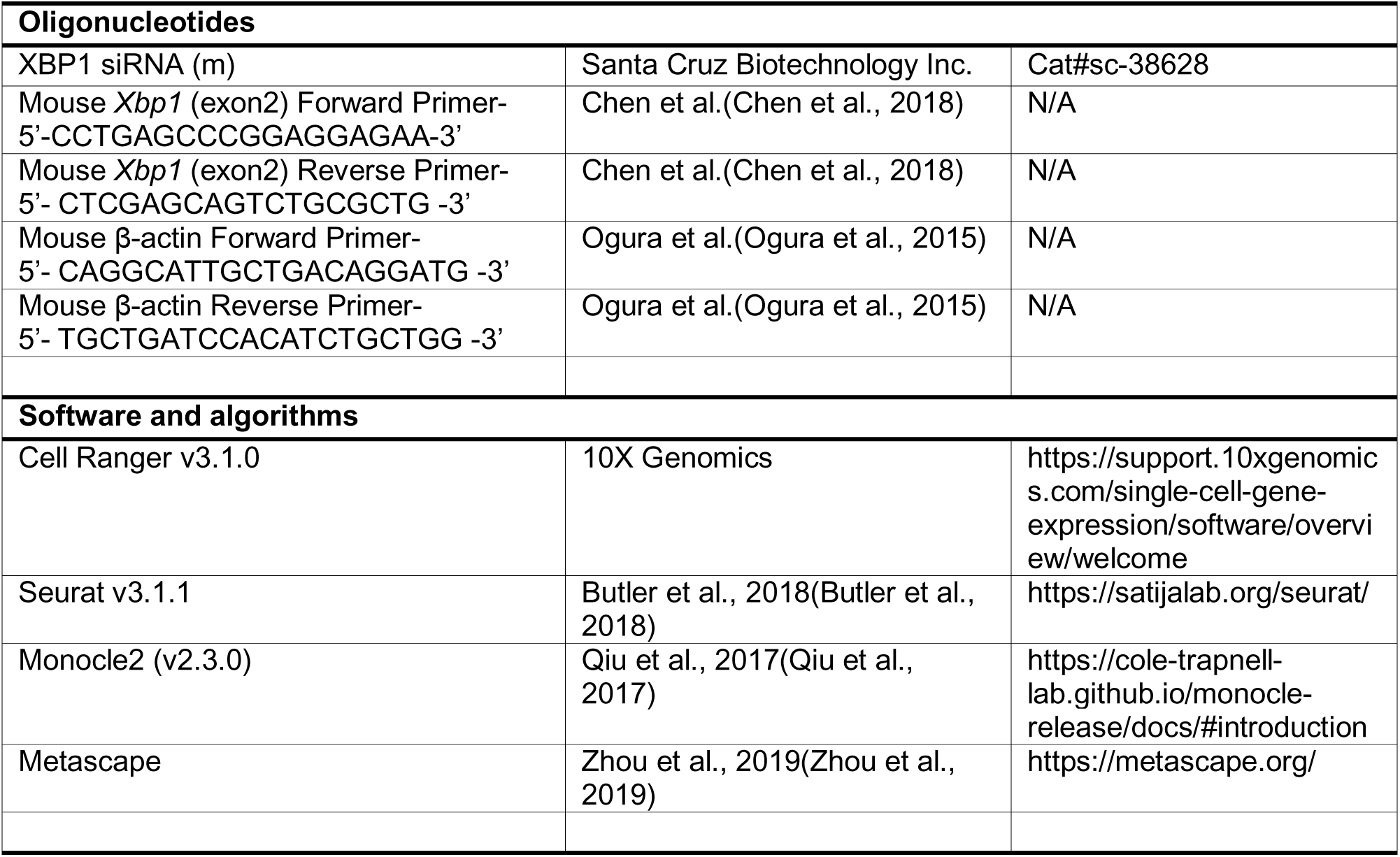

### Resource availability

Further information and requests for resources and reagents should be directed to and will be fulfilled by the lead contact, Ashok Kumar

### Materials availability

This study did not generate any novel reagents.

### Data availability

- All the raw data files for snRNA-seq experiment can be found on NCBI SRA repository using the accession code PRJNA1118297.
- This paper does not produce original codes.
- Any additional information required to reanalyze the data reported in this paper is available from lead contact upon request.

### Experimental model

#### Animals

Floxed *Xbp1* (*Xbp1^fl/fl^*) mice were crossed with MCK-Cre (Strain: B6.FVB(129S4)-Tg(Ckmm-cre)5Khn/J, Jackson Laboratory, Bar Harbor, ME) mice to generate muscle-specific XBP1 knockout (*Xbp1*^mKO^) and littermate control (Xbp1^fl/fl^ mice) as described (Bohnert et al., 2019; Roy et al., 2021). All mice were in the C57BL/6 background, and their genotype was determined by PCR from tail DNA. Both male and female mice were used in this study. TA muscle of adult mice was injected 50 µl of 1.2% BaCl_2_ (Sigma Chemical Co.) dissolved in saline to induce necrotic injury as described (Hindi and Kumar, 2016). The mice were euthanized at day 5 or 21 after injury and the TA muscle was collected and processed for histological analysis or snRNA-seq. All the experiments were performed in strict accordance with the recommendations in the Guide for the Care and Use of Laboratory Animals of the National Institutes of Health. All the animals were handled according to approved institutional animal care and use committee protocol (PROTO201900043) of the University of Houston. All surgeries were performed under anaesthesia, and every effort was made to minimize suffering.

### Method details

#### Histology, immunohistochemistry, and morphometric analysis

Uninjured, 5d- or 21d-injured TA muscle of mice was isolated and frozen in liquid nitrogen and sectioned in a microtome cryostat. For the assessment of muscle morphology, 8-mm thick transverse sections of TA muscle were stained with Hematoxylin and Eosin (H&E). The sections were examined under Nikon Eclipse TE 2000-U microscope (Nikon). For immunohistochemistry study, frozen TA muscle sections were fixed in acetone or 4% paraformaldehyde (PFA) in PBS, blocked in 1% bovine serum albumin in PBS for 1 h and incubated with anti-eMyHC (1:200, DSHB, University of Iowa, Iowa City, IA) or anti-Pax7 (1:100, DSHB, University of Iowa, Iowa City, IA) and anti-laminin (1:500, Sigma) in blocking solution at 4°C overnight under humidified conditions. The sections were washed briefly with PBS before incubating with Alexa Fluor 488- or 546-conjugated secondary antibody (1:1500, Invitrogen) for 1 h at room temperature and then washed three times for 5 min with PBS. DAPI was used to counterstain nuclei. The slides were mounted using fluorescence medium (Vector Laboratories) and visualized at room temperature on Nikon Eclipse Ti-2E Inverted Microscope (Nikon), a digital camera (Digital Sight DS-Fi3, Nikon), and Nikon NIS Elements AR software (Nikon). Image levels were equally adjusted using Adobe Photoshop CS6 software (Adobe). For quantitative analysis, CSA of eMyHC^+^ myofibers and number of Pax7^+^ cells in TA muscle were analysed. For each muscle, the distribution of myofiber CSA was calculated by analysing ∼200 myofibers.

#### Isolation, culture, and staining of single myofibers

Single myofibers were isolated from EDL muscle of mice after digestion with collagenase II (Sigma-Aldrich) and trituration as described (Gallot et al., 2016). Suspended fibers were cultured in 60-mm horse serum-coated plates in DMEM supplemented with 10% FBS (Invitrogen), 2% chicken embryo extract (Accurate Chemical and Scientific Corporation), 10 ng/ml basis fibroblast growth factor (PeproTech), and 1% penicillin-streptomycin for 3 days. Freshly isolated fibers (0 h) and cultured fibers (48 or 72 h) were then fixed in 4% PFA and stained with anti-Pax7 (1:100, Developmental Studies Hybridoma Bank [DSHB]), MyoD (1:200, sc-377460, Santa Cruz Biotechnology Inc.), and anti-Ki67 (1:200, BD Biosciences).

#### Primary myoblast cultures and transfections

Primary myoblasts were isolated from hind limb muscle of wild-type mice as described (Hindi et al., 2017). For myotube formation, primary myoblasts were incubated in differentiation medium (DMEM, 2% Horse serum) for 48 h.

Myotube cultures were transfected with control or XBP1 siRNA (SantaCruz Biotechnology) using Lipofectamine RNAiMAX Transfection Reagent (ThermoFisher Scientific) and collected 24 h later for protein extraction.

#### Satellite cell isolation

Satellite cells were isolated from hindlimb muscles of *Xbp1^fl/fl^*and *Xbp1^mKO^* mice using Satellite Cell Isolation Kit, mouse (Miltenyi Biotec). Briefly, hindlimb muscles were isolated, washed in PBS, minced into coarse slurry and enzymatically digested at 37 °C for 1 h by adding 400 U/ml collagenase II (Gibco, Life Technologies). The digested slurry was spun, pelleted and triturated several times and then passed through a 70-μm and then 30-μm cell strainer (BD Falcon). The filtrate was spun at 1,000g and the cell pellets were used for satellite cell isolation using the manufacturer’s protocol.

#### Western blot

Frozen TA and GA muscles of *Xbp1^fl/fl^* and *Xbp1^mKO^* mice, and cultured myotubes were homogenized in lysis buffer (50 mM Tris-Cl (pH 8.0), 200 mM NaCl, 50 mM NaF, 1 mM dithiothreitol, 1 mM sodium orthovanadate, 0.3% IGEPAL and protease inhibitors).

Approximately, 100μg of protein was resolved on each lane on 10% SDS–polyacrylamide gel electrophoresis, electro-transferred onto nitrocellulose membrane, probed using specific antibody and detected by chemiluminescence. Uncropped Western blot images are presented in supplemental **Fig. S13**.

#### Analysis of single-cell RNA sequencing dataset

Using the publicly available dataset GSE143437, raw data was processed using Seurat package on R software, as described for snRNA-Seq analysis. Integrated Seurat object containing transcriptomic profiles of muscle tissue at day 0, 2, 5 and 7 post-injury was used to analyse the expression levels of genes related to ER stress-UPR, ERAD and ER stress-overload response. Expression levels and percentage of cells expressing the genes were visualized by dot plots.

#### SnRNA-seq and data processing

Nuclei were isolated from freshly isolated 5d-injured TA muscle of mice using 10X Genomics Chromium nuclei isolation kit following a protocol suggested by the manufacturers (10X Genomics, Pleasanton, CA). The nuclei were pooled from 3-4 mice in each group. snRNA-seq libraries were generated using Chromium Next GEM Single Cell 3′ Gene Expression v3.1 kit (10X Genomics) according to the manufacturers’ protocol.

Sequencing was performed on an Illumina NextSeq 2000 system with the pair-end sequencing settings Read1 – 28 bp, i7 index – 10bp, i5 index – 10 bp and Read2 – 90 bp. Estimated number of cells sequenced were ∼2800 for *Xbp1^fl/fl^*and ∼2100 for *Xbp1^mKO^* groups. The CellRanger Software Suite (10X Genomics, v3.1.0) was used for data demultiplexing, transcriptome alignment, and UMI counting. Raw base call files were demultiplexed using cellranger mkfastq. The mouse genome, refdata-cellranger-mm10-1.2.0 from the 10x Genomics support website, was used as reference for read alignments and gene counting with cellranger count. ∼60,000 mean reads per cell were obtained and ∼19,000 total genes were detected in each sample. Elimination of RNA background was performed using SoupX R package.

#### Bioinformatics analysis

The downstream analysis of pre-processed data was performed on R software (v4.2.2), using the Seurat package (v4.3.0). Firstly, the nuclei for each object (*Xbp1^fl/fl^* and *Xbp1^mKO^*) were filtered based on unique feature counts, with the exclusion criteria of nuclei containing less than 500 and more than 20,000 unique features. The filtered nuclei were further refined by identifying and excluding doublets using the DoubletFinder package and filtering the nuclei with percent mitochondrial counts less than 5%. Nuclei from each Seurat object (*Xbp1^fl/fl^* and *Xbp1^mKO^*) were separately normalized using SCTransform function and subjected to Principal Component Analysis using RunPCA function. Nuclei that expressed at least 500 and a maximum of 20,000 genes (nFeature_RNA) were selected for analysis. FindNeighbors and FindClusters functions were used for clustering nuclei and visualized using RunUMAP function (Uniform Manifold Approximation and Projection). To eliminate the batch effects, the individually processed Seurat objects were integrated using the SCTIntegration workflow. Cell type identification of nuclei in unbiasedly obtained clusters was performed by analyzing the expression of previously described gene markers. In addition, differentially expressed genes (Log2FC (fold change) ≥ 1 and p-value <0.05) were identified across all clusters and the biological processes associated with enriched genes in each cluster were used for further validation of cell types.

Differential gene expression analysis between corresponding clusters of *Xbp1^fl/fl^* and *Xbp1^mKO^* nuclei was performed using the FindMarkers function with absolute Log2FC ≥ |0.5| (or |0.25| to increase the range) and p-value < 0.05 considered as significant. Pathway enrichment analysis of all differentially expressed genes was performed using Metascape tool (metascape.org). Genes of interest were investigated for their expression patterns using heatmaps, violin plots, feature plots, and ridge plots. Furthermore, identification of transcriptional regulators was performed using TRRUST database, visualized on Metascape analysis platform. The Monocle2 (v2.3.0) package was utilized to perform pseudotime analysis and trajectory mapping through reversed graph embedding, followed by differential gene expression analysis and generation of heatmaps across pseudotime.

#### Statistical Analysis

Results are represented as mean ± SEM unless indicated otherwise. Unpaired Student’s *t* test or two-way analysis of variance (ANOVA) followed by Tukey’s multiple comparison test was performed to analyse the data. A value of p ≤ 0.05 was considered significant.

## Acknowledgements

We thank Dr. Laurie Glimcher of Dana Farber Cancer Institute for providing floxed XBP1 mice. This work was supported by the National Institute of Health grants AR081487 and AR059810 to AK. The graphical abstract was created in BioRender.com.

## Authors’ contribution

A.K., R.D., and Y.L. designed the work. A.S.J. wrote the manuscript and all authors edited the manuscript. A.S.J., M.B.C., and M.T.S., performed the experiments. P.H.G. helped with genomics experiments and bioinformatics analysis of RNA-Seq experiments. The snRNA-seq was performed in the University of Houston Sequencing Core (UH-SEQ).

## Supplemental Figures S1-S13 and Legends

**Figure S1.**
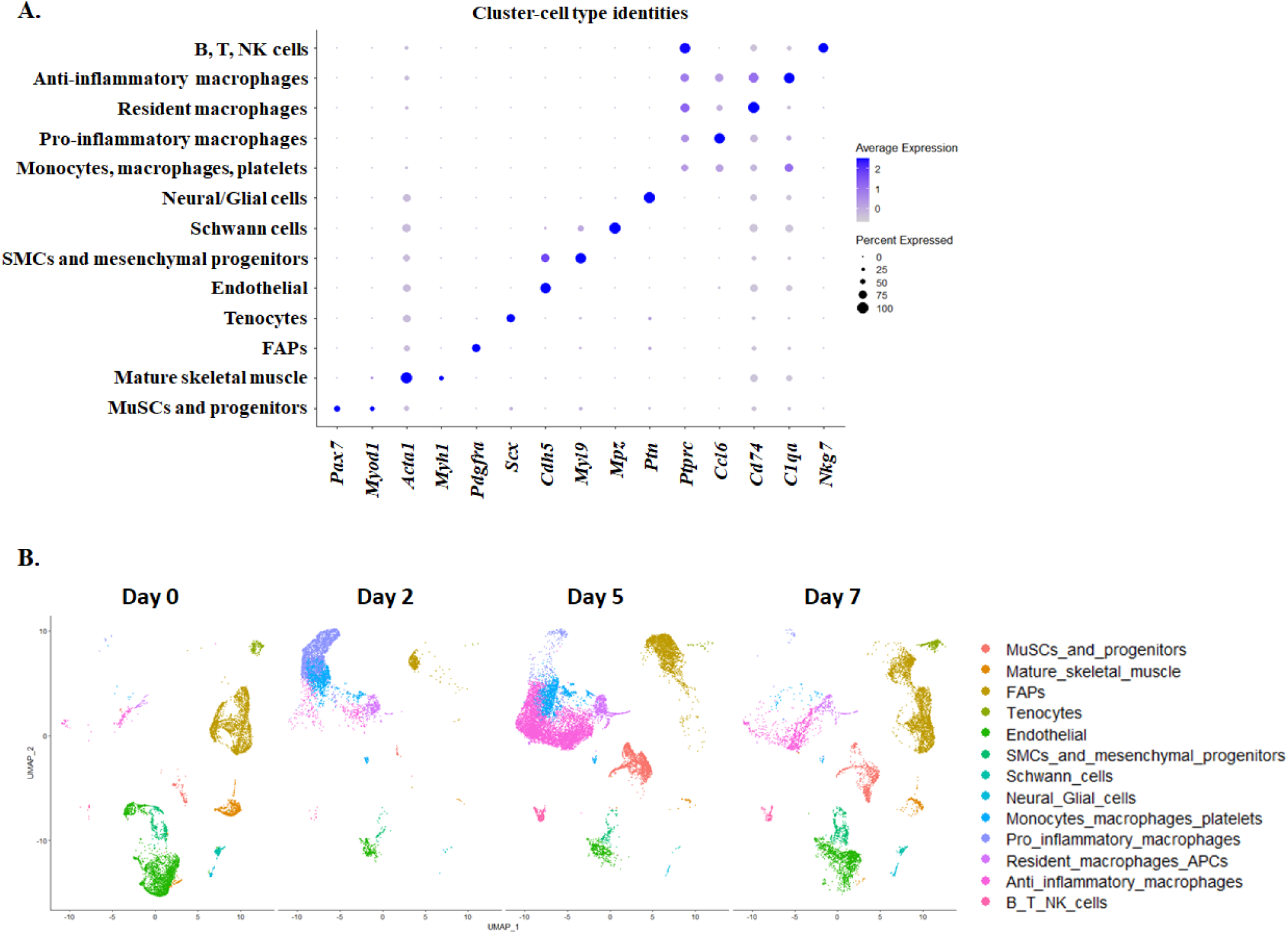
Analysis of scRNA-seq dataset identifies distinct cell types during muscle regeneration. The scRNA-seq dataset (GSE143437), consisting of regenerating muscle tissue samples at day 0, 2, 5 and 7 post injury, was analyzed using R software (v4.2.2). Gene expression of various markers of cellular identification were analyzed to annotate distinct cell types of the spatially distributed clusters. **(A)** Dot plot shows average expression and percentage of cells expressing the indicated genes in different cell types. **(B)** Split-UMAPs visually represent the changes in proportion of various cell types at different time points of muscle regeneration.

**Figure S2.**
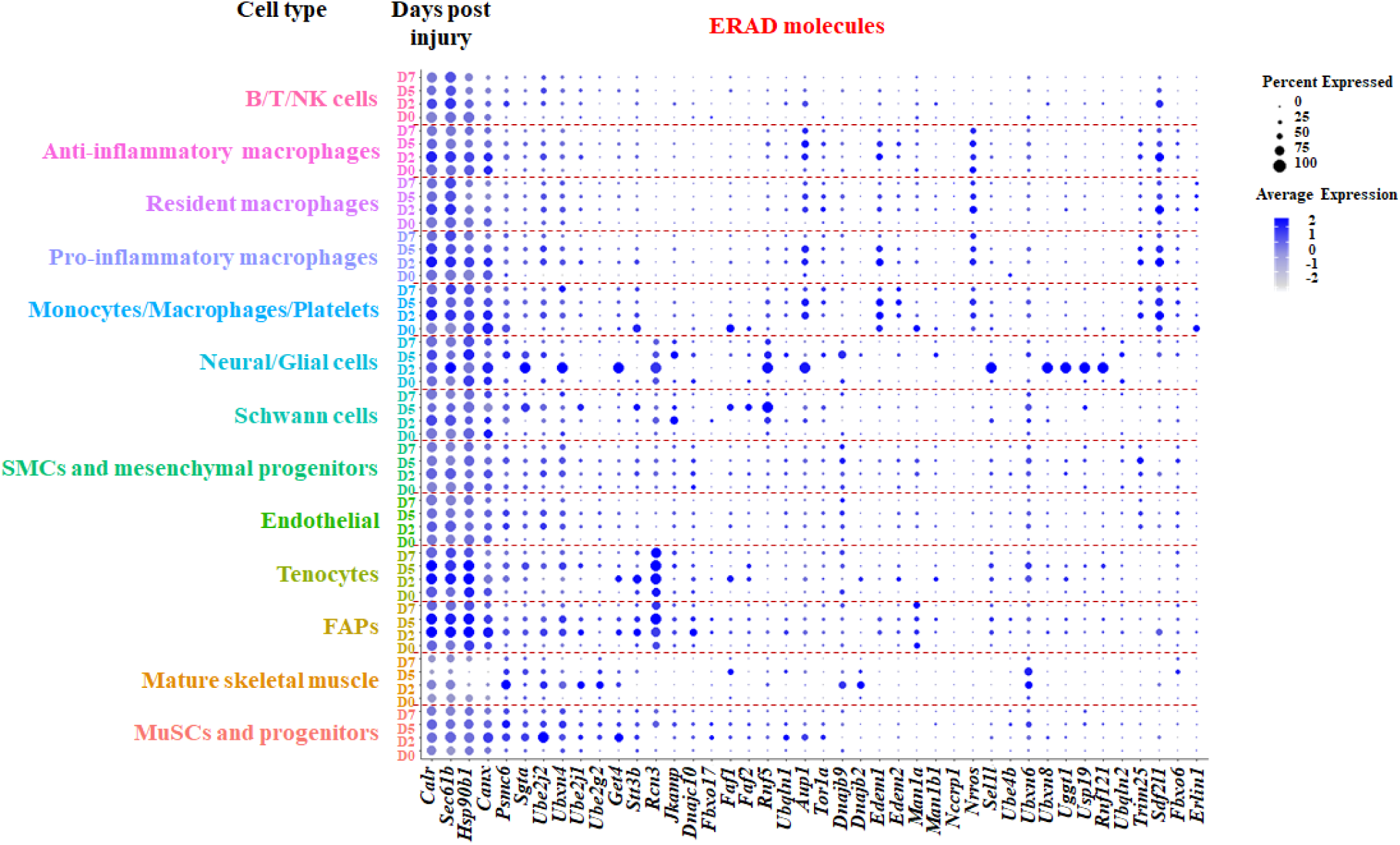
Gene expression of ERAD molecules during muscle regeneration. The scRNA-seq dataset was used to analyze the average expression levels and the percent expression of various genes. Dot plots show relative changes in gene expression of various ERAD molecules in different cell types and at indicated time points after muscle injury.

**Figure S3.**
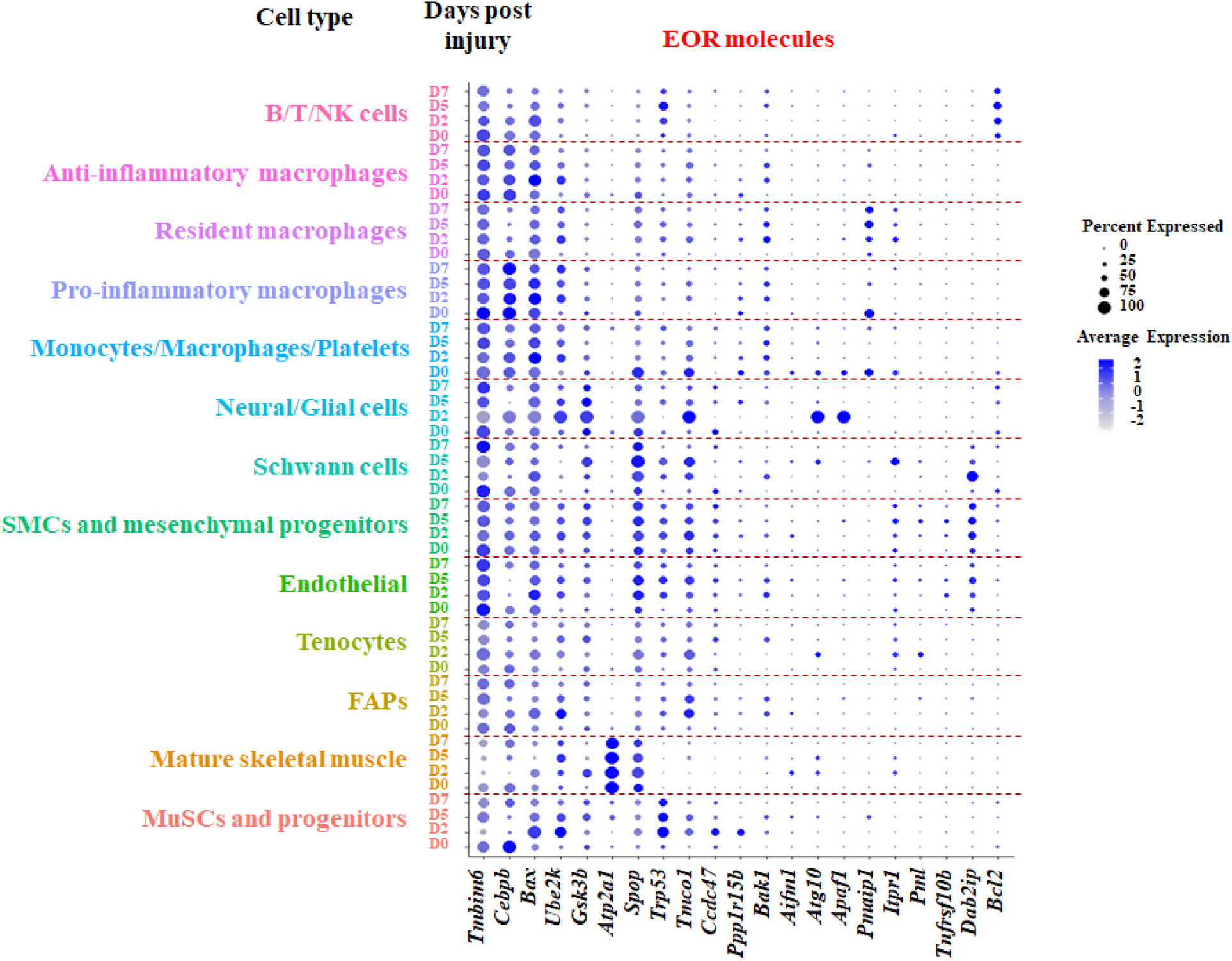
Gene expression of EOR molecules during muscle regeneration. The scRNA-seq dataset was used to analyze the average expression levels and the percentage expression of various genes. Dot plots show relative changes in gene expression of various EOR pathway molecules in various cell types and at indicated time points after muscle injury.

**Figure S4.**
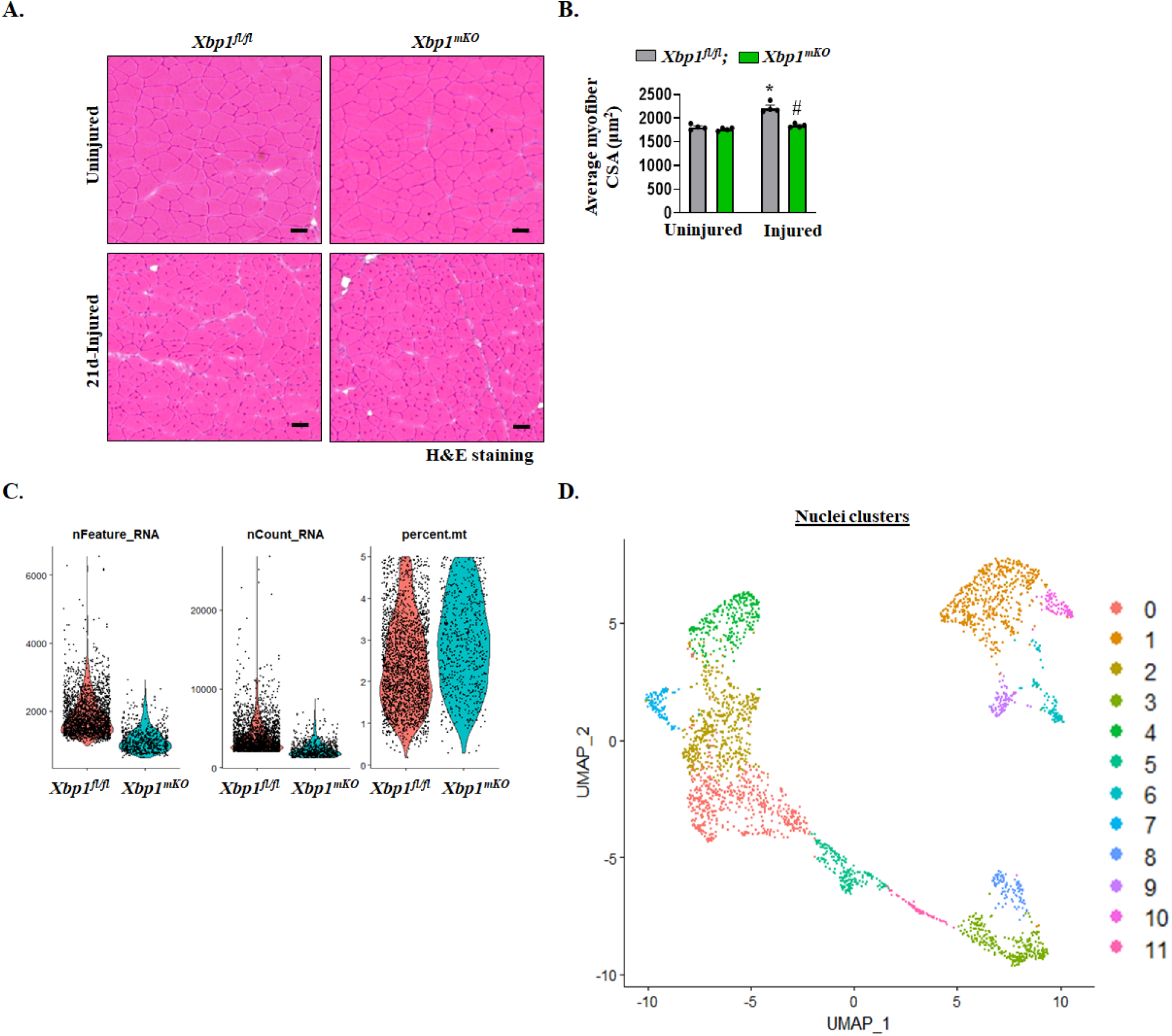
Morphometric assessment and preliminary analysis of single nucleus transcriptomics of regenerating muscle. TA muscle of *Xbp1^fl/fl^* and *Xbp1^mKO^*mice was injured using intramuscular injection of 1.2% BaCl_2_ and the muscle was collected at day 5 or 21 post injury. Uninjured muscle served as control. **(A)** Representative images of TA muscle sections after performing H&E staining. Scale bar, 50 µm. **(B)** quantification of average myofiber CSA of uninjured and 21d-injured TA muscle of *Xbp1^fl/fl^* and *Xbp1^mKO^* mice. n=4 mice in each group. Data are presented as mean ± SEM. *p ≤ 0.05, values significantly different from uninjured muscle of *Xbp1^fl/fl^* mice, and #p ≤ 0.05, values significantly different from 21d-injured muscle of *Xbp1^fl/fl^* mice analyzed by two-way ANOVA, followed by Tukey’s multiple comparison test. **(C)** 5d-injured TA muscle of *Xbp1^fl/fl^*and *Xbp1^mKO^* mice was used for single nucleus RNA-Sequencing (snRNA-seq). Nuclei were filtered based on 500 < nFeature_RNA < 20,000 and percent.mt < 5 for both the groups. **(D)** Using R software, nuclei were clustered for assessing spatial distribution. The UMAP plot shows 12 spatially distributed nuclei clusters in injured TA muscle of integrated dataset of *Xbp1^fl/fl^*and *Xbp1^mKO^* mice.

**Figure S5.**
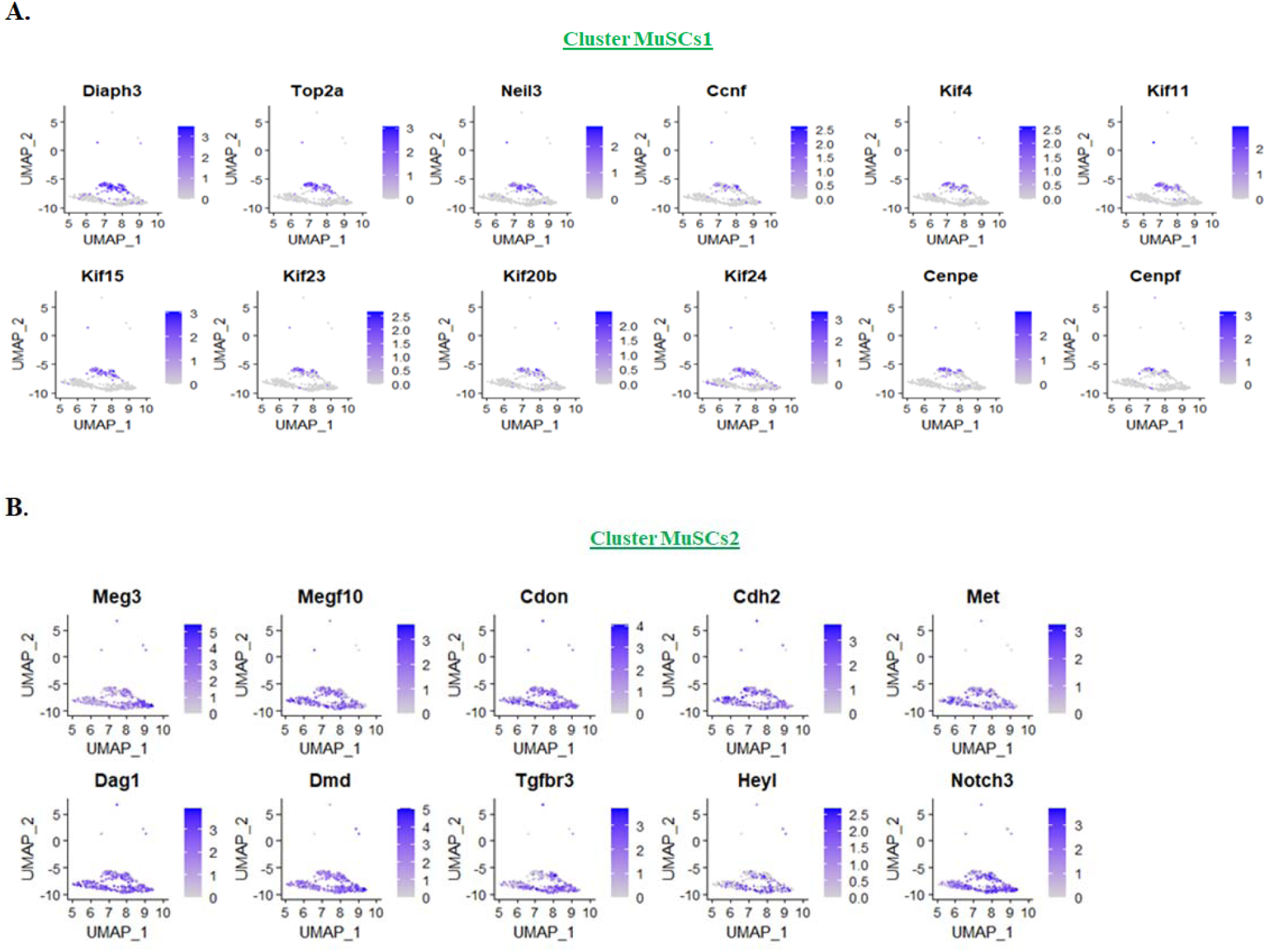
Characterization of functional differences in MuSCs1 and 2 clusters. Multi-clustering in snRNA-seq dataset were screened for the expression of various genes to identify distinct functional characteristics of **(A)** MuSCs1, and **(B)** MuSCs2 sub clusters.

**Figure S6.**
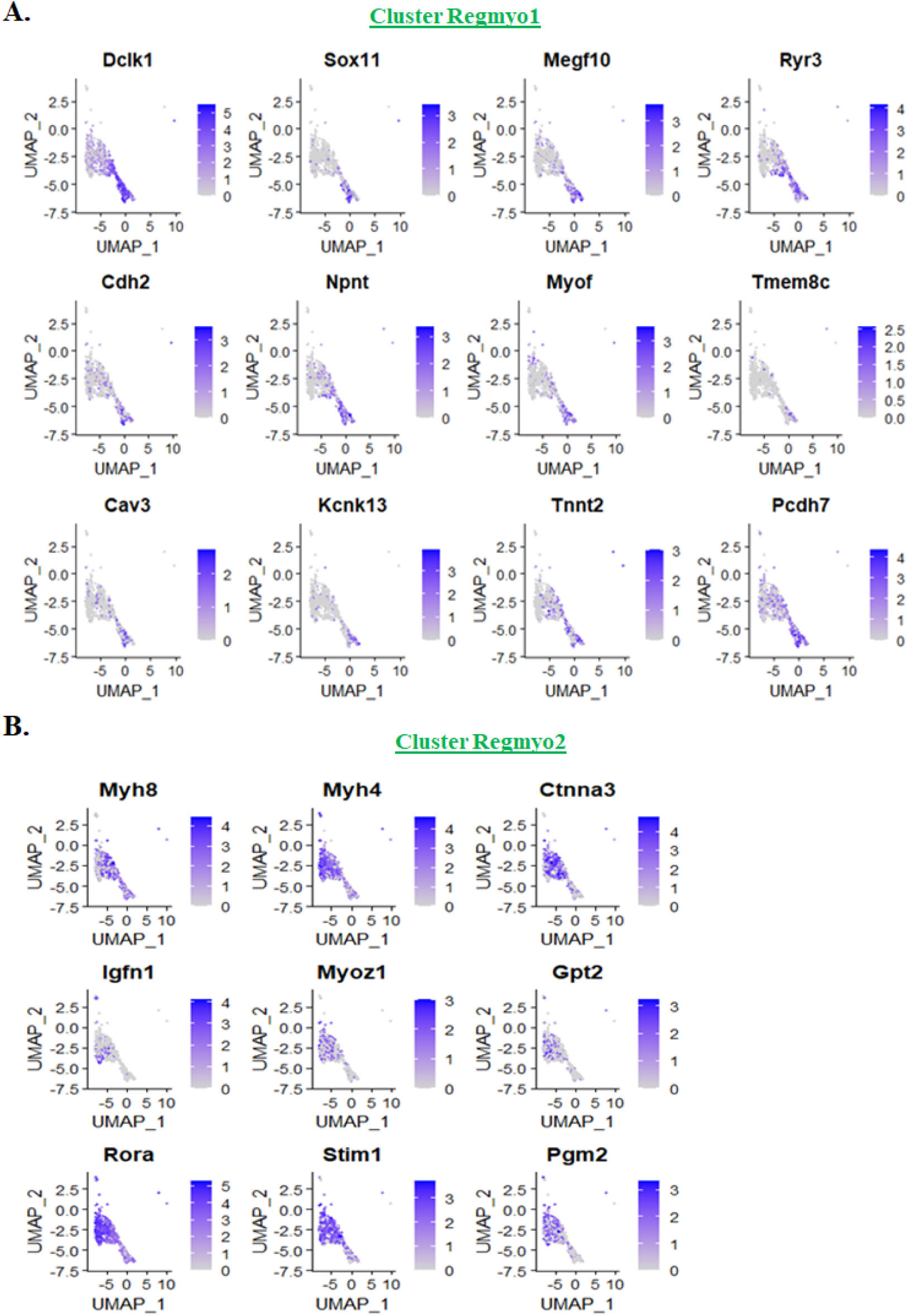
Characterization of functional differences in Regmyo1 and 2 clusters. Multi-clustering in snRNA-seq dataset were screened for the expression of various genes to identify distinct functional characteristics of **(A)** Regmyo1 and **(B)** Regmyo2 sub clusters.

**Figure S7.**
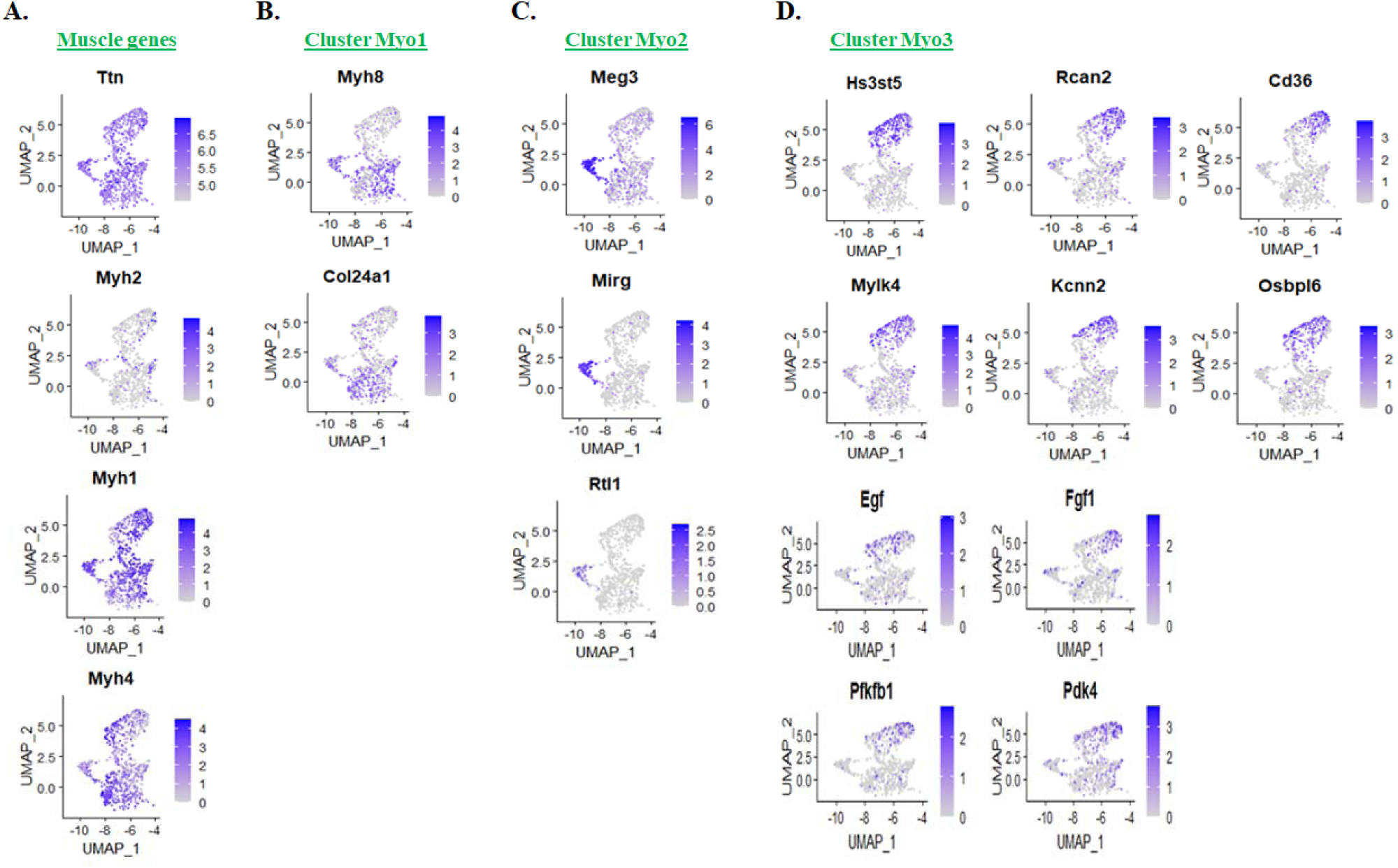
Characterization of functional differences in Myo1, 2 and 3 clusters. Multi-clustering in snRNA-seq dataset were screened for the expression of various genes to identify distinct functional characteristics of **(A)** muscle genes, **(B)** Myo1, **(C)** Myo2, and **(D)** Myo3 sub clusters.

**Figure S8.**
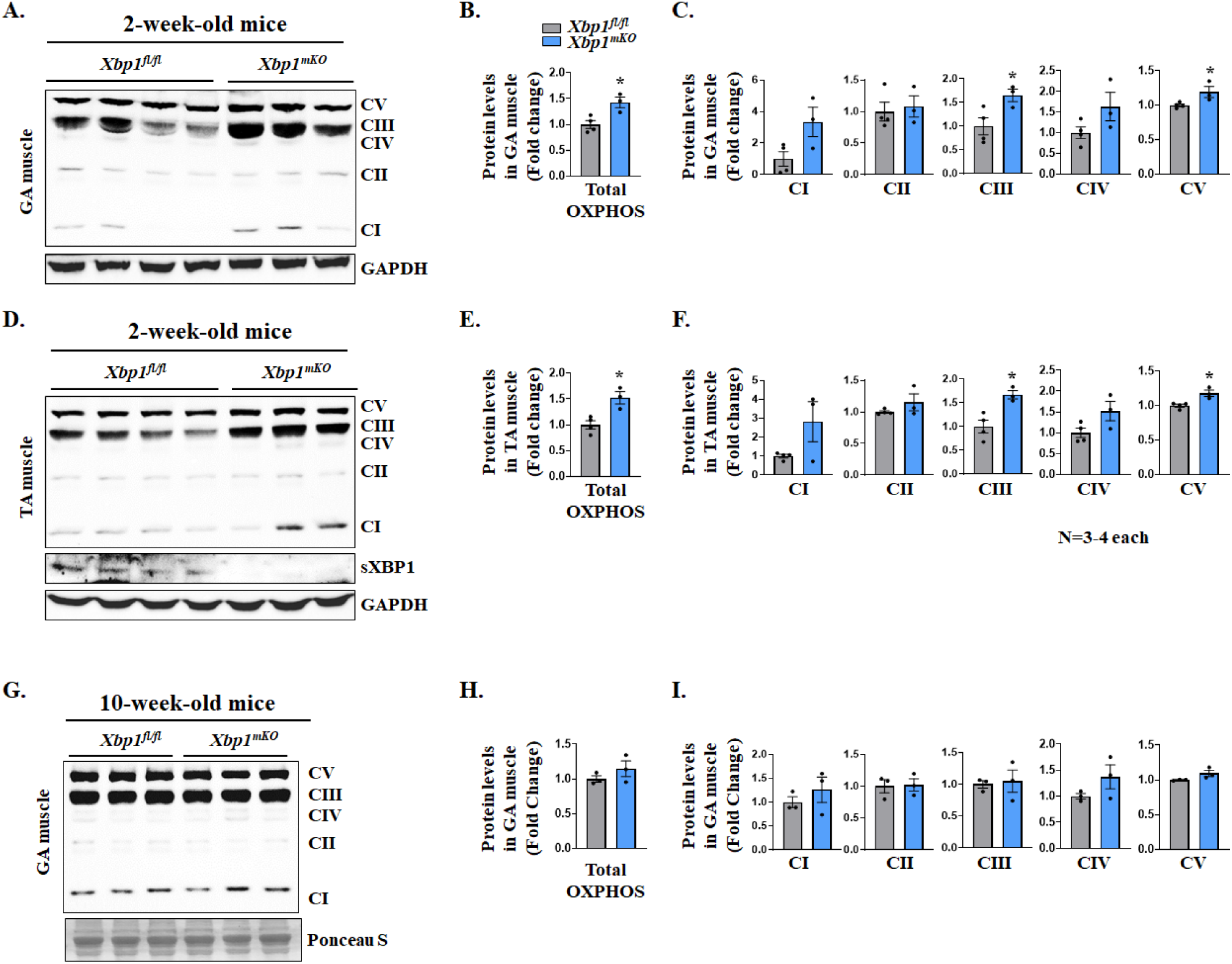
XBP1 regulates levels of mitochondrial OXPHOS protein during postnatal muscle development. TA and GA muscles of 2- and 10-week-old *Xbp1^fl/fl^* and *Xbp1^mKO^*mice were isolated and analyzed for the levels of mitochondrial OXPHOS proteins. **(A)** Immunoblots, and densitometry analysis of **(B)** total OXPHOS and **(C)** individual OXPHOS proteins of GA muscle of 2-week-old *Xbp1^fl/fl^* and *Xbp1^mKO^* mice. **(D)** Immunoblots and densitometry analysis of **(E)** total OXPHOS and **(F)** individual OXPHOS proteins of TA muscle of 2-week-old *Xbp1^fl/fl^* and *Xbp1^mKO^* mice. **(G)** Immunoblots and densitometry analysis of **(H)** total OXPHOS and **(I)** individual OXPHOS proteins in TA muscle of 10-week-old *Xbp1^fl/fl^*and *Xbp1^mKO^* mice. n=3-4 mice per group. All data are presented as mean ± SEM. *p ≤ 0.05; values significantly different from corresponding muscle of *Xbp1^fl/fl^* mice analyzed by unpaired Student *t* test.

**Figure S9.**
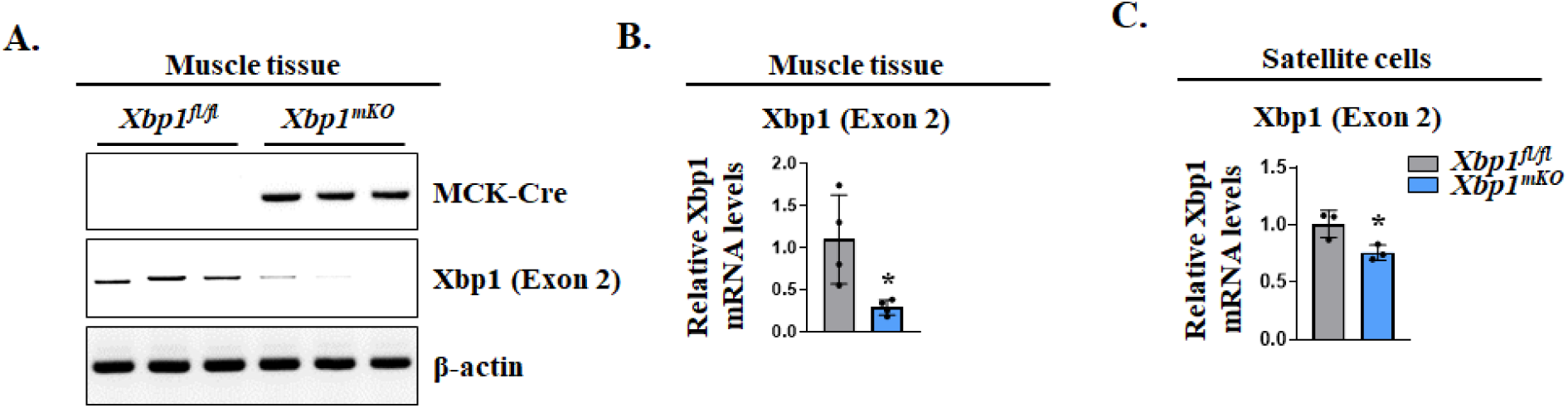
Transcript levels of *Xbp1* in skeletal muscle and satellite cells of Xbp1^mKO^ mice. **(A)** TA muscle of *Xbp1^fl/fl^* and *Xbp1^mKO^* mice was used for DNA extraction, followed by semi-quantitative PCR for Cre recombinase, *Xbp1* (exon2), and β-actin. Agarose gel images are presented here. n=3 mice in each group. **(B)** TA muscle of *Xbp1^fl/fl^* and *Xbp1^mKO^* mice was used for RNA extraction, followed by qRT-PCR analysis for *Xbp1* (exon2) transcripts levels. Relative mRNA levels of *Xbp1* (exon2) in TA muscle of *Xbp1^fl/fl^* and *Xbp1^mKO^*mice are presented. n=4 mice in each group. **(C)** Freshly isolated satellite cells from hindlimb muscles of *Xbp1^fl/fl^* and *Xbp1^mKO^* mice were analyzed for *Xbp1* (exon2) mRNA levels by performing qRT-PCR assay. Relative mRNA levels of *Xbp1* (exon2) in the two group are presented. n=3 biological replicates per group. All data are presented as mean ± SEM. *p ≤ 0.05; values significantly different from TA muscle or satellite cells of *Xbp1^fl/fl^*mice analyzed by unpaired Student *t* test.

**Figure S10.**
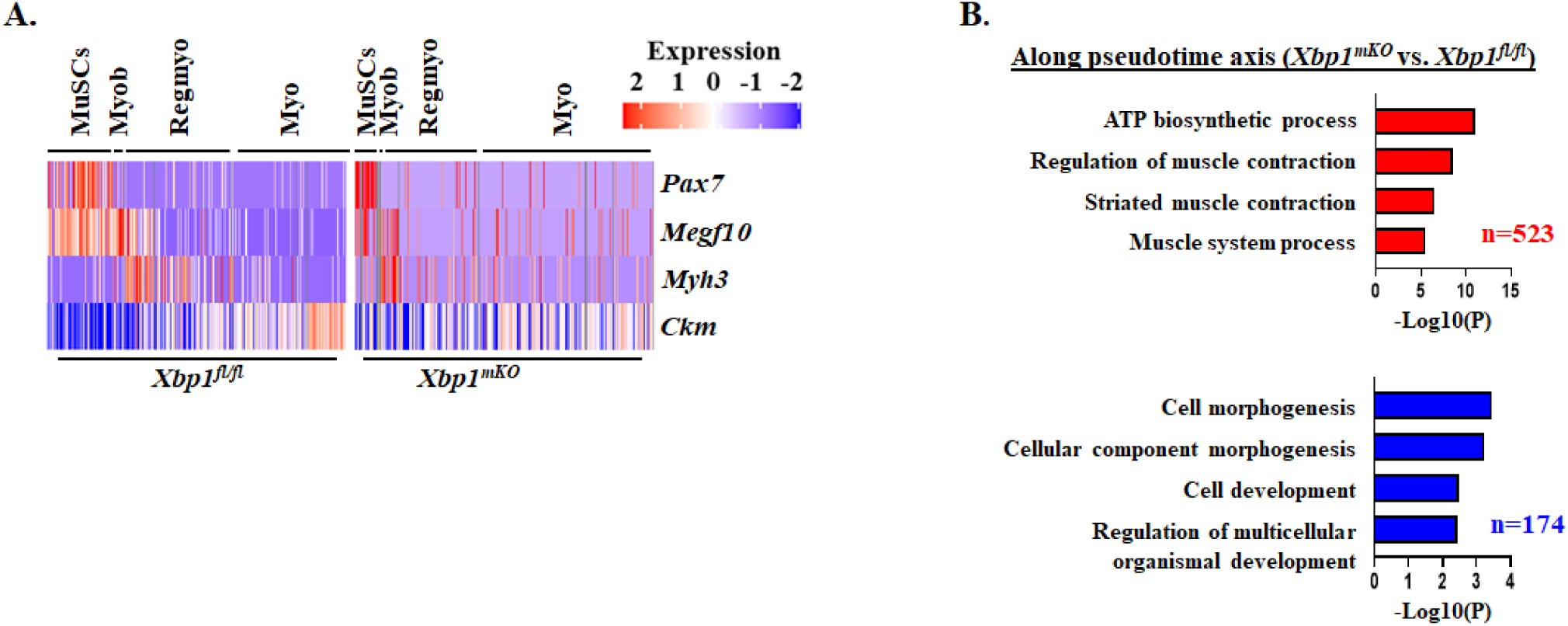
XBP1 regulates myogenesis trajectory and alters population and function of progenitor cells during muscle regeneration. **(A)** Heatmaps of *Pax7*, *Megf10*, *Myh3* and *Ckm* gene expression in the snRNA-seq dataset of regenerating muscle of *Xbp1^fl/fl^* and *Xbp1^mKO^* mice. **(B)** Pseudotime-based trajectory analysis was performed using Monocle2 package. Gene Ontology (GO) term analysis for identification of biological processes associated with upregulated (red) and downregulated (blue) genes along the pseudotime axis.

**Figure S11.**
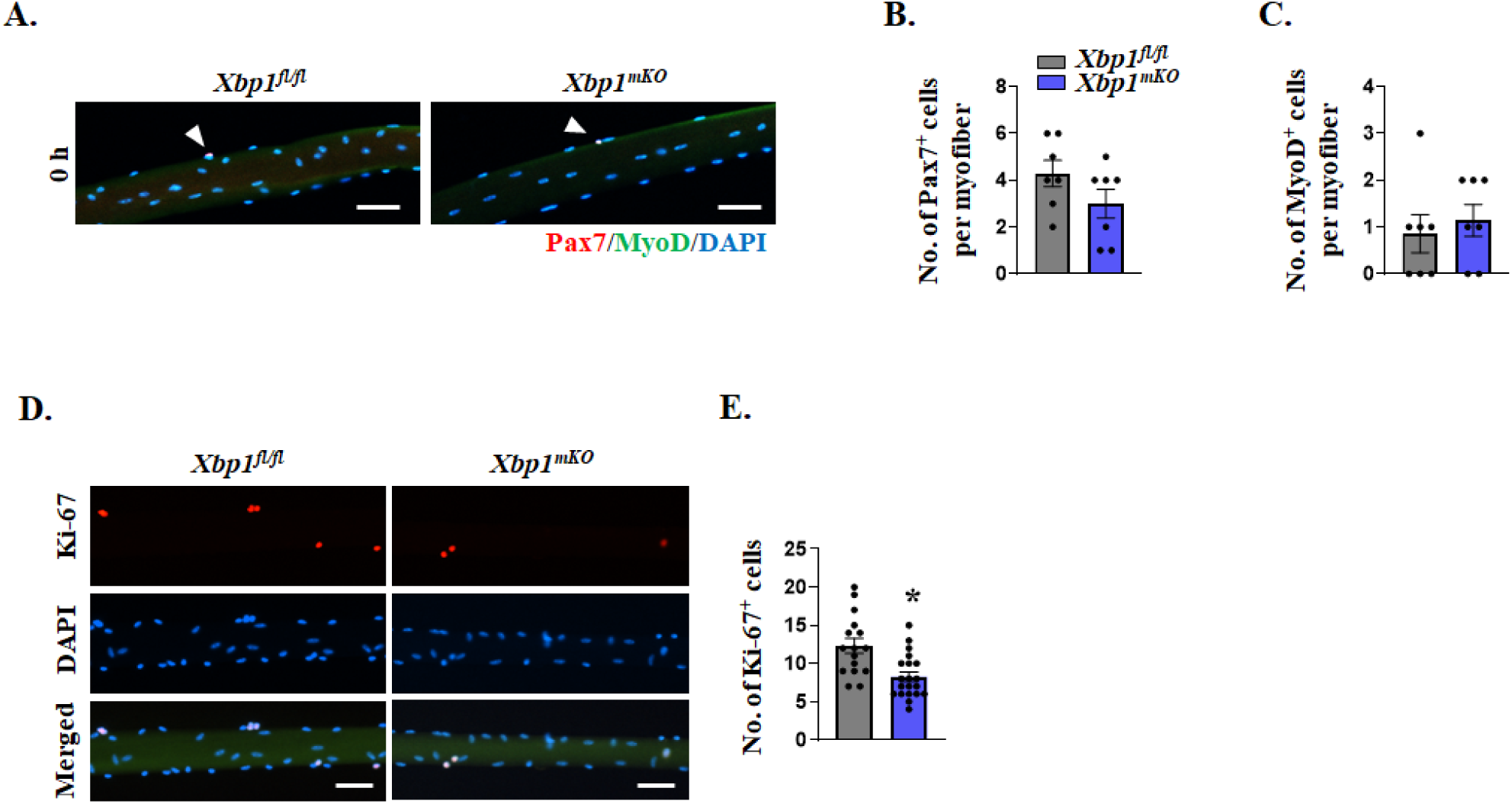
XBP1 regulates proliferation of satellite cells in an *ex vivo* model of muscle injury. Single myofiber cultures were established from EDL muscle of *Xbp1^fl/fl^* and *Xbp1^mKO^* mice and fixed immediately or after 48 h in culture, followed by immunostaining analysis. **(A)** Representative images of single myofibers isolated from EDL muscle of *Xbp1^fl/fl^* and *Xbp1^mKO^*mice after immediate fixation and immunostaining for Pax7 and MyoD protein. DAPI was used to identify nuclei. Scale bar, 50 µm. Quantification of number of **(D)** Pax7^+^ and **(E)** MyoD^+^ cells per myofiber. Scale bar, 50 µm. n=7 biological replicates per group. After 48 h of culturing, the myofiber-associated cells were stained with anti-Ki67 and DAPI to identify proliferating cells. **(D)** Representative images and **(E)** quantification of number of Ki67^+^ cells per myofiber in *Xbp1^fl/fl^* and *Xbp1^mKO^* cultures. n=15-20 myofibers per group. Data are presented as mean ± SEM. *p ≤ 0.05; values significantly different from myofibers of *Xbp1^fl/fl^* mice analyzed by unpaired Student *t* test.

**Figure S12.**
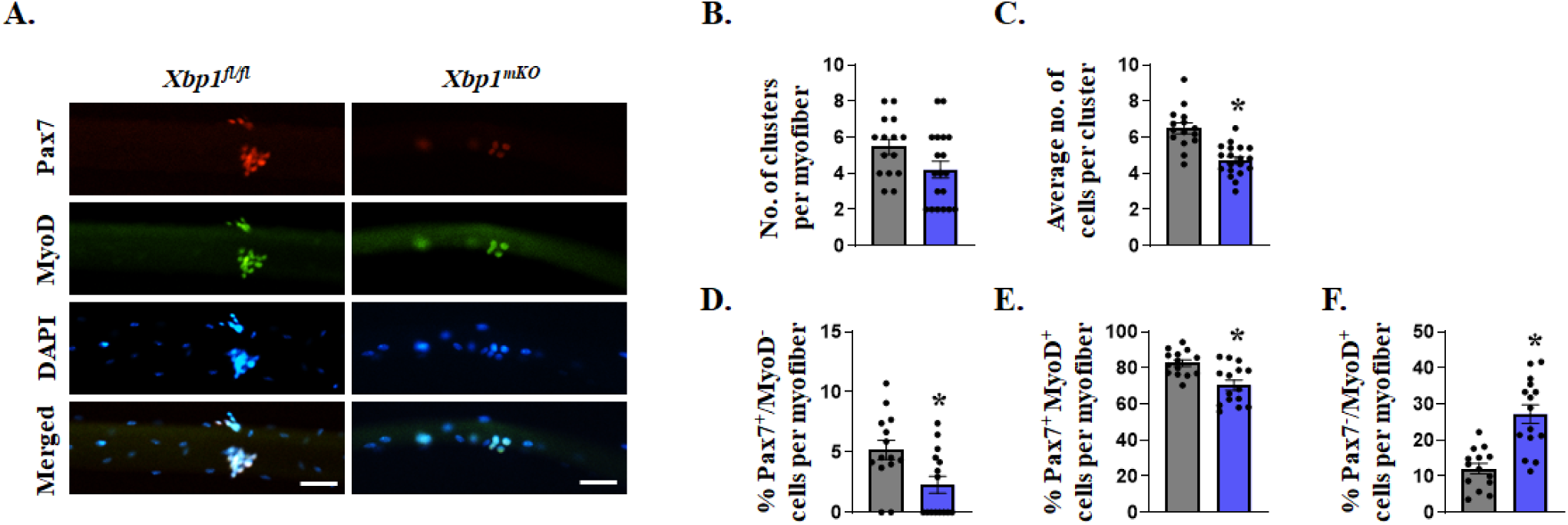
XBP1 regulates self-renewal and differentiation potential of satellite cells. Single myofibers were isolated from EDL muscle of *Xbp1^fl/fl^* and *Xbp1^mKO^* mice and cultured for 72 h, followed by immunostaining with anti-Pax7, anti-MyoD and DAPI to identify self-renewing (Pax7^+^/MyoD^-^), proliferating (Pax7^+^/MyoD^+^) and differentiating (Pax7^-^/MyoD^+^) satellite cells. **(A)** Representative images immunostained myofibers. Scale bar, 50 µm. Quantification of **(B)** number of clusters per myofiber, **(C)** average number of cells per cluster, and proportion of **(D)** self-renewing, **(E)** proliferating, and **(F)** differentiating satellite cells per myofiber of *Xbp1^fl/fl^* and *Xbp1^mKO^* mice. n=15-20 myofibers per group. All data are presented as mean ± SEM. *p ≤ 0.05; values significantly different from myofibers of *Xbp1^fl/fl^* mice analyzed by unpaired Student *t* test.

**Figure S13.**
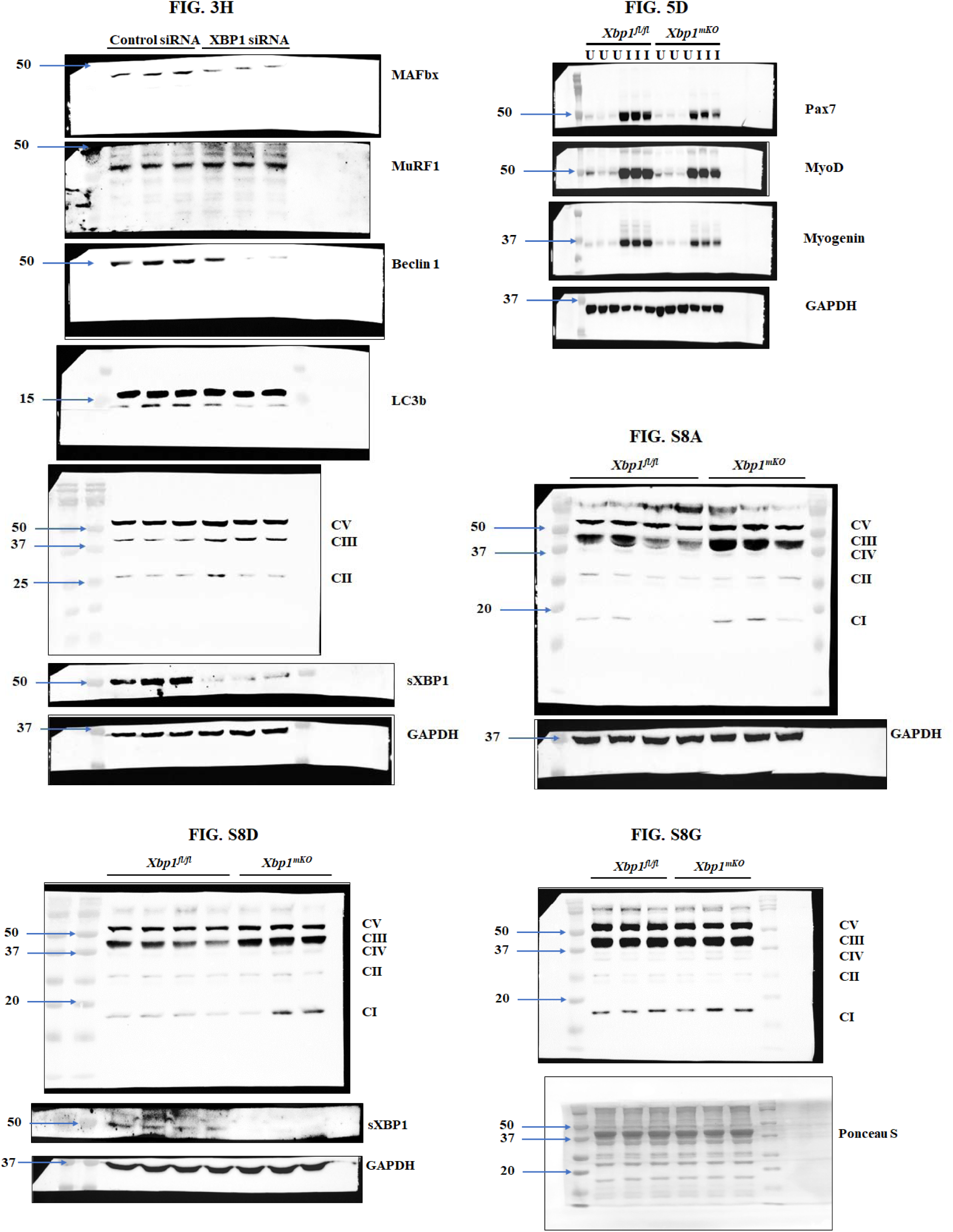
Uncropped Western blot images. Original immunoblots generated in the present study.

