## Supplementary material for "Single-nucleus transcriptomic analysis reveals the regulatory circuitry of myofiber XBP1 during regenerative myogenesis": Figures S1-S13

### Supplemental Figures S1-S13 and Legends

#### FIGURE S1

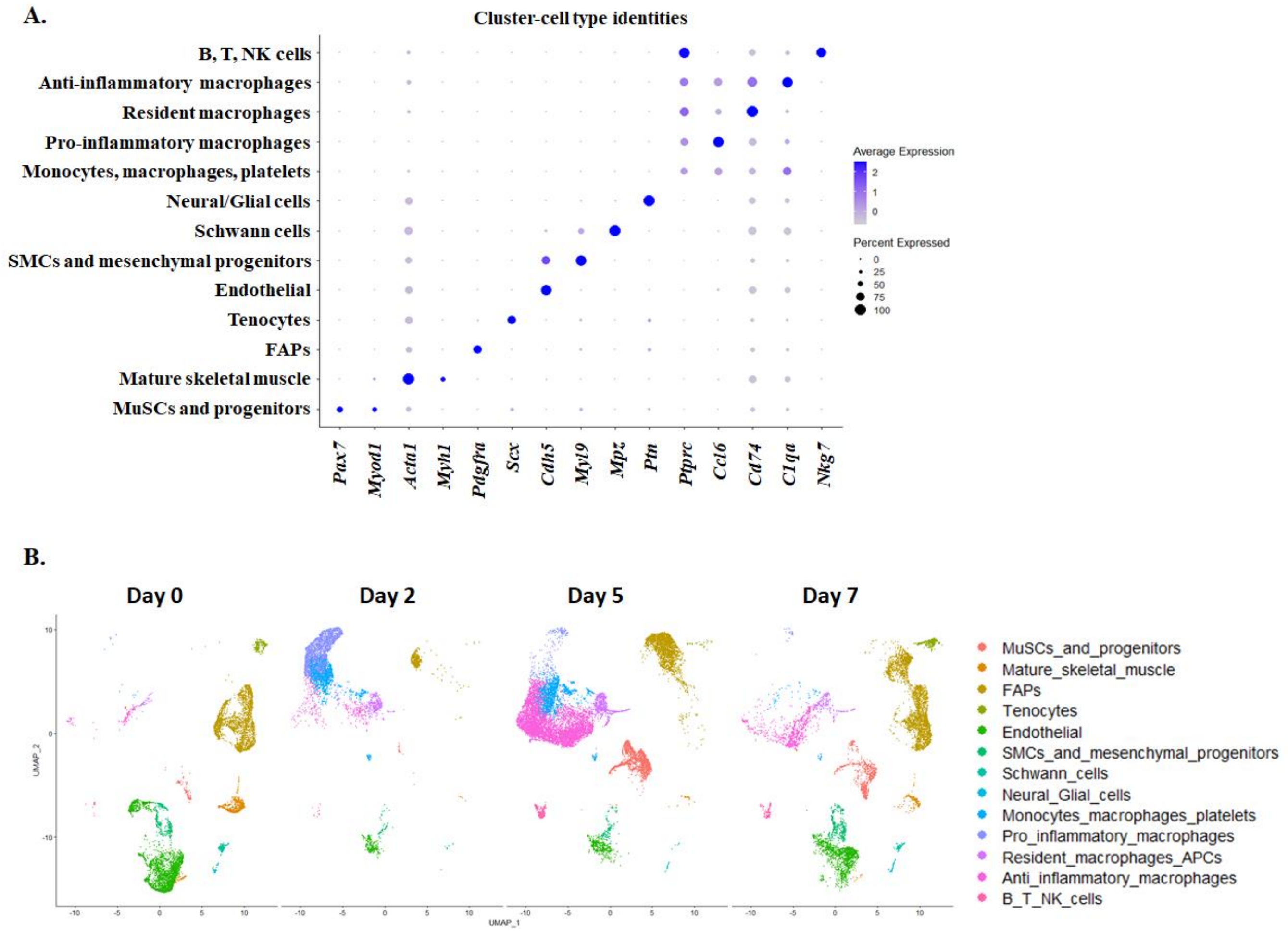


**Figure S1. Analysis of scRNA-seq dataset identifies distinct cell types during muscle regeneration.** The scRNA-seq dataset (GSE143437), consisting of regenerating muscle tissue samples at day 0, 2, 5 and 7 post injury, was analyzed using R software (v4.2.2). Gene expression of various markers of cellular identification were analyzed to annotate distinct cell types of the spatially distributed clusters. **(A)** Dot plot shows average expression and percentage of cells expressing the indicated genes in different cell types. **(B)** Split-UMAPs visually represent the changes in proportion of various cell types at different time points of muscle regeneration.

**FIGURE S2**

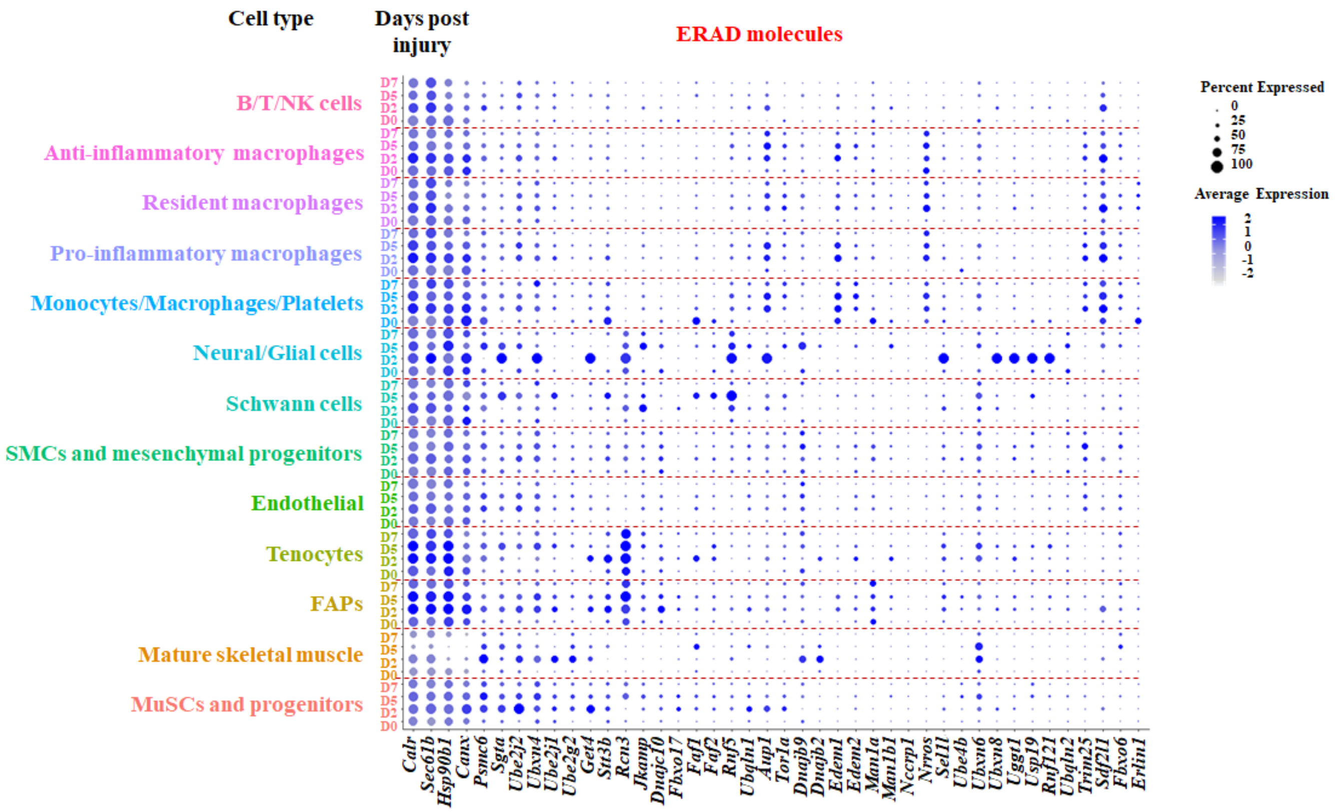


**Figure S2. Gene expression of ERAD molecules during muscle regeneration.** The scRNA-seq dataset was used to analyze the average expression levels and the percent expression of various genes. Dot plots show relative changes in gene expression of various ERAD molecules in different cell types and at indicated time points after muscle injury.

FIGURE S3

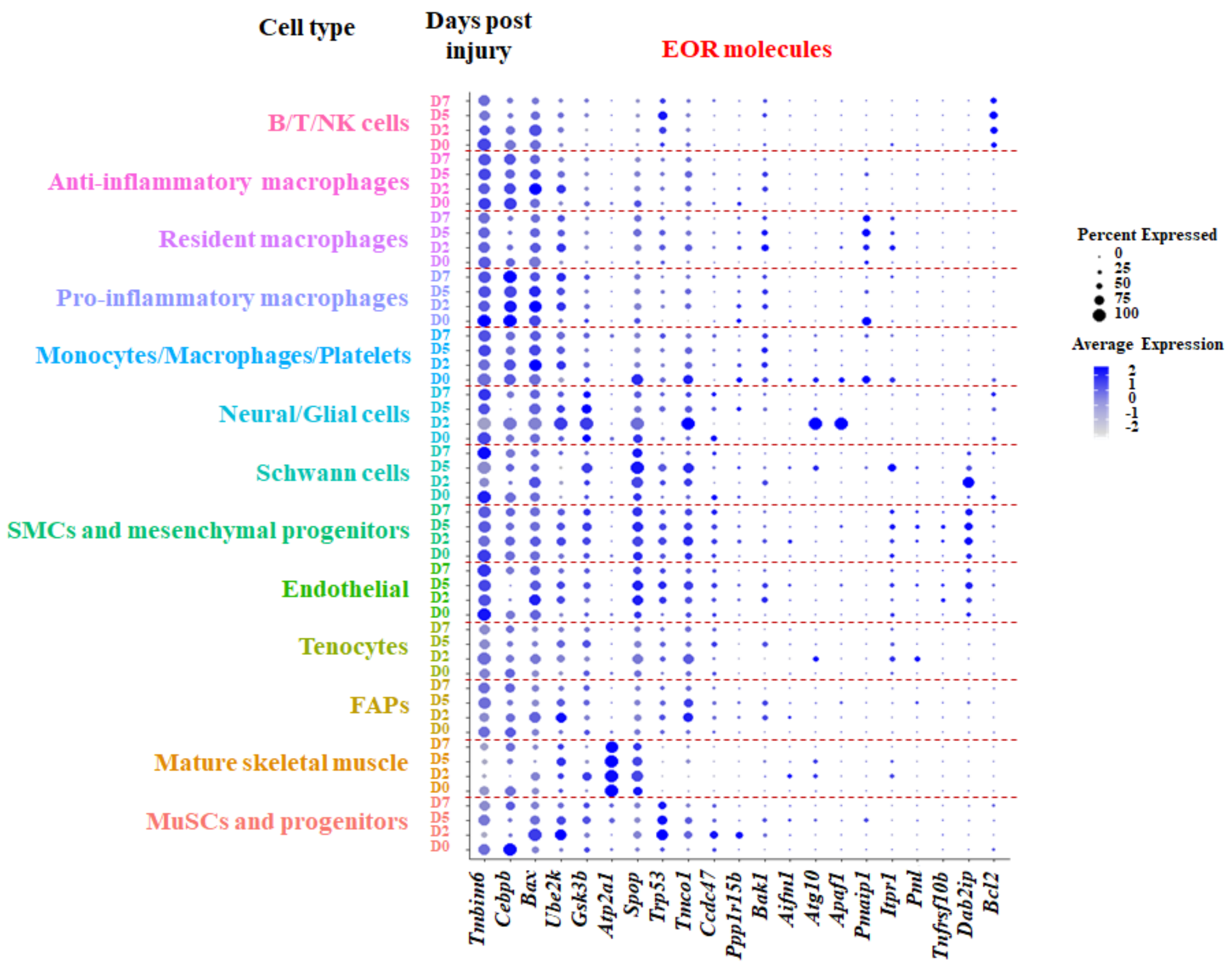


**Figure S3. Gene expression of EOR molecules during muscle regeneration.** The scRNA-seq dataset was used to analyze the average expression levels and the percentage expression of various genes. Dot plots show relative changes in gene expression of various EOR pathway molecules in various cell types and at indicated time points after muscle injury.

**FIGURE S4**

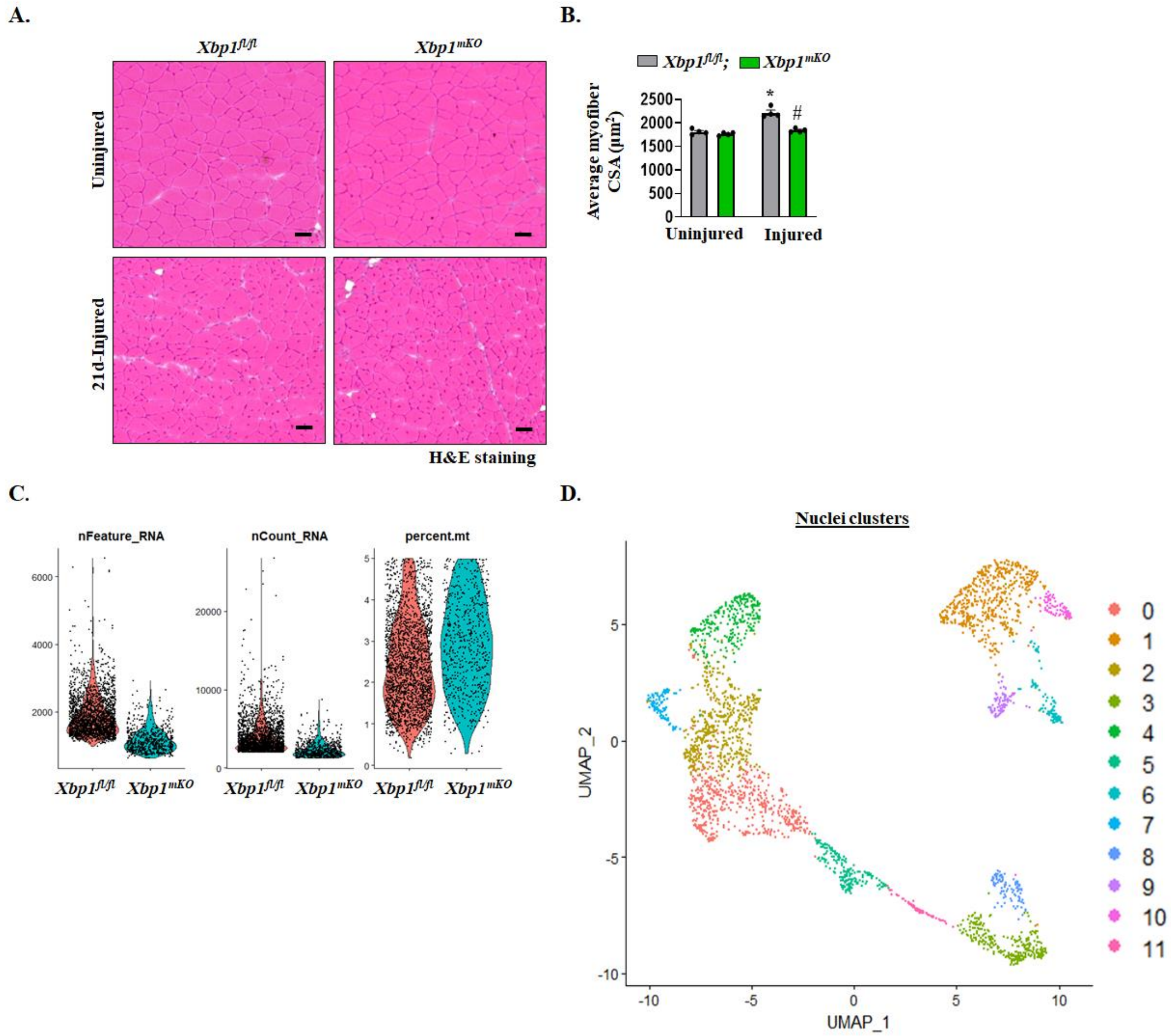


**Figure S4. Morphometric assessment and preliminary analysis of single nucleus transcriptomics of regenerating muscle.** TA muscle of *Xbp1*$^{fl/fl}$ and *Xbp1*$^{mKO}$ mice was injured using intramuscular injection of 1.2% $BaCl_2$ and the muscle was collected at day 5 or 21 post injury. Uninjured muscle served as control. **(A)** Representative images of TA muscle sections after performing H&E staining. Scale bar, 50 µm. **(B)** quantification of average myofiber CSA of uninjured and 21d-injured TA muscle of *Xbp1*$^{fl/fl}$ and *Xbp1*$^{mKO}$ mice. n=4 mice in each group. Data are presented as mean ± SEM. *$p \leq 0.05$, values significantly different from uninjured muscle of *Xbp1*$^{fl/fl}$ mice, and #$p \leq 0.05$, values significantly different from 21d-injured muscle of *Xbp1*$^{fl/fl}$ mice analyzed by two-way ANOVA, followed by Tukey's multiple comparison test. **(C)** 5d-injured TA muscle of *Xbp1*$^{fl/fl}$ and *Xbp1*$^{mKO}$ mice was used for single nucleus RNA-Sequencing (snRNA-seq). Nuclei were filtered based on 500 < nFeature_RNA < 20,000 and percent.mt < 5 for both the groups. **(D)** Using R software, nuclei were clustered for assessing spatial distribution. The UMAP plot shows 12 spatially distributed nuclei clusters in injured TA muscle of integrated dataset of *Xbp1*$^{fl/fl}$ and *Xbp1*$^{mKO}$ mice.

**FIGURE S5**

**A.**

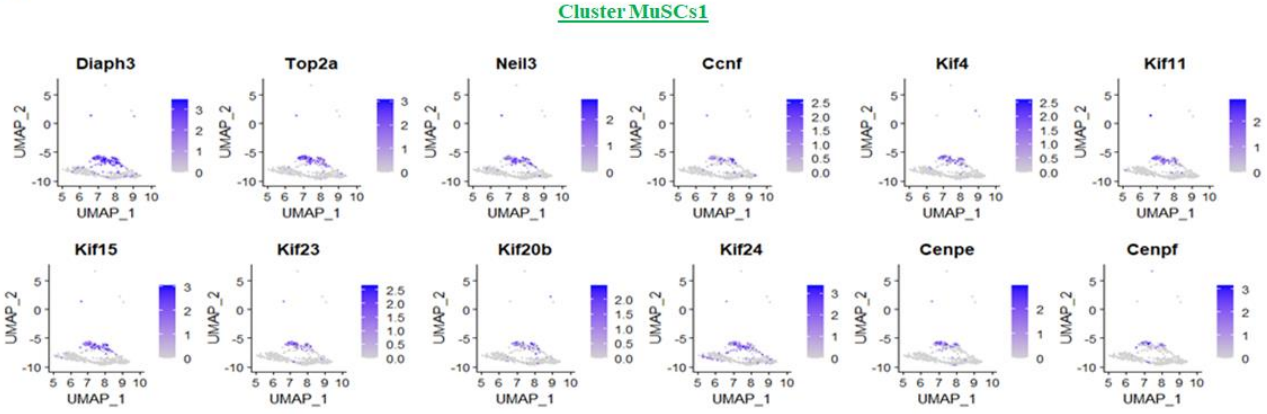


**B.**

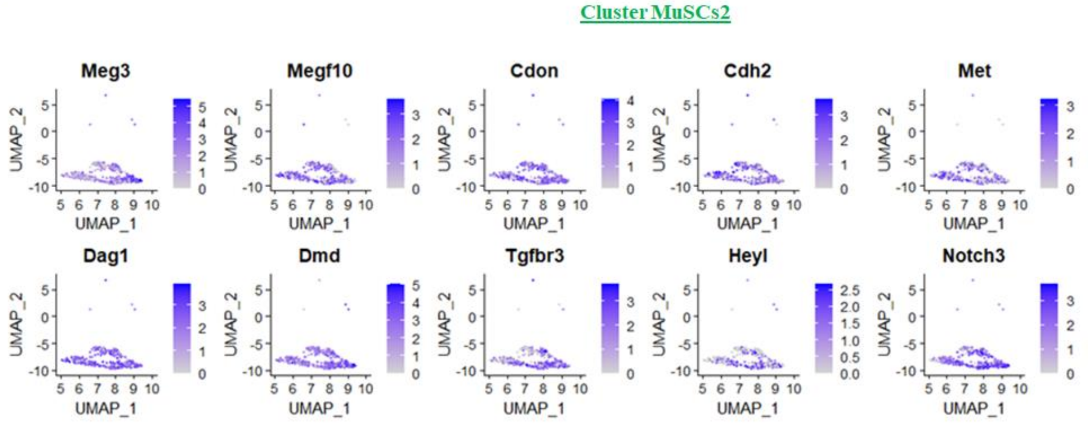


**Figure S5. Characterization of functional differences in MuSCs1 and 2 clusters.** Multi-clustering in snRNA-seq dataset were screened for the expression of various genes to identify distinct functional characteristics of **(A)** MuSCs1, and **(B)** MuSCs2 sub clusters.

**FIGURE S6**

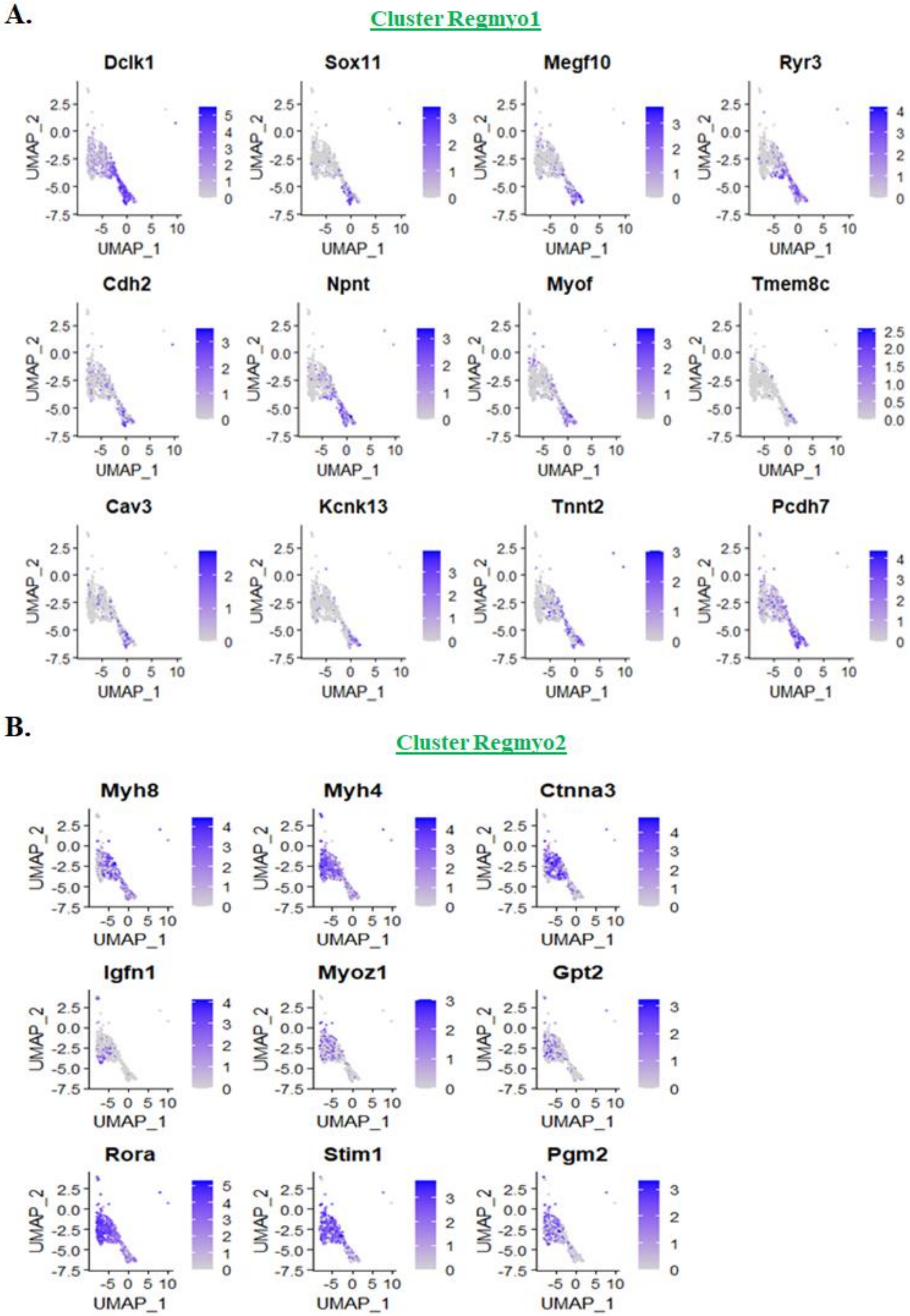


**Figure S6. Characterization of functional differences in Regmyo1 and 2 clusters.** Multi-clustering in snRNA-seq dataset were screened for the expression of various genes to identify distinct functional characteristics of **(A)** Regmyo1 and **(B)** Regmyo2 sub clusters.

FIGURE S7

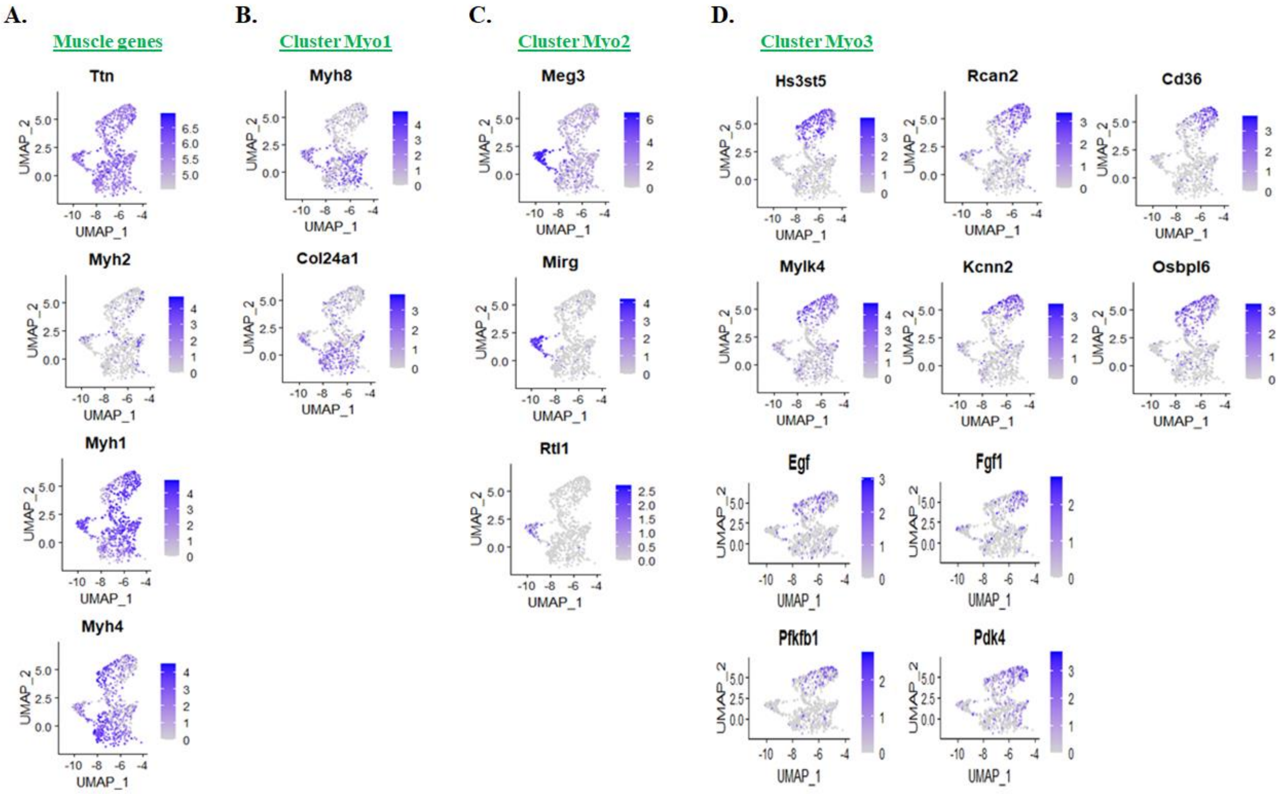


**Figure S7. Characterization of functional differences in Myo1, 2 and 3 clusters.** Multi-clustering in snRNA-seq dataset were screened for the expression of various genes to identify distinct functional characteristics of **(A)** muscle genes, **(B)** Myo1, **(C)** Myo2, and **(D)** Myo3 sub clusters.

FIGURE S8

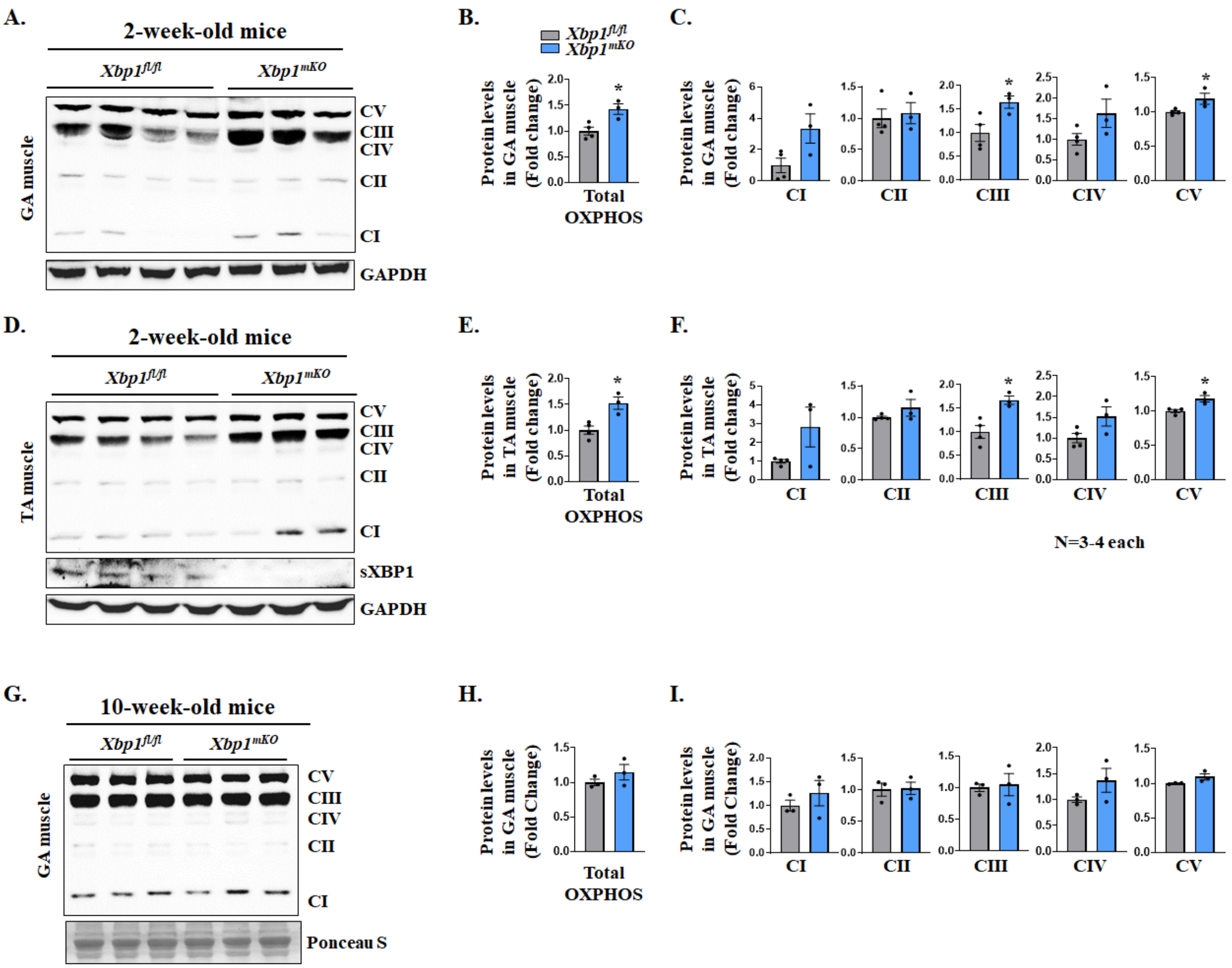


**Figure S8. XBP1 regulates levels of mitochondrial OXPHOS protein during postnatal muscle development.** TA and GA muscles of 2- and 10-week-old *Xbp1*$^{fl/fl}$ and *Xbp1*$^{mKO}$ mice were isolated and analyzed for the levels of mitochondrial OXPHOS proteins. **(A)** Immunoblots, and densitometry analysis of **(B)** total OXPHOS and **(C)** individual OXPHOS proteins of GA muscle of 2-week-old *Xbp1*$^{fl/fl}$ and *Xbp1*$^{mKO}$ mice. **(D)** Immunoblots and densitometry analysis of **(E)** total OXPHOS and **(F)** individual OXPHOS proteins of TA muscle of 2-week-old *Xbp1*$^{fl/fl}$ and *Xbp1*$^{mKO}$ mice. **(G)** Immunoblots and densitometry analysis of **(H)** total OXPHOS and **(I)** individual OXPHOS proteins in TA muscle of 10-week-old *Xbp1*$^{fl/fl}$ and *Xbp1*$^{mKO}$ mice. n=3-4 mice per group. All data are presented as mean ± SEM. *$p \leq 0.05$; values significantly different from corresponding muscle of *Xbp1*$^{fl/fl}$ mice analyzed by unpaired Student *t* test.

FIGURE S9

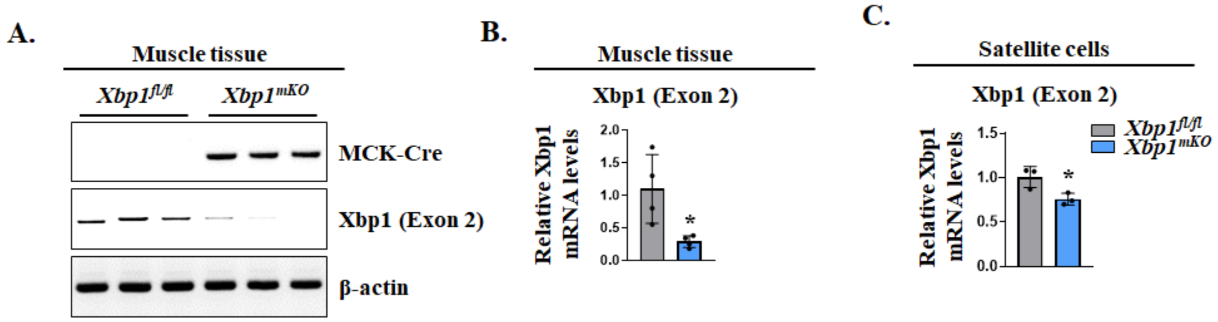


**Figure S9. Transcript levels of *Xbp1* in skeletal muscle and satellite cells of Xbp1$^{mKO}$ mice.** **(A)** TA muscle of *Xbp1*$^{fl/fl}$ and *Xbp1*$^{mKO}$ mice was used for DNA extraction, followed by semi-quantitative PCR for Cre recombinase, *Xbp1* (exon2), and β-actin. Agarose gel images are presented here. n=3 mice in each group. **(B)** TA muscle of *Xbp1*$^{fl/fl}$ and *Xbp1*$^{mKO}$ mice was used for RNA extraction, followed by qRT- PCR analysis for *Xbp1* (exon2) transcripts levels. Relative mRNA levels of *Xbp1* (exon2) in TA muscle of *Xbp1*$^{fl/fl}$ and *Xbp1*$^{mKO}$ mice are presented. n=4 mice in each group. **(C)** Freshly isolated satellite cells from hindlimb muscles of *Xbp1*$^{fl/fl}$ and *Xbp1*$^{mKO}$ mice were analyzed for *Xbp1* (exon2) mRNA levels by performing qRT-PCR assay. Relative mRNA levels of *Xbp1* (exon2) in the two group are presented. n=3 biological replicates per group. All data are presented as mean ± SEM. *$p \leq 0.05$; values significantly different from TA muscle or satellite cells of *Xbp1*$^{fl/fl}$ mice analyzed by unpaired Student *t* test.

**FIGURE S10**

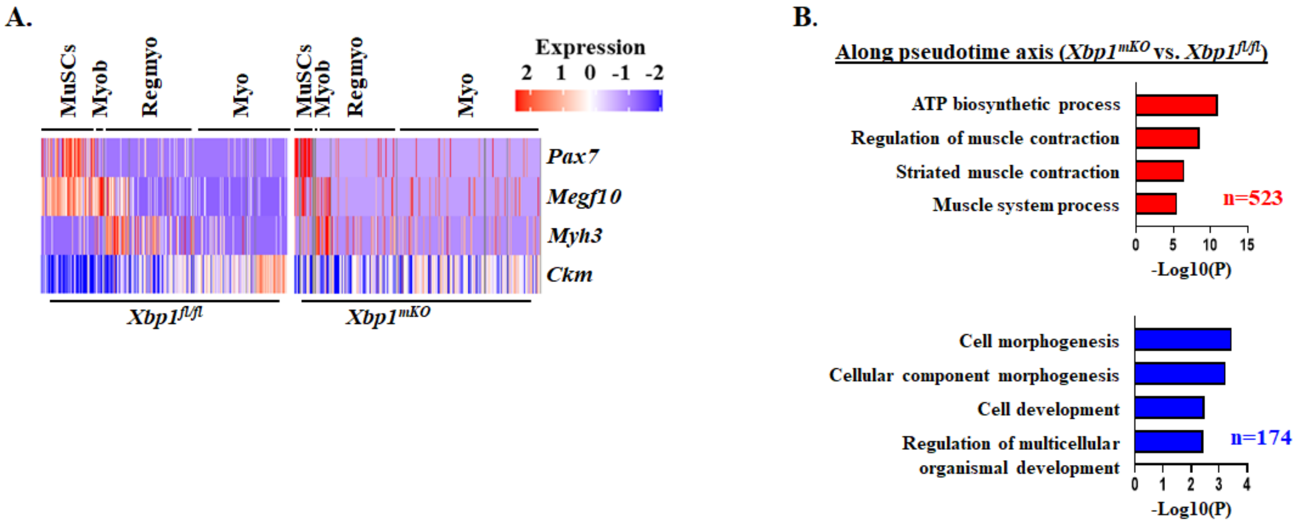


**Figure S10. XBP1 regulates myogenesis trajectory and alters population and function of progenitor cells during muscle regeneration. (A)** Heatmaps of *Pax7*, *Megf10*, *Myh3* and *Ckm* gene expression in the snRNA-seq dataset of regenerating muscle of *Xbp1*$^{fl/fl}$ and *Xbp1*$^{mKO}$ mice. **(B)** Pseudotime-based trajectory analysis was performed using Monocle2 package. Gene Ontology (GO) term analysis for identification of biological processes associated with upregulated (red) and downregulated (blue) genes along the pseudotime axis.

**FIGURE S11**

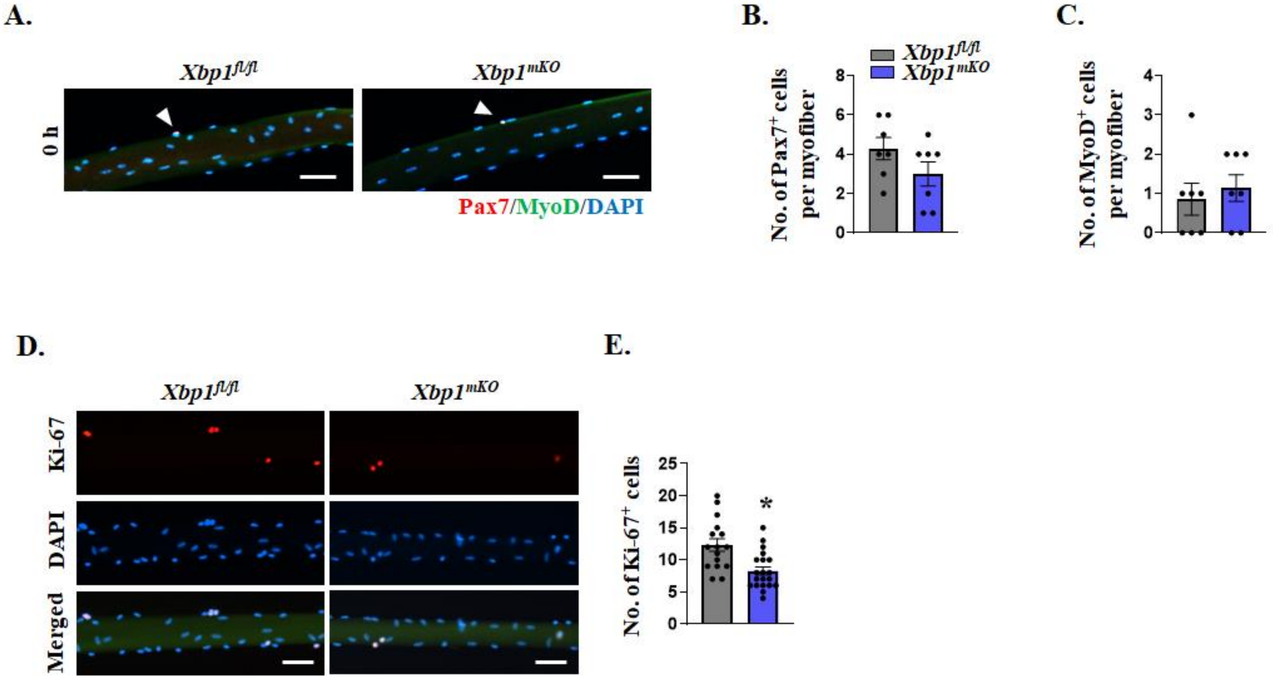


**Figure S11. XBP1 regulates proliferation of satellite cells in an *ex vivo* model of muscle injury.** Single myofiber cultures were established from EDL muscle of *Xbp1*$^{fl/fl}$ and *Xbp1*$^{mKO}$ mice and fixed immediately or after 48 h in culture, followed by immunostaining analysis. **(A)** Representative images of single myofibers isolated from EDL muscle of *Xbp1*$^{fl/fl}$ and *Xbp1*$^{mKO}$ mice after immediate fixation and immunostaining for Pax7 and MyoD protein. DAPI was used to identify nuclei. Scale bar, 50 µm. Quantification of number of **(D)** Pax7$^{+}$ and **(E)** MyoD$^{+}$ cells per myofiber. Scale bar, 50 µm. n=7 biological replicates per group. After 48 h of culturing, the myofiber-associated cells were stained with anti-Ki67 and DAPI to identify proliferating cells. **(D)** Representative images and **(E)** quantification of number of Ki67$^{+}$ cells per myofiber in *Xbp1*$^{fl/fl}$ and *Xbp1*$^{mKO}$ cultures. n=15-20 myofibers per group. Data are presented as mean ± SEM. *$p \leq 0.05$; values significantly different from myofibers of *Xbp1*$^{fl/fl}$ mice analyzed by unpaired Student *t* test.

**FIGURE S12**

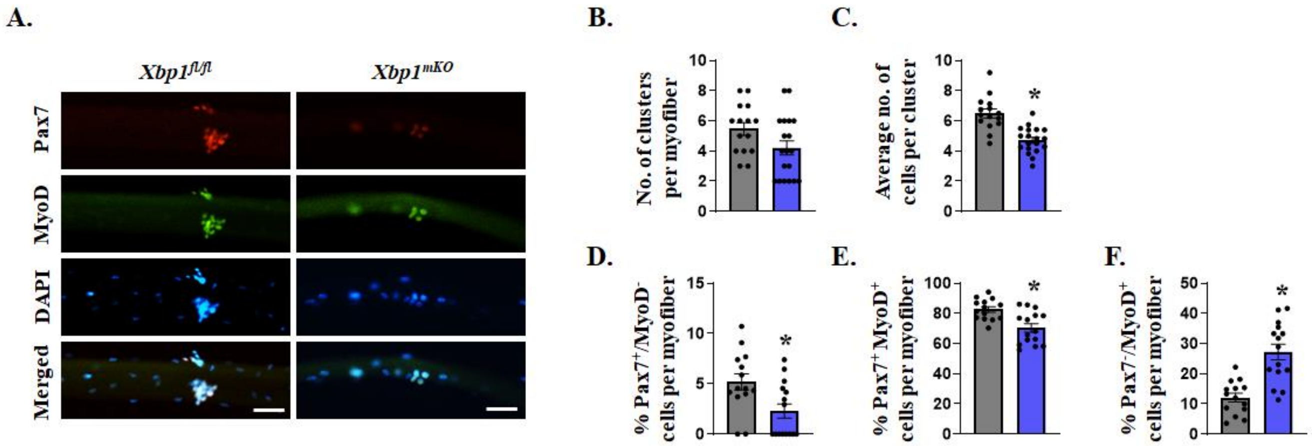


**Figure S12. XBP1 regulates self-renewal and differentiation potential of satellite cells.** Single myofibers were isolated from EDL muscle of *Xbp1*$^{fl/fl}$ and *Xbp1*$^{mKO}$ mice and cultured for 72 h, followed by immunostaining with anti-Pax7, anti-MyoD and DAPI to identify self-renewing (Pax7$^{+}$/MyoD$^{-}$), proliferating (Pax7$^{+}$/MyoD$^{+}$) and differentiating (Pax7$^{-}$/MyoD$^{+}$) satellite cells. **(A)** Representative images immunostained myofibers. Scale bar, 50 µm. Quantification of **(B)** number of clusters per myofiber, **(C)** average number of cells per cluster, and proportion of **(D)** self-renewing, **(E)** proliferating, and **(F)** differentiating satellite cells per myofiber of *Xbp1*$^{fl/fl}$ and *Xbp1*$^{mKO}$ mice. n=15-20 myofibers per group. All data are presented as mean ± SEM. *$p \leq 0.05$; values significantly different from myofibers of *Xbp1*$^{fl/fl}$ mice analyzed by unpaired Student *t* test.

**FIGURE S13**

FIG. 3H

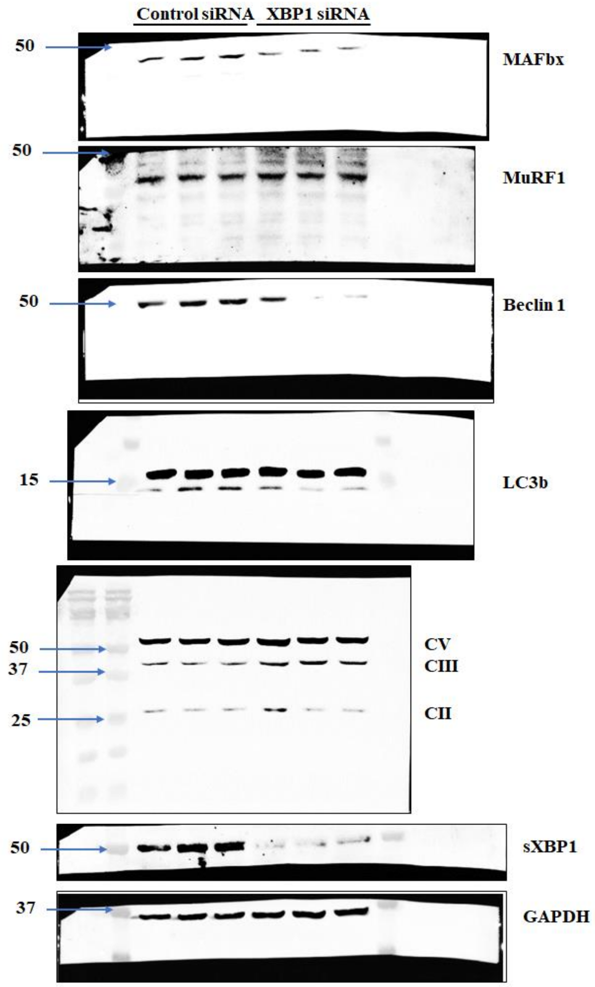


FIG. 5D

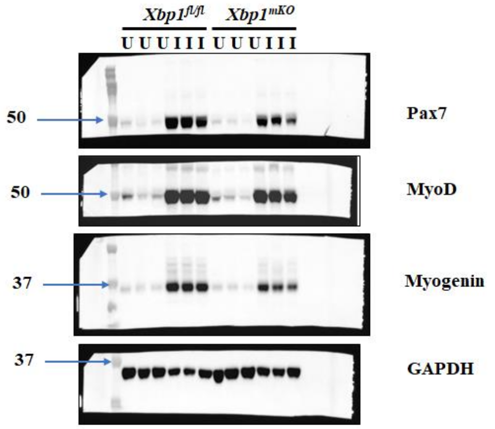


FIG. S8A

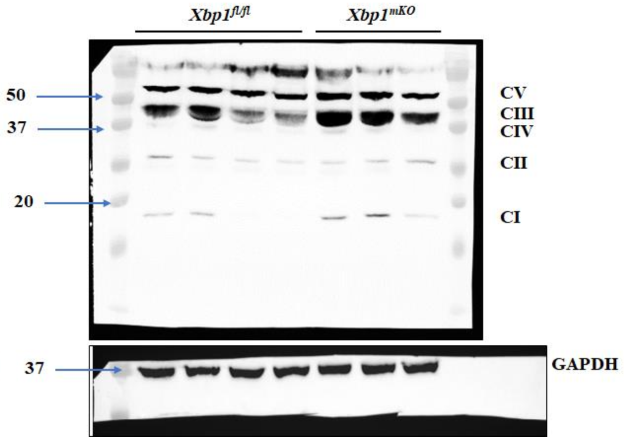


FIG. S8D

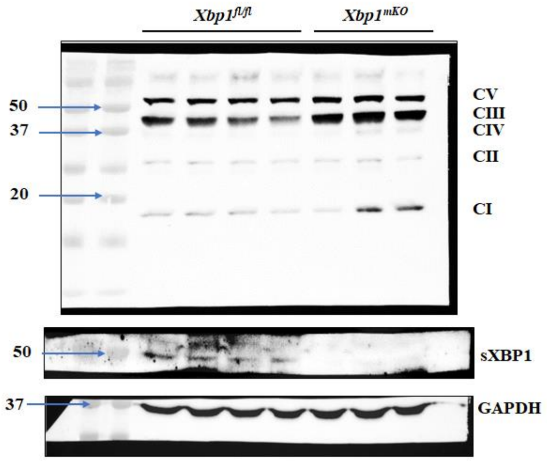


FIG. S8G

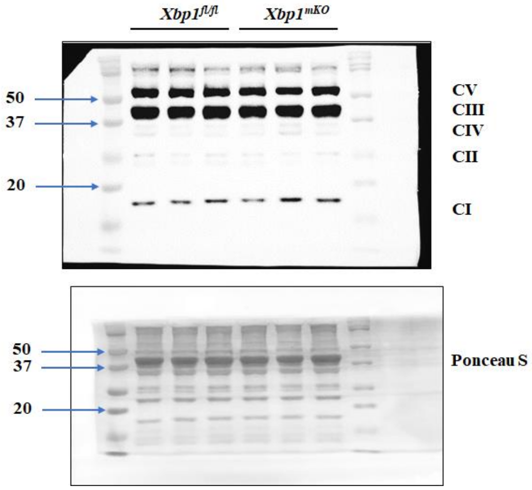


**Figure S13. Uncropped Western blot images.** Original immunoblots generated in the present study.
